# Photoswitchable microtubule inhibitors enabling robust, GFP-orthogonal optical control over the tubulin cytoskeleton

**DOI:** 10.1101/716233

**Authors:** Li Gao, Yvonne Kraus, Maximilian Wranik, Tobias Weinert, Stefanie D. Pritzl, Joyce Meiring, Rebekkah Bingham, Natacha Olieric, Anna Akhmanova, Theobald Lohmüller, Michel O. Steinmetz, Oliver Thorn-Seshold

**Affiliations:** Department of Pharmacy, Ludwig-Maximilians University, Butenandtstrasse 5-13, Munich 81377, Germany; Paul Scherrer Institut, Forschungsstrasse 111, CH-5232 Villigen PSI, Switzerland; Chair for Photonics and Optoelectronics, Nano-Institute Munich, Department of Physics, Ludwig-Maximilians University, Königinstrasse 10, 80539 Munich, Germany; Nanosystems Initiative Munich and Center for Nanoscience, Schellingstraße 4, 80799 Munich, Germany; Cell Biology, Department of Biology, Faculty of Science, Utrecht University, Padualaan 8, 3584 Utrecht, the Netherlands; Biozentrum, University of Basel, Klingelbergstrasse 70, CH-4056 Basel, Switzerland

**Keywords:** tubulin polymerisation inhibitor, microtubule dynamics, photopharmacology, colchicine, cytoskeleton

## Abstract

Here we present GFP-orthogonal optically controlled reagents for reliable and repetitive in cellulo modulation of microtubule dynamics and its dependent processes. Optically controlled reagents (“photopharmaceuticals”) have developed into powerful tools for high-spatiotemporal-precision control of endogenous biology, with numerous applications in neuroscience, embryology, and cytoskeleton research. However, the restricted chemical domain of photopharmaceutical scaffolds has constrained their properties and range of applications. Styrylbenzothiazoles are an as-yet unexplored scaffold for photopharmaceuticals, which we now rationally design to feature potent photocontrol, switching microtubule cytoskeleton function off and on according to illumination conditions. We show more broadly that this scaffold is exceptionally chemically and biochemically robust as well as substituent-tolerant, and offers particular advantages for intracellular biology through a range of desirable photopharmaceutical and drug-like properties not accessible to the current classes of photoswitches. We expect that these reagents will find powerful applications enabling robust, high precision, optically controlled cell biological experimentation in cytoskeleton research and beyond.

## Introduction

Molecular photoswitches have been used to install optical control over a broad range of phenomena, with applications from material sciences^1,2^ through to reversible photocontrol of ligand binding affinities^3^ and manipulation of diverse cellular processes in chemical biology^4,5^. For studies of temporally-regulated and spatially anisotropic biological systems, particularly those that simultaneously support several cellular functions, photoswitchable inhibitors conceptually enable a range of powerful studies not accessible with other tool systems.^6-8^

A prime example of such spatiotemporally regulated, multifunctional systems is the microtubule (MT) cytoskeleton. This complex cellular network plays central roles in nearly all directed mechanical processes, such as intracellular transport and cell motility; its crucial function in cell proliferation has also made it a central anti-cancer drug target.^9-11^ Yet, whereas cytoskeleton research typically aims to study a subset of MT-dependent processes that are spatially and/or temporally localised, nearly all MT inhibitors reported as tool compounds for biological research are drugs that are active wherever they are distributed, including at sites and at times where drug activity is not desired.^12^ This restricts the scope of applications and utility of these inhibitors for selective research into the various, highly dynamic, anisotropic processes dependent on MTs.^13^

The structure of the colchicine site MT inhibitor combretastatin A-4 (**CA4**; Fig 1a)^14^ has recently inspired photoswitchable solutions to the problem of achieving spatiotemporal control over MT inhibition. **CA4** is a stilbene whose *Z*-isomer (*cis*) binds tubulin, acts as a low nanomolar cytotoxin *in cellulo*, and reached Phase III trials as an anticancer drug.^15,16^ Crucially, its *E*-isomer (*trans*) is several orders of magnitude less bioactive.^17^ An approach to microtubule photocontrol *via* photoisomerisation of its bridging C=C bond has been proposed, whereby bioactive *Z*-combretastatins should be generated *in situ* from inactive *E-*precursors, allowing spatially and temporally precise application of antimitotic bioactivity.^18^ However, to the best of our knowledge this concept has not been realised in biology, hindered by the bio-incompatible short-wavelength illumination required (λ_max_ ∼ 300 nm) and the irreversible photochemical degradations that stilbenes undergo in aerobic conditions (e.g. oxidative 6π-electrocyclisation).^19^

**Figure 1.**
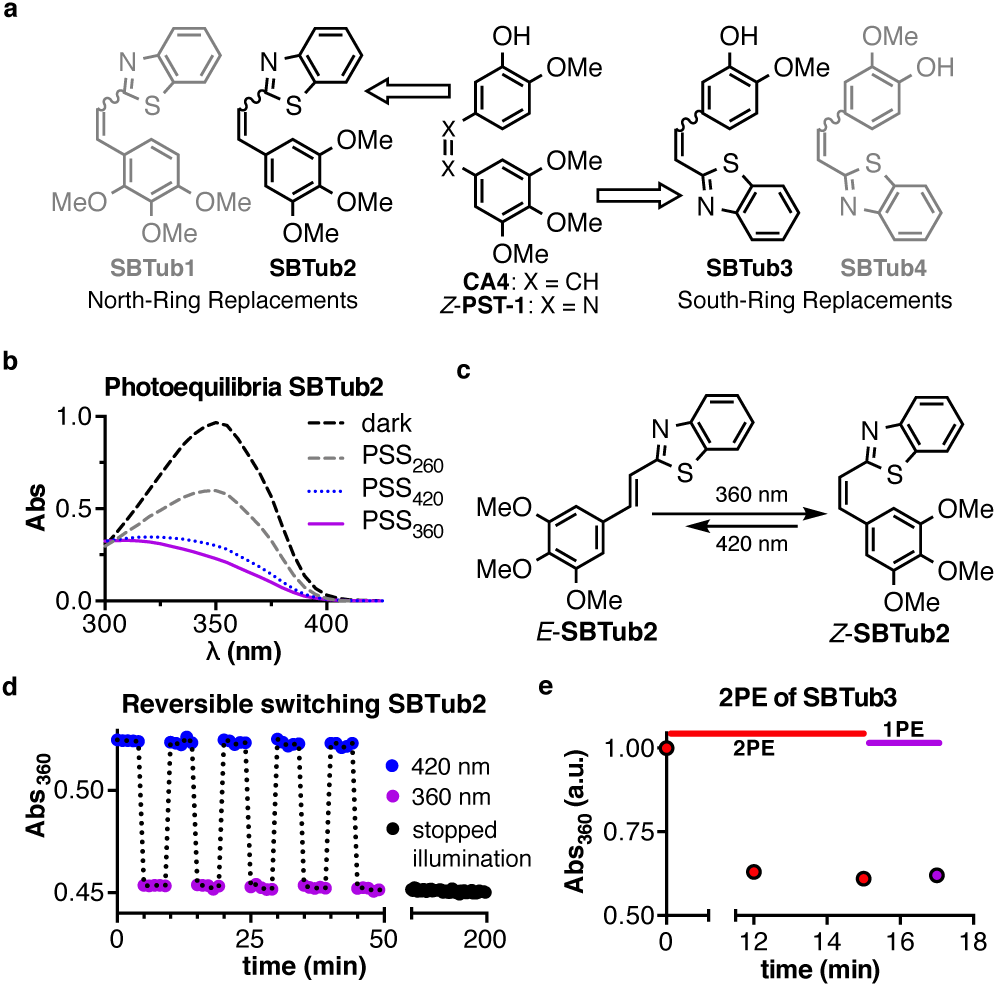
**a** Styrylbenzothiazole **SBTub**s based on the **CA4** pharmacophore. **b SBTub2** has illuminated PSSs showing a strong majority of the *Z*-form**. c** and **d SBTub**s can be switched from all-*E* to majority-*Z* using 360 nm, partially switched back with 260 or 420 nm, and they persist at PSS without appreciable thermal relaxation for hours when illumination is stopped. **e** Photoisomerisation of **SBTub3** *via* 2-photon excitation (2PE). (**b**: 25 µM and **d**: 50 µM **SBTub2**; in PBS, pH 7.4, 10% DMSO, room temperature, under closed air atmosphere; **e**: 5 mM **SBTub3**, 100% DMSO).

Instead, designing **CA4** analogues incorporating the synthetic azobenzene photoswitch scaffold inside their pharmacophore – a design strategy known as “azologisation”^20^ – delivered the biocompatibly-switchable, azoben-zene-based Photostatins (**PST**s; Fig 1a)^6,21-23^, which undergo optically reversible, bidirectional photoswitching between inactive *E* and MT-inhibiting *Z* isomers using low-intensity visible light. The **PST**s have enabled specific optical control over MT structure and dynamics in live cells and animals^6^ and have resolved key biological questions in mammalian development and neuroscience. These applications illustrate the power of photopharmacology to enable previously inaccessible studies of spatiotemporally anisotropic processes without genetic engineering, which is particularly important for studies of cytoskeletal scaffold proteins, since no optogenetic methods have yet been developed that can control their structure-dependent functions.^7,8,24,25^

However, azobenzene-based approaches to photocontrol of intracellular biology incur serious general disadvantages, which are well illustrated by the example of the **PST**s. Firstly, *in cellulo*, **PST**s were shown to be ∼2 orders of magnitude less potent than **CA4**. In general, we are also unaware of any azobenzene-based photopharmaceuticals that feature strongly photoswitchable bioactivity against cytoplasmic intracellular targets *in cellulo* (i.e. the diazene is intact, to provide photoswitchability), yet do not exhibit substantial loss of potency compared to their parent structures. Potency loss has been presumed^22^ to result from sequestration/degradation initiated by glutathione (GSH) addition to the N=N double bond once the azobenzene enters the cytoplasm ([GSH] ∼ 5-10 mM); this GSH susceptibility is known to affect the majority of azobenzenes^26,27^ and can cause rapid destruction particularly of electron-rich systems.^28^ This loss of potency, and the potential for uncharacterised degradative metabolites, obstruct robust applications of “azologues” especially *in vivo* (high doses required, degradation byproducts have to be evaluated) and this is expected to be a general phenomenon for azobenzenes applied to intracellular protein targets.^22,29^ Secondly, the *Z-to-E* spontaneous relaxation halflives of the water-soluble **PST** derivatives appropriate for *in vivo* work were restricted to the range of 1 – 60 min, requiring re-illumination of samples during most studies. This especially disfavored long-term applications: we have observed re-illuminations to cause photobleaching of fluorescent labels on tubulin^6^ as well as nonspecific photodamage in cell biology. Also, long-term spatially precise re-illuminations are problematic to deliver in (motile) embryos and animals. Thirdly, azobenzenes such as **PST**s are strongly photoisomerised by excitation wavelengths of typical imaging labels used in biological studies, being rapidly switched to ∼3:1 *E*:*Z* under 488 nm (GFP/fluorescein imaging) and to ∼4:1 *E*:*Z* under 514 nm (YFP/Cy3). Since the spatiotemporally-resolved application of the photoswitch is therefore prevented by imaging under these conditions, its practical potential to fulfil research uses both in cell biology and particularly in transgenic animals (where the most common imaging tag is GFP) is limited.^30^

We perceive that these three problems intrinsically associated to the azobenzene’s N=N double bond (non-orthogonality to standard imaging conditions, cellular/metabolic liabilities in intracellular settings, and substituent-dependent limits on thermal halflife scope) have largely prevented the translation of azobenzene photopharmaceuticals for intracellular targets from cell-free / cell culture settings, into more rewarding yet more complex *in vivo* research applications for controlling biological model systems up to higher animals. We were therefore motivated to develop new photoswitchable scaffolds avoiding these three problems – both for the specific case of photoswitchable MT inhibitors, as well as for general applications to chemical biology – and to examine their performance features that could suit them to more robust, orthogonal, biological photocontrol in complex systems.

To this purpose, we here explored styrylbenzothiazoles (SBTs) as alternative photoswitchable scaffolds, and evaluated them for chemical biology use by creating SBT-based MT-inhibiting photopharmaceuticals whose applications we demonstrate especially in live cell studies with *in situ* photoswitching. We expected that, as compared to azobenzenes, the central C=C double bond of styrylbenzothiazoles^31,32^ would endow them with increased robustness towards GSH reduction and higher thermal stability of the *Z* state, while avoiding photoresponse to wavelengths above 460 nm (due to the absence of an n-π* band)^33^; yet that, compared to stilbenes, the benzothiazole would redshift the π-π* band enough to be photoswitched with visible light^34^ while also blocking photochemical degradation.^35^ Accordingly, we synthesise a rationally designed series of **SBTub**s (**SBT**-based t**ub**ulin-inhibitors) and evaluate them as biochemically robust, GFP-orthogonal, photoswitchable analogues of **CA4**. We obtain ligand-protein crystal structures confirming their two modes of tubulin binding, then demonstrate their optical control over MT network integrity, cell division and cell death, which is to our knowledge the first use of styrylbenzothiazoles as photopharmaceuticals in chemical biology. Finally, using their unique photoresponse characteristics, we demonstrate that they enable temporally reversible, optical modulation of MT dynamics independently of ongoing imaging, in live GFP-tagged cells.

## Results

### Design and Synthesis

We designed two orientations of **SBTub**s as closely isosteric replacements of the MT inhibitor **CA4** in the expectation that these **SBTub**s would display *Z*-isomer-specific MT-disrupting effects. These became **SBTub2** (where the benzothiazole ring replaces the “north,” isovanillinyl ring of **CA4**), and **SBTub3** (where the benzothiazole replaces the “south,” trimethoxyphenyl ring; Fig 1a). Since we expected that the benzothiazole would give similar space occupancy, geometry and polarity intermediate as the replaced rings, and since the colchicine site is known to be somewhat accommodating^16,36^ particularly of north ring bicyclic aromatics, we expected that both these *Z-***SBTub**s would bind satisfactorily despite having sacrificed some potency-enhancing point interactions. *E-***SBTub2** has in fact been reported as a cytotoxic resveratrol analogue targeting tubulin, where its *E*-geometry was established explicitly and extensive docking simulations were performed to rationalise observed activity^37,38^. However, the *E-*activity stated in those reports disagrees with our experience^6,39,40^ and literature understanding^16^ of the requirements of the colchicine binding site. We decided therefore to pursue our paper’s design logic paying particular attention to experimentally verify the isomer status and binding mode of the compound (discussion in Supporting Information).

In order to test the target-specificity of the **SBTub**s’ bioactivities, we also designed permutation controls (designed-inactive compounds) by swapping the positions of key bioactivity-controlling substituents. We have previously reported permutation controls as a convenient and stringent method^39^ to distinguish between biological disruption resulting from molecularly-specific binding to the target protein (which should be a feature of the designed-active compounds only), versus bioactivity from nonspecific interactions expected for typically hydrophobic photoswitch compounds (e.g., promiscuous binding to proteins, aggregation on proteins, compound precipitation, membrane disruption, phototoxicity, or photoswitch scaffold toxicity, which should also be observed for closely regioisomeric permutations). To this end, we permuted methoxy and hydro groups of **SBTub2** to create designed-inactive **SBTub1**, and permuted hydroxy and methoxy groups of **SBTub3** to create control **SBTub4**. *E*-**SBTub4** has also previously been synthesised in the context of ligands for platinum complexes^41^, although its isomerisation to *Z* was not considered; here we require it specifically as a *Z-*isomeric control and accordingly we pursued its synthesis and characterisation (discussion in Supporting Information).

*E-***SBTub1**–**4** were synthesised in good yields by aldol condensations of 2-methylbenzothiazole with the corresponding aldehydes in the presence of a strong base (Fig 1a and Supporting Information).

### Photoswitch performance

The spectral characteristics of the **SBTub**s were similar (Fig S1). The shift of π→π* absorption maxima between the *E*-(∼360 nm) and the *Z*-isomers (∼305 nm) enabled directional photoisomerisations. Pleasingly, both isomers’ absorptions decreased sharply towards zero above 410 nm. This is important for avoiding photoisomerisation under GFP imaging (488 nm), since we and others have found that small absorption “tails” far beyond absorbance maxima can cause substantial photoswitching in microscopy experiments (e.g. in the azobenzene-based **PST**s, the 561 nm laser line can photoisomerise a *Z*-isomer with absorbance maximum 445 nm). Our expectation was therefore, that the **SBTub**s’ absorption cutoff occurs at the optimum position to avoid *E*↔*Z* photoisomerisations under 488 nm (“GFP-orthogonality”), while still permitting their photoswitching in standard cell biology experiments by using the most common short-wavelength microscopy laser line, 405 nm.

Under physiological conditions, the **SBTub**s could be optimally *E*→*Z* photoisomerised with 360 nm light, giving a photostationary state equilibrium (PSS) with *Z:E* ratio of ∼85:15 (Fig S6); applying shorter or longer wavelengths back-isomerised the **SBTub**s towards more *E-*enriched PSSs (Fig 1b-1c). We selected “dark” (all-*E*) and “lit” (360 nm, mostly-*Z*; or 420 nm, mixed *E* and *Z*) illumination conditions for further use in long-term cell culture.^42^ Pleasingly for our design aims, reversible photoisomerisations with high-power illuminations repeatedly traversing the biologically applicable range 360 – 420 nm showed no signs of causing photodegradation, under aerobic aqueous conditions (Fig 1d).

The **SBTub** spectra and photoswitching properties were entirely robust to variations of pH and solvent (Fig S2). The metastable *Z*-isomers of all **SBTub**s could be quantitatively relaxed to *E* by warming to 50-60°C in DMSO overnight, although at 25°C they showed no significant thermal relaxation within hours at pH∼7 in aqueous media (Fig 1d). However, *para-*hydroxy **SBTub4** thermally relaxed at pH∼9 (Fig S3). We interpret its pH-dependent relaxation resulting from resonance between the deprotonated phenolate (where the bridging bond is a C=C double bond) and a quinoidal form (bridging C-C single bond, whose free rotation to the thermodynamically more stable *E* conformation might occur quickly). This offers two opportunities for future applications of styrylbenzothiazole photopharmaceuticals. Firstly, the *Z-*stability of *para-*hydroxy **SBTub4**’s *Z* isomer at pH∼7 contrasts to both azobenzene and hemithioindigo photoswitches with *para*-(or *ortho*-) hydroxy or aniline substituents, that typically feature millisecond (azobenzene) to second (hemithioindigo) aqueous relaxation halflives even at pH∼7. This underlines the broader chemocompatibility of styrylbenzothiazoles as a photoswitch scaffold for these functional groups, that are highly prevalent in bioactive molecules for many biological targets due to e.g. their importance in creating high-affinity ligand-protein interactions. Secondly, the pH-sensitivity of *para*-hydroxy **SBTub4**’s relaxation rate suggests that pK_a_-modulation could deliver styrylbenzothiazoles relaxing on a second or minute timescale at pH∼7, such that local *E*→*Z* photoswitching could be combined with sample-wide thermal relaxation to improve spatiotemporal localisation of the *Z-*isomer.

Finally, we investigated the possibility of two-photon absorption/excitation (2PA/2PE) to photoisomerise **SBTub**s, which would promise increased spatial resolution of isomerisation compared to single-photon processes.^43^ This may be especially beneficial in cell-free studies requiring high spatial resolution, such as time-resolved spectroscopic studies of tubulin structural rearrangements following ligand isomerisation inside a lattice. We used a mode-locked Ti-Sapphire laser at 780 nm to bulk photoisomerise **SBTub3** in concentrated (5 mM) DMSO solution, delivering 2PA/2PE inside a single voxel and relying on diffusion to reach *Z/E-*isomer equilibrium in the whole 8 µL sample. After 12 min of 2PA/2PE the isomer equilibrium did not evolve with additional 2PA/2PE illumination or with single-photon 360 nm illumination (Fig 1e, Fig S7), indicating that ∼85% *Z* was reached by 2PA/2PE alone. The **SBTub** scaffold can therefore be *E* to *Z* photoswitched *via* efficient, spatially precise 2PA/2PE isomerisation.

### (Bio)chemical and photochemical stability

We next wished to assess key aspects of the **SBTub** scaffold’s photochemical and biochemical robustness. We assayed the stability of both the *E* and *Z*-**SBTub**s to high-photon-flux irradiation (Fig S4-5), observing them to be entirely photostable to >1 h continuous high-intensity UV irradiation (Rayonet photoreactor), underlining their outstanding photochemical stability. We also assayed their sensitivity to GSH addition (10 mM) (Fig 2a, Fig S8), monitoring changes due to thiol adduct formation, thiol addition-elimination (that should return *E*-**SBTub** due to the sp^3^ intermediate), reductive degradation (involving a second equivalent of GSH) or other compound loss mechanisms, yet no such changes were observed. This contrasts markedly to the instability in our hands of similar *Z-*azobenzenes when challenged by GSH, including **PST-1** (note however that our GSH challenge assay, which avoided the biologically irrelevant exogenous phosphine nucleophile TCEP, resulted in substantially less adduct formation/photoswitch destruction than literature methods^22^; Fig S8-9). We also performed an early *in vitro* metabolic assessment of **SBTub2** and **SBTub3** for their suitability as *in vivo* drugs, examining stability to processing by liver microsomes (Fig 2b), inhibition of representative cytochromes (Fig S11), and hERG channel inhibition (Fig S12). Especially for **SBTub3** these assays did not indicate typical drug development problems. We are unaware of such a demonstration for other photoswitch scaffolds.

**Figure 2.**
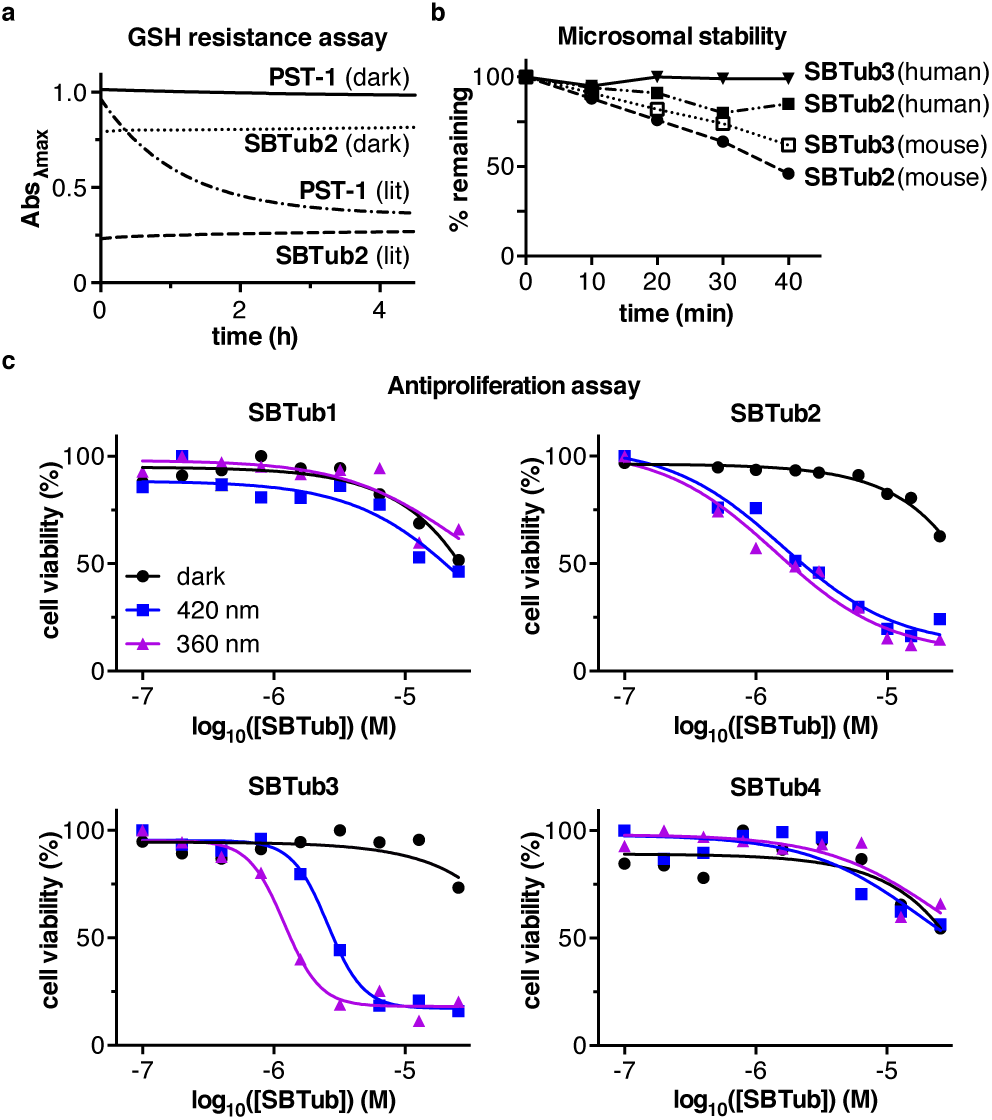
**a** Glutathione (GSH) resistance assay, monitoring change of photoswitch absorbance during incubation with 10 mM GSH. Representative styrylbenzothiazole **SBTub2** (25 µM) and representative azobenzene **PST-1** (50 µM dark, 300 µM lit) under dark or lit conditions (**PST**: 395 nm, **SBT**: 360 nm) were monitored at their π→π* band maximum (except lit **PST-1**, where the n→π* band was followed). **b SBTub2** and **SBTub3**’s resistance to human and mouse liver microsomes. **c** Antiproliferation assay of **SBTub**s shows light dependent cell toxicity for **SBTub2** and **SBTub3**. HeLa cells, 40 h incubation either in dark conditions (all-*E*), or under pulsed illuminations (75 ms per 15 s) with 360 nm or 420 nm LEDs (<1 mW/cm^2^). One representative experiment of three independent experiments shown.

Taken together, we consider these features of styrylbenzothiazoles’ performance as a photoswitch scaffold, as well as their differentiating features compared to azobenzenes, very promising for further biological applications towards *in vivo* use. We thus began exploring the applicability of **SBTub**s as photoswitches for MT control, and more generally as a new scaffold for photopharmaceutical use in cell biology research.

### SBTubs photoswitchably inhibit microtubules *in cellulo*

Since prolonged inhibition of cellular MT functions results in block of cell proliferation and ultimately in cell death, we first assayed the light-dependent antiproliferative activity of the **SBTub**s by the MTT cell viability assay (Fig 2c) on HeLa cervical cancer cell line (incubation either in the dark, or under pulsed illuminations with 360 nm or 420 nm light). Both **SBTub2** and **SBTub3** were photoswitchably cytotoxic with ≫20-fold cytotoxicity enhancement under 360 nm illumination conditions (IC_50_ ∼1-2 µM with ∼85% *Z* isomer) as compared to the dark experiments (IC_50_ ≫ 20 µM with ∼100% *E* isomer), highlighting the suitability of this photoswitch scaffold for long-term intracellular photopharmacology. Neither of the permutation controls **SBTub1**/**SBTub4** displayed significant or photoswitchable bioactivity. We attributed the weak antiproliferative activity of all **SBTub**s at high concentrations in the dark, and of permutation controls **SBTub1** and **SBTub4** also under illuminated conditions, as aggregation-dependent effects resulting from poor solubility of the compounds at high concentrations as is known for similar motifs (this is examined in more detail below).^6,39^ At this stage, the light-dependent *in cellulo* activity matching the design principles of the active **SBTub**s, as well as the lack of obvious toxicity for their regioisomeric permutation controls, supported the assumption that **SBTub2** and **SBTub3**’s cellular antiproliferative potency is based primarily on MT inhibition by the *in situ-*photogenerated *Z-*isomers only.

To examine the molecular mechanism of light-dependent cellular bioactivity, we solved the structure of *Z*-**SBTub2** and *Z*-**SBTub3** in complex with the tubulin-DARPin D1 complex^44^ to 2.05 and 1.86 Å-resolution by X-ray crystallography, respectively (Table S1). Both active *Z*-**SBTub**s bind to the colchicine site (located at the interface of α- and β-tubulin; Fig 3a, Fig S17-S22), and as designed, they bind in opposite poses despite contacting identical residues in the binding pocket. *Z-***SBTub2** adopts the “top-down” pose characteristic of trimethoxy-phenyl-bearing **CA4** analogues^16^, whereas *Z-***SBTub3** binds in the opposite “bottom-up” pose (Fig S20). The benzothiazole can therefore occupy the same space as either the trimethoxyphenyl or the isovanillyl unit of typical colchicine-site inhibitors,^40^ giving an intriguing top-vs-bottom desymmetrisation of the photoswitch. This should also allow rational design and tuning of **SBTub**-based photopharmaceuticals that project substituents such as functionally diverse side-chains and reporters outwards from the colchicine binding site.^39^

**Figure 3.**
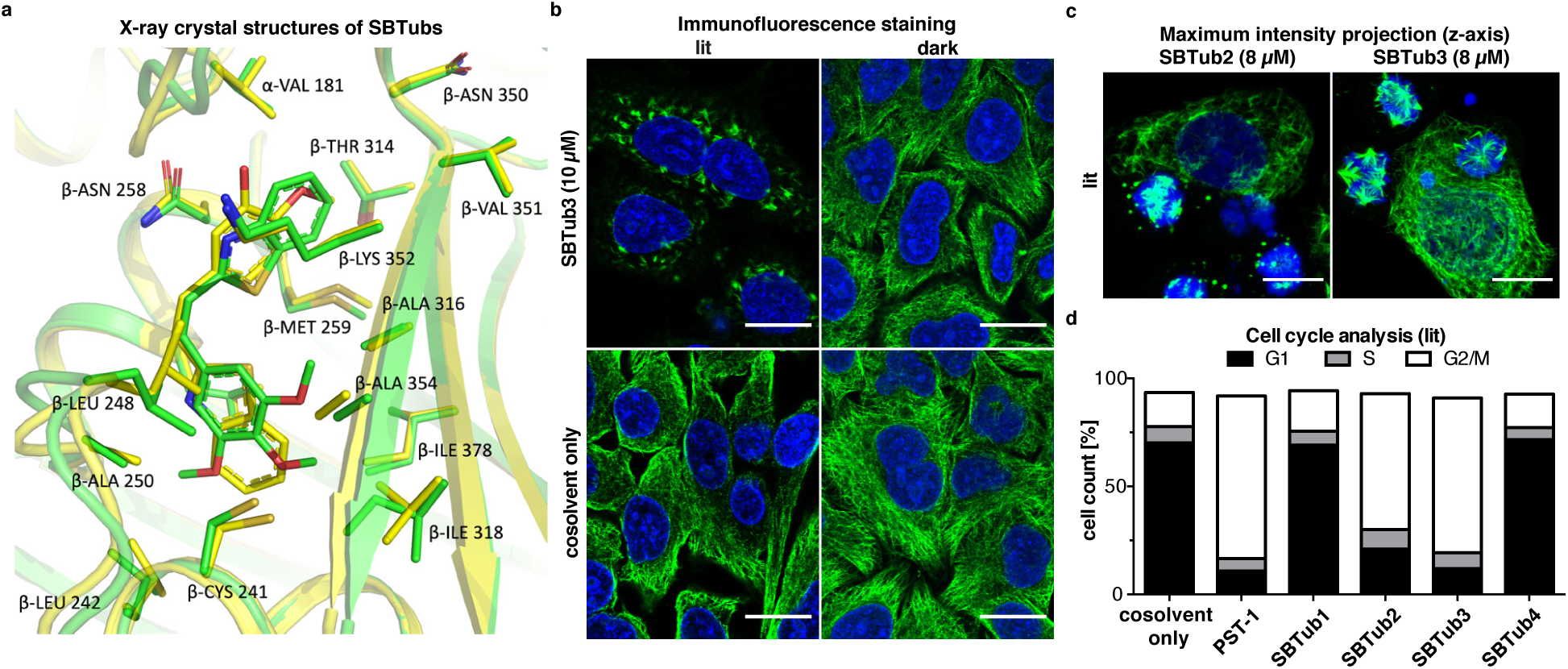
**a** Superimposition of structures showing the colchicine site of tubulin in complex with **SBTub2** and with **SBTub3**. Tubulin is shown in cartoon representation; interacting residues are shown in stick representation and are labeled. Carbons are coloured green in the **SBTub2**:tubulin structure, and yellow in the **SBTub3**:tubulin structure, with oxygens and nitrogens coloured red and blue, respectively. **b SBTub3** dose-dependently induces MT breakdown under 360 nm pulsing (“lit”), while MT structure remains unaffected in the dark (see also Fig S14). **c** Maximum intensity projections along the z-axis showing light-specific mitotic arrests. (**b & c**: HeLa cells incubated for 20 h; α-tubulin in green, DNA stained with DAPI in blue, scale bars 20 µm). **d** Cell cycle analysis of **SBTub1**-**4** (20 µM) alongside positive control **PST-1** (2.5 µM) shows significant G_2_/M arrest for **SBTub2**, **SBTub3** and **PST-1** under 360 nm pulsing (see also Fig S13).

We next assayed the potency of the **SBTub**s as light-dependent tubulin polymerisation inhibitors in cell-free assays using purified tubulin protein. **SBTub**s showed no inhibition of tubulin polymerisation under light-excluded conditions (∼100% *E*) but strong dose-dependent inhibition after illumination (Fig S10) clarifying their mechanism of action as tubulin polymerisation inhibitors (also known as microtubule depolymerisers), as is expected for colchicine site ligands.

We then began examining the cellular mechanism of **SBTub** bioactivity, by performing immunofluorescence imaging of the endogenous MT network structure in cells incubated with *E/Z*-**SBTub**s. **SBTub2** and **SBTub3** caused no effects on MT network structure as the *E-*isomers, but dose-dependently caused mitotic arrest and MT depolymerisation under 360 nm pulsed illumination with similar dose-dependency as in the viability assays (Fig 3b; Fig S14). Z-stack images revealed cells not clearly visible to single focal plane IF imaging, which were substantially accumulated into rounded, mitotically arrested states (Fig 3c). These are hallmarks of treatment with MT-inhibiting antimitotics, supporting the conjecture that **SBTub2** and **SBTub3**’s strongly photoswitchable cytotoxicity *in cellulo* arises from the *Z-*isomer inhibiting MT function. By contrast, the permutation controls **SBTub1** and **SBTub4** showed no disruption of MT network structure under *either* illumination condition even at high concentrations (Fig S14c), further supporting the MT-specific binding activity of *Z*-**SBTub2-3**.

If their major mechanism of bioactivity *in cellulo* is direct inhibition of MT function, *Z-***SBTs** should induce light-dependent G_2_/M-phase cell cycle arrest,^16^ and we tested this using flow cytometric analysis of **SBTub**-treated cells. We observed G_2_/M-arrest for both *Z-***SBTub2** and *Z-***SBTub3** (Fig 3d): but not for their *E*-isomers, nor for either isomer of the permutation controls **SBTub1** and **SBTub4**, even at the highest concentrations tested (Fig S13a). This supports the notion that the *E-***SBTub**s have negligible *specific* MT-inhibiting effects while the appropriately substituted *Z* isomers potently and structure-specifically inhibit MT function, suiting both **SBTub2** and **SBTub3** to photoswitching-based optical control of MT-dependent processes.

With their photoswitchable cellular bioactivity now characterised, we then set out to demonstrate some of the conceptual advantages that the photophysical and biochemical properties of the SBT scaffold endow upon the **SBTub**s, when compared to azobenzenes and other major photoswitch types and when considering their most likely domains of biological application.

### Bulk isomerisation of SBTubs enables straightforward long-term MT photocontrol

Firstly, noting the stability of their *Z*-isomers to biochemical challenge as well as to spontaneous relaxation under *in vitro* conditions (Fig 2a-b), we were also interested to study whether a single *E*→*Z* isomerisation event could be used to apply long-lasting bioactivity in cells without requiring re-illuminations throughout the assay (as is necessary with azo-benzene-based **PST**s that have relatively short relaxation halflives).^6^ We accordingly applied *E*-**SBTub3** to cell culture, isomerised it *in situ* to a majority *Z* population in the whole well using 18 seconds of low-power LED illumination at 360 nm, then shielded the cells from all outside light sources and observed cell viability 40 hours later. The *Z*-**SBTub**s’ antiproliferative potency in this single-shot experiment (Fig 4a) nearly matched that observed with pulsed re-illuminations, indicating the stability of the *Z*-**SBTubs** *in cellulo* in the long term (despite the presence of e.g. cellular thiols). This result should be compared with azobenzene-based **PST**s which were essentially inactive in the single-shot experiment, since reversible thiol addition-elimination and/or degradative metabolism (see Fig 2a) as well as thermal relaxation deplete *Z*-**PST** levels rapidly (literature suggests within an hour^22^).

**Figure 4.**
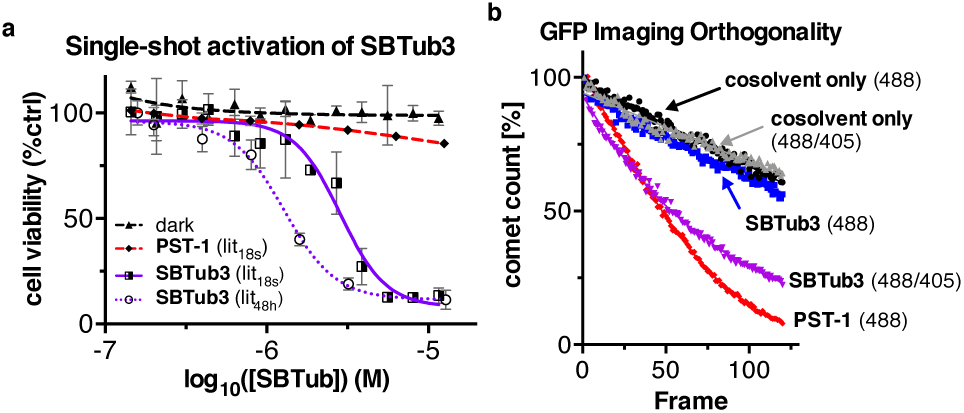
**a** Comparison of cellular antiproliferative activity assays of **SBTub3** and **PST-1** under “single-shot photoactivation” (lit_18s_: 75 ms pulses of ∼1 mW/cm^2^ LED illumination per 15 s during a 1 h period) and ongoing illumination (lit_48h_: same pulsed illuminations, continued over 48 h) protocols, shows the **SBTub**’s sustained pharmacological activity after a single photoactivation period, differentiating it from the cognate azobenzene (HeLa cells, 48 h). **b** Disappearance of EB3-GFP “comets” (due to nonspecific photobleaching and/or specific inhibition of tubulin polymerisation) during live cell imaging under 488 nm. 488 nm-only imaging in the presence of **SBTub3** (10 µM) reproduces the comet depletion observed with cosolvent-only controls (native bleaching dynamics with 488 nm or alternating 488/405 nm pulses) indicating no *E*→*Z* photoconversion under 488 nm laser (i.e. *E-***SBTub** is GFP-orthogonal). With alternating 488/405 nm illuminations, **SBTub3** induces a rapid shutdown of tubulin polymerisation dynamics similarly to **PST-1** under 488 nm illumination only (i.e. the azobenzene **PST-1** cannot be used GFP-orthogonally).

### SBTubs are photocontrollable orthogonally to GFP/YFP imaging

We next performed a conceptually similar experiment with single cell imaging of microtubule dynamics and showed that bulk application of activated *Z-***SBTub** maintains a blockade of MT dynamics without significant evolution over the experimental timecourse even under 488 nm laser imaging (Fig S15). This encouraged us to more rigorously evaluate **SBTub**s for their ability to effect *in situ* photocontrol over MT dynamics while avoiding interference from imaging of the most common fluorescent labels GFP and YFP, which would allow fully orthogonal imaging and photocontrol in live systems. This is a particularly important unsolved challenge for intracellular photopharmacology, which typically relies on imaging-based readouts, since conventional azobenzene^4,6^ and hemithioindigo^39^ photopharmaceutical scaffolds almost always feature uni- or bidirectional photoresponse to these imaging wavelengths such that they would require complex and impractical light-compensation protocols to counteract isomerisation caused by monitoring the experimental readout. **SBTub** *E* to *Z* photoisomerisation by 405 nm rapidly halted normal MT polymerisation dynamics, but as expected from their sharp absorption cutoff above 410 nm, neither high-intensity GFP (488 nm excitation) nor YFP (514 nm) imaging lasers significantly impacted either the formed *Z-***SBTub** or, in a separate experiment, the starting *E-***SBTub** isomer (readout by EB3-GFP and EB3-YFP MT “comet” assays^6^, Fig 4b and Fig S16; discussion in the Supporting Information). As photocontrol over the SBT scaffold is fully orthogonal to GFP/YFP imaging, we consider SBTs may be particularly well-suited for photopharmaceutical applications in biological systems where GFP and YFP reporters are used.

### SBTubs can deliver temporally reversible, MT modulation *in cellulo* during GFP imaging

While the present **SBTub**s could be locally *E*→*Z* photoisomerised with high spatiotemporal precision, they were explicitly not designed for spontaneous *Z*→*E* reversion on biological timescales. However, we observed that addition of *Z*-**SBTub3** to cell bath solutions stopped cellular MT dynamics extremely quickly (Fig S16b), implying fast membrane penetration kinetics. This is an encouraging result, since achieving temporally reversible, spatially localised cellular applications of lit-active photopharmaceuticals^39^ – which brings the major advantages of photopharmacology to cytoskeleton research – conceptually only requires spatiotemporal localisation of the activating photoisomerisation, as long as diffusion across membranes can reduce the intracellular concentration of the active isomer below its inhibitory threshold. If inhibitor-target association is kinetically highly reversible and diffusion across the cell membrane is fast, this concentration change may occur on a biologically relevant timescale; as long as inhibitor dissociation immediately restores the protein’s function (as for colchicine domain inhibitors) – and particularly in systems with a highly nonlinear dose-response (such as the MT cytoskeleton) – such concentration changes may result in full biological recovery on this timescale. When photoactivation is restricted to a single cell, rather than the whole solution (as was done in Fig 4), then diffusion-based depletion of the the photogenerated bioactive isomer can allow cellular functional recovery on the compound’s transmembrane partitioning timescale.

We therefore designed experiments to test the temporal reversibility of **SBTub**’s cell-localised modulation of MT dynamics using another EB3-GFP “comet” assay, this time with cell-specific photoactivations. These allowed us to repeatedly pause and recover functional MT polymerisation during GFP imaging with statistics closely aligning over many cells (averaged data in Fig 5a; single cells in Movie M1 and Fig 5b), demonstrating that **SBTub**s can achieve highly reproducible, GFP-orthogonal, temporally reversible MT modulation in live cells.

**Figure 5.**
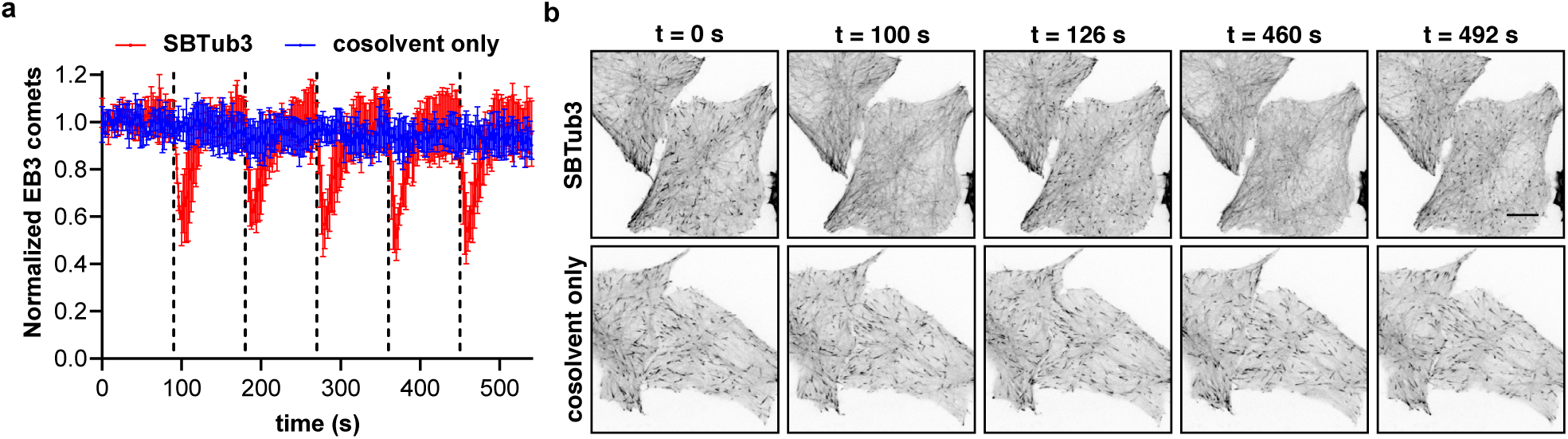
Data related to Movie M1: HeLa cells with or without 5 µM **SBTub3** were exposed to cell-localised single pulses of 405 nm every 90 s while imaging EB3-GFP comets at 491 nm, achieving temporally reversible MT modulation. **a** MT polymerisation dynamics, quantified by EB3 comet counts, are reversibly modulated by cell-localised photoactivations with **SBTub3** (red) but are unaffected in its absence (blue) (dotted lines indicate photoactivation pulses, n = 5 movies acquired per condition, data represented as mean comet count over time, normalised to 1 at t = 0 s, with standard deviation). **b** Representative still images of EB3 comets at t = 0 s (baseline), 10 s after the first activation (t = 100 s, comets vanished), 36 s after the first photoactivation (t = 126 s, comets recovered), 10 s after the fifth activation (t = 460 s, comets vanished), and 42 s after the fifth photoactivation (t = 492 s, comets recovered). Scale bar indicates 10 µm.

## Discussion and Outlook

Photopharmacological approaches to generate high-spatiotemporal-precision reagents for non-invasive studies of endogenous protein function have made significant progress over recent years. Their original applications to neuroscience^45,46^ have now been extended to studies of membrane^47,48^ and intracellular^6^ biology, and recent advances such as long-wavelength-responsive photoswitches^26,49,50^ offer beguiling prospects for *in vivo* photopharmacology. However, the conceptual scope of photopharmacology has been somewhat chemically restricted, in that the overwhelming majority of photopharmaceutical designs employ azobenzenes as the photoswitch moiety, and that independent of the desired biological target, the azobenzene motif incurs functional limitations on biological applications: (1) the diazene’s nucleophilic and metabolic susceptibility is problematic for addressing intracellular targets or when aiming at long-term, systemic *in vivo* applications; (2) its isomerisation-inducing n-π* transition in the blue-green spectral region overlaps with imaging of common biological tags (e.g. GFP/YFP) such that orthogonal photocontrol is not possible; (3) azobenzenes face restricted substituent scope for cytosolic applications as photopharmaceuticals, since if strong *ortho/para*-electron-donating groups (OH, NH_2_ etc) or push-pull substituent configurations are present, this causes µs relaxation rates meaning that the products are not bulk photoisomerisable under biocompatible conditions in aqueous media (especially problematic since these substituents are powerful point interaction groups for establishing ligand-target interactions and specificity). Only recently has the range of photoswitchable scaffolds incorporated into bioactive pharmacophores been expanded to include dithienylethenes^51^ and hemithioindigos,^39^ and the development of novel scaffolds with promising photoswitchability is recognised as a valuable milestone even before achieving cellular applications.^52^

Here we aim to usefully expand the toolbox of photoswitches for chemical biology, by demonstrating that styrylbenzothiazoles (SBTs) – a chemical scaffold with simplicity comparable to that of azobenzenes – can deliver spatiotemporally-precise, photoswitching-based control over critical cell biology processes, but with (1) added biochemical/metabolic robustness important for intracellular targets and *in vivo* application, (2) full orthogonality to common imaging conditions enabling easy adoptation of the photoswitch in existing biological systems, (3) alternative ranges of isomerisation-tolerant substituents (eg. *para*-OH) and relaxation rates. To this end, we have demonstrated the biological applicability of SBTs by rationally designing two alternative geometries of **SBTub**, each targeting tubulin with high specificity, highlighting the flexibility of this scaffold. We have shown in long-term cellular studies as well as in short-term live cell imaging that **SBTub**s can be used as a potent, rapidly-responsive, photochemically and biologically robust, GFP-orthogonal and cell-compatible tools for non-invasive photocontrol of endogenous MT dynamics and MT-dependent processes.

While these first incarnations of cell-targeted SBT-based photopharmaceuticals do not allow majority *Z*→*E* photoisomerisation in the near-UV or visible spectrum, we find that the photoswitching performance of the as-yet unexplored SBT scaffold merits further attention for biological applications. For example, we succeeded in demonstrating that **SBTub**s still enable temporally reversible live cell photocontrol studies, using standard microscopy illuminations. Diffusion-reliant reversibility can also be delivered by intracellular photouncaging, however, even in this protocol SBTs offer several biologically important performance advantages. Typical photocage groups (e.g. *ortho*-nitrobenzyl derivatives) feature a slow intermediate hydrolysis step that is rate-limiting for cargo release (up to the minute-scale), which drastically diminishes their precision of temporal reversibility in both photouncaging (off-to-on) and diffusion (on-to-off) processes; whereas the SBTs feature near-instantaneous photoresponse and reproducibly high-temporal-precision biological control. Furthermore, photouncaging byproducts are often panspecifically toxic and may also be phototoxic^53^, in stark contrast to **SBTub**’s clean and photochemically robust *E*→*Z* photoconversion which allows cell-tolerated long-term experimentation. In a practical sense for experimenters, accidentally light-exposed SBT stocks can be thermally relaxed quantitatively back to the *E* isomer by warming, in contrast to photouncaging probes that must be protected from irreversible ambient light-induced photodegradation during synthesis, storage and use. Lastly, we show that SBTs can also be straightforwardly integrated inside a bioactive scaffold avoiding penalties of compound complexity, in contrast to photouncaging strategies that increase synthetic cost and molecular weight, often penalising biodistribution.^53,54^ We consider these as promising features for SBT applications to high-precision, reproducible, “clean” phototriggered biological control.

By unifying photochemical and biochemical robustness, the SBT scaffold’s performance is particularly desirable for tissue slice or *in vivo* contexts. The excellent GSH resistance of SBTs as compared to azobenzenes (stability to intracellular environments) recommends them for addressing intracellular targets, and their microsomal stability may enable systemic applications in animals. Furthermore, **SBTub** bioactivity can be photolocalised in the long term via an experimentally straightforward single illumination. This avoids the requirement for repeated precisely-relocalised illuminations which can be experimentally problematic to apply in motile tissues of free-moving animals – and which more rapidly-relaxing photopharmaceutical systems such as azobenzenes (when the latter can survive long-term exposure to intracellular conditions) require. It also reduces the overall photon flux SBTs require for operation. The **SBTub**s’ violet-light operation and their extreme photochemical robustness under biological conditions also differentiates them from previously proposed but not yet *in situ*-demonstrated^55^ deep-UV-operated stilbene-based probes, which are likely to suffer electrocyclisation-oxidation degradation^56^.

Particularly in the context of *in vivo* microtubule research it should be pointed out that **CA4**-like MT inhibitors (which **SBTub**s mimic) are most often applied as vascular disrupting agents and studied using surface-illumination imaging by one- and two-photon microscopy^57^ – which we here demonstrate are feasible approaches to photoswitch the SBT scaffold. In these superficial tissue settings, the penetration of violet light required for **SBTub** isomerisation is not problematic; our ongoing results as well as published reports^58^ have shown successful subcutaneous isomerisation of photoswitches even with simple transdermal 360 nm LED illumination. Rather than light accessibility for tissues, we consider that two compound-specific factors are likely to determine the success of reaching *in vivo* applications with *in situ*-actuated photoswitches for intracellular targets. One is systemic pharmacokinetics (PK); while no reports of photopharmaceuticals’ PK have yet been published, metabolic stability and druglikeness – as the SBTs feature – are positive signs. The second is the ability to robustly photomodulate intracellular biology, which relies on the intracellular biochemical stability of the bioactive isomer, and in this respect SBTs show superior performance compared to the cognate azobenzenes. We therefore consider the **SBTub**s to offer exciting potential for *in vivo* translation within the context of MT photopharmacology.

Lastly, the design logic of scaffold hopping from azobenzene to SBT is in our opinion an exciting advance for photopharmaceutical design. Progress within photopharmacology has recently been stimulated by general strategies to reshape aspects of azobenzene performance without requiring pharmacological redesign; and in this vein, diazocines^59,60^, tetra-*ortho*-substitutions^50^ and azoniums^27^ are all garnering rapid and widespread applications. The rational design and biological success of the present work similarly suggests that much of the existing azobenzene-based photopharmacology can be straightforwardly grafted onto the new and potentially more robust SBT scaffold when it is desirable to improve performance on intracellular targets or *in vivo*, or to achieve more straightforward optically orthogonal applications to live-imaging-based biological studies.

In closing, we predict that the **SBTub**s themselves will contribute greatly to high-precision tubulin cytoskeleton studies and manipulation in neuroscience, motility, and embryology, in the cellular and potentially also the animal contexts. Beyond the power of **SBTub**s as photopharmaceutical MT inhibitors, we believe that their imaging-orthogonal biocompatible isomerisation, high photochemical and biological robustness, druglikeness, and tolerance for drug-important polar functional groups, are promising for broader SBT-based photopharmaceutical tool development addressing biology research applications that are inaccessible to current photopharmaceuticals: especially in the context of intracellular photocontrol of embryos and primary cell isolates of GFP/YFP animal models, as are extensively used in the fields of development and neurodegeneration.

## Methods

*Full and detailed experimental protocols can be found in the Supporting Information*

### Compound synthesis and characterisation

All reactions and characterisations were performed with unpurified, undried, non-degassed solvents and reagents from commercial suppliers (Sigma-Aldrich, TCI Europe N.V., Fisher Scientific etc.), used as obtained, under closed air atmosphere without special precautions. Manual flash column chromatography was performed on Merck silica gel Si-60 (40–63 µm). MPLC flash column chromatography performed on a Biotage Isolera Spektra system, using Biotage prepacked silica cartridges. Thin-layer chromatography (TLC) was run on 0.25 mm Merck silica gel plates (60, F-254), with UV light (254 nm and 365 nm) as visualizing agents. NMR characterisation was performed on a Bruker Ascend 400 (400 MHz & 100 MHz for ^1^H and ^13^C respectively). HRMS was carried out by the Zentrale Analytik of the LMU Munich. Analytical HPLC-MS was performed on an Agilent 1100 SL coupled HPLC-MS system with H_2_O:MeCN eluent gradients through a Thermo Scientific Hypersil GOLD™ C18 column (1.9 µm; 3 × 50 mm) maintained at 25°C, detected on an Agilent 1100 series diode array detector and a Bruker Daltonics HCT-Ultra spectrometer (ESI mode, unit m/z). IR spectra were recorded on a PerkinElmer Spectrum BX II FT-IR system. Full experimental details are given in the Supporting Information.

### Photocharacterisation

UV-Vis absorption spectra measurements in cuvette were acquired on a Varian CaryScan 60 (1 cm pathlength) at room temperature with default compound concentrations of 25 µM and default solvents of PBS at pH ∼7.4 with 10–20% of DMSO as cosolvent. “Star” LEDs (H2A1-models spanning 360–435 nm Roithner Lasertechnik; FWHM ∼25 nm and 260 nm HP-LED Sahlmann Photochemical Solutions) were used for photoisomerisations in cuvette that were also predictive of what would be obtained in LED-illuminated cell culture. Spectra of pure *E* and *Z* isomers were acquired from the HPLC’s inline Agilent 1100 series diode array detector (DAD) over the range 200–550 nm, manually baselining across each elution peak of interest to correct for eluent composition. Two-photon excitation was performed using a mode-locked Ti-Sapphire Laser operating at 780 nm (Spectra Physics, Tsunami) with a pulse repetition frequency of 80 MHz (pulse with <100 fs) and an output power of 0.65 W, coupled into an upright Zeiss Axiotech 100 microscope and focused with a 40× reflective objective (Thorlabs, LMM-40X-UVV, measuring sample transmittance using a HAL 100 illuminator (Carl Zeiss) and a SP300i spectrograph equipped with a MicroMAX CCD camera for signal recording (both Princeton Instruments), between 360 nm and 380 nm.

### Tubulin Polymerisation *in vitro*

99% purity tubulin from porcine brain was obtained from Cytoskeleton Inc. (cat. #T240) and polymerisation assays run according to manufacturer’s instructions. Tubulin was pre-incubated for 10 min at 37 °C with “lit”- or “dark”-**SBTub** (20 µM) in buffer (with 3% DMSO, 10% glycerol); at time zero, GTP (1 mM) was added and the change in absorbance at 340 nm was monitored over 15 mins at 37°C^61^.

### Cell Culture

HeLa cells were maintained under standard cell culture conditions in Dulbecco’s modified Eagle’s medium supplemented with 10% fetal calf serum (FCS), 100 U/mL penicillin and 100 U/mL streptomycin, at 37°C in a 5% CO_2_ atmosphere. Cells were typically transferred to phenol red free medium prior to assays. Compounds and cosolvent (DMSO; 1% final concentration) were added *via* a D300e digital dispenser (Tecan). Cells were either incubated under “dark” (light-excluded) or “lit” conditions (where pulsed illuminations were applied by multi-LED arrays to create and maintain the wavelength-dependent photostationary state isomer ratios throughout the experiment, as previously described^6^).

### MTT Antiproliferation Assay

Cells seeded in 96-well plates at 5,000 cells/well and left to adhere for 24 h were treated with *E*-**SBTub**s under the indicated lighting conditions for 48 h (1% DMSO; six technical replicates). Cells were then treated with 0.5 mg/mL (3-(4,5-dimethylthiazol-2-yl)-2,5-diphenyl tetrazolium bromide (MTT) for 3 h; the medium was aspirated and formazan crystals were re-dissolved in DMSO (100 µL) before measuring absorbance at 550 nm using a FLUOstar Omega microplate reader (BMG Labtech). Absorbance was averaged over the technical replicates, and normalised as viability by reference to the cosolvent-only control (set as 100%) and to zero absorbance (set as 0%).

### Cell Cycle analysis

*E*-**SBTub**s were added to HeLa cells in 24-well plates (seeding density: 50,000 cells/well) and incubated under “dark” or “lit” conditions for 24 h. Cells were collected and stained with 2 µg/mL propidium iodide (PI) at 4°C for 30 min. Following PI staining, cells were analysed by flow cytometry using a FACS Canto II flow cytometer (Becton Dickinson) run by BD FACSDiva software. 30,000 cells were measured per condition and the data were transferred to *Flowing* software for cell cycle analysis. Cells were sorted into sub-G1, G1, S and G_2_/M phase according to DNA content (PI signal).

### Immunofluorescence Staining

HeLa cells seeded on glass coverslips in 24-well plates (50,000 cells/well) were left to adhere for 18 h then treated for 24 h with **SBTub**s under “dark” or “lit” conditions as described above. Cover slips were washed then fixed with 0.5% glutaraldehyde, quenched with 0.1% NaBH_4_, blocked with PBS + 10% FCS, and treated with rabbit alpha-tubulin primary antibody (abcam ab18251; 1:400 in PBS + 10% FCS) for 1 h; after washing with PBS, cells were incubated with goat-anti-rabbit Alexa fluor 488 secondary antibody (Abcam, ab150077; 1:400 in PBS + 10% FCS) for 1 h. After washing with PBS, coverslips were mounted onto glass slides using Roti-Mount FluorCare DAPI (Roth) and imaged with a Zeiss LSM Meta confocal microscope. Images were processed using Fiji software. Postprocessing was performed only to improve visibility. For maximum intensity projections, images were recorded at different focal planes by incrementally stepping through the sample (step size 1-2 µm) and maximum intensity projections were obtained using Fiji software.

### EB3 Comet Assay with whole-sample photoisomerisation

^30^ HeLa cells (12,000 cells/well) were seeded on 8-well ibiTreat µ slides (ibidi) 24 h prior to transfection. Cells were transiently transfected with *EB3-GFP* or *EB3-YFP* plasmids using jetPRIME reagent (Polyplus) according to the manufacturer’s instructions. Cells were imaged 24 h later, under 37°C and 5% CO_2_ atmosphere using an UltraVIEW Vox spinning disc confocal microscope (PerkinElmer) equipped with an EMCCD camera (Hammamatsu, Japan) and operated with *Volocity* software. For photobleaching/wavelength orthogonality experiments, **SBTub** was added cautiously after focussing on cells on the microscope stage, and the compound was incubated for 10 min before imaging; this avoids exposure of the **SBTub** to focusing light, preventing unwanted isomerisation prior to imaging that could falsify results when testing for GFP/YFP orthogonality. Cells were imaged either at 488 nm (GFP; 23% laser power, 400 ms exposure time, 45 frames/min) or 514 nm (YFP; similar parameters); optionally, cells were additionally exposed to interleaved lower-intensity frames (250 ms) of **SBTub**-isomerizing 405 nm light for compound activation (45 frames/min). For EB3 comet statistics, 6 cells per condition from three independent trials were taken. First-order exponential decay curves were fitted to each cell’s comet count timecourses and their values were normalised to 100% at time zero, enabling intercomparison of cells with different starting comet counts (depending on their size, the position of the focal plane, etc). EB3 comets were counted with a *Fiji* software plugin based on the “Find maxima” function from the NIH.

### EB3 Comet Assay with temporally reversible cell-specific photoisomerisation

HeLa cells were transfected with *EB3-GFP* using FuGENE 6 (Promega) according to manufacturer’s instructions. Cells were imaged on a Nikon Eclipse Ti microscope equipped with a perfect focus system (Nikon), a spinning disk-based confocal scanner unit (CSU-X1-A1, Yokogawa) and an Evolve 512 EMCCD camera (Photometrics) with a stage top incubator INUBG2E-ZILCS (Tokai Hit) and lens heating calibrated for incubation at 37°C with 5% CO_2_. Microscope image acquisition was controlled using MetaMorph 7.7, with GFP imaging at 491 nm (0.17 mW, 300 ms every 2 s) and compound activation at 405 nm (77 µW, 100 ms every 90 s), and images were acquired using a Plan Apo VC 60× NA 1.4 oil objective. Comet count analysis performed in ImageJ using the ComDet plugin (E. Katrukha, University of Utrecht, https://github.com/ekatrukha/ComDet).

### Protein production, crystallisation and soaking

The DARPin D1 was prepared as previously described^44^. Tubulin from bovine brain was purchased from the Centro de Investigaciones Biológicas (Microtubule Stabilizing Agents Group), CSIC, Madrid, Spain. The tubulin-DARPin D1 (TD1) complex was formed by mixing the respective components in a 1:1.1 molar ratio. The TD1 complex was crystallised overnight by the hanging drop vapor diffusion method (drop size 2 µL, drop ratio 1:1) at a concentration of 15.9 mg/mL and at 20°C with a precipitant solution containing 18% PEG 3350, 0.2 M ammonium sulfate and 0.1 M bis-tris methane, pH 5.5. All drops were subsequently hair-seeded with crystalline material obtained in previous PEG-screening, which resulted in single and large (0.5 µm) TD1 complex crystals. The crystals were fished and transferred into new precipitant solution drops containing 10% DMSO and compounds (*E-***SBTub2**/*E-***SBTub3**) at final concentrations 2 mM, then 360 nm LED illumination was applied for 5 min to generate *Z*-**SBTub**s *in situ*. After 4 h of soaking in the dark, the crystals were mounted for X-ray diffraction data collection.

### X-ray diffraction data collection, processing and refinement

Data were collected at beamline X06SA at the Swiss Light Source (Paul Scherrer Institute, Villigen, Switzerland). The beam was focused to 30 × 30 µm, the flux was 3 × 10^10^ photons/s and the data collection speed was 2°/s at an oscillation range of 0.2° per frame (see Table S1). For TD1-**SBTub2** and TD1-**SBTub3**, 210° and 220° of data were collected, respectively. Data processing was done with XDS^62^. Due to anisotropy, the data were corrected using the Staraniso server (http://staraniso.globalphasing.org/). The structures were solved by molecular replacement using PDB ID 5NQT as a search model^63^. The ligands and restraints were generated with the grade server (http://grade.globalphasing.org/) using their SMILES annotation. The structures were then refined iteratively in PHENIX^64^ with manual editing cycles in COOT^65^.

## Acknowledgements

This research was supported by funds from the German Research Foundation (DFG: SFB1032 Nanoagents for Spatiotemporal Control project B09 to O.T.-S. and project A08 to T.L.; SFB TRR 152 project P24 number 239283807 to O.T.-S; and Emmy Noether grant to O.T.-S.); the Swiss National Science Foundation (31003A_166608 to M.O.S.); and the Munich Centre for NanoScience initiative (CeNS). We thank S. Schmidt (LMU) for help with synthesis, K. T. Wanner (LMU) for synthetic access and collegial discussions, H. Harz and I. Solvei (LMU microscopy platform CALM), K. Bartel, A. Vollmar, H. Leonhardt and S. Zahler (LMU) for access to general biology and microscopy facilities, and D. Hörl (LMU) for measuring microscope laser power. X-ray diffraction data were collected at the beamline X06SA at the Swiss Light Source (Paul Scherrer Institut, Villigen PSI, Switzerland).

## Author Contributions

L.G. performed synthesis, single-photon photocharacterisation, in vitro studies and coordinated data assembly. Y.K. performed in cellulo studies. M.W. performed tubulin protein production, purification, crystallisation, crystal handling, and X-ray data collection, processing, and refinement. T.W. performed tubulin crystal handling, data collection, data processing and refinement. S.P. performed two-photon photocharacterisation. J.M. performed temporally reversible live cell studies. R.B. performed in vitro tubulin polymerisation assays. N.O. performed tubulin protein production, purification and crystallisation. A.A. supervised temporally reversible cell studies. T.L. supervised two-photon photocharacterisation. M.O.S. supervised protein crystallography. O.T.-S. designed the concept and experiments, supervised all other experiments, coordinated data assembly and wrote the manuscript with input from all authors.

## Additional Information

Supplementary Information accompanies this paper: (1) **Supporting Information PDF** of (i) synthetic protocols; (ii) photocharacterisation; (iii) biochemistry; (iv) cell biology; (v) X-ray crystal structure information; (vi) NMR spectra; and **Supporting Movie M1** (*GFP-orthogonal temporally reversible modulation of MT polymerisation dynamics achieved by pulsed illuminations of SBTub3-treated (5 µM) HeLa cells transfected with EB3-GFP microtubule polymerisation reporter (one pulse of 405 nm laser per 90 s, pulse frames indicated by blue dot, imaging conducted over 9 min, timestamp in mm:ss). Movie is one representative movie from the five acquired and averaged to generate the statistics shown in Fig 5a; selected still images are shown in Fig 5b*).

## Competing Interests

The authors declare no competing interests.

*We dedicate this paper to GFP’s discoverer Osamu Shimomura, whose devoted research has made modern chemical biology possible.*

## Graphical Abstract

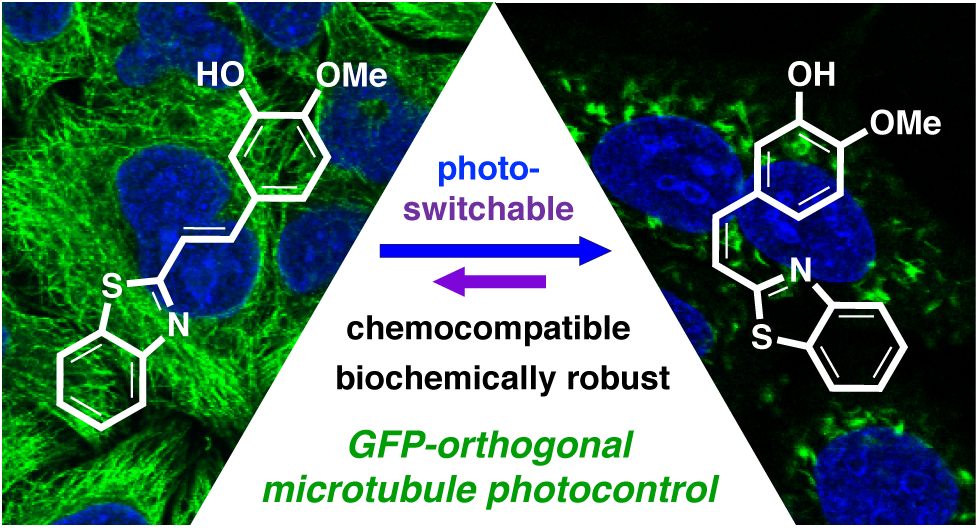

## References

1 Zhou, H. et al. Photoswitching of glass transition temperatures of azobenzene-containing polymers induces reversible solid-to-liquid transitions. Nat. Chem. 9, 145, doi:10.1038/nchem.2625 (2017).

2 Kumar, G. S. & Neckers, D. C. Photochemistry of azobenzene-containing polymers. Chem. Rev. 89, 1915–1925, doi:10.1021/cr00098a012 (1989).

3 Velema, W. A., van der Berg, J. P., Szymanski, W., Driessen, A. J. M. & Feringa, B. L. Orthogonal control of antibacterial activity with light. ACS Chem. Biol. 9, 1969–1974, doi:10.1021/cb500313f (2014).

4 Hüll, K., Morstein, J. & Trauner, D. In Vivo Photopharmacology. Chem. Rev. 118, 10710–10747, doi:10.1021/acs.chemrev.8b00037 (2018).

5 Dong, M., Babalhavaeji, A., Samanta, S., Beharry, A. A. & Woolley, G. A. Red-Shifting Azobenzene Photoswitches for in Vivo Use. Acc. Chem. Res. 48, 2662–2670, doi:10.1021/acs.accounts.5b00270 (2015).

6 Borowiak, M. et al. Photoswitchable Inhibitors of Microtubule Dynamics Optically Control Mitosis and Cell Death. Cell 162, 403–411, doi:10.1016/j.cell.2015.06.049 (2015).

7 Zenker, J. et al. A microtubule-organizing center directing intracellular transport in the early mouse embryo. Science 357, 925–928, doi:10.1126/science.aam9335 (2017).

8 Zenker, J. et al. Expanding Actin Rings Zipper the Mouse Embryo for Blastocyst Formation. Cell 173, 776–791, doi:10.1016/j.cell.2018.02.035 (2018).

9 Dumontet, C. & Jordan, M. A. Microtubule-binding agents: A dynamic field of cancer therapeutics. Nat. Rev. Drug Discov. 9, 897, doi:10.1038/nrd3313 (2010).

10 Peterson, J. R. & Mitchison, T. J. Small Molecules, Big Impact. Chem. Biol. 9, 1275–1285, doi:10.1016/s1074-5521(02)00284-3 (2002).

11 Kingston, D. G. I. Taxol, a molecule for all seasons. Chem. Commun., 867–880, doi:10.1039/B100070P (2001).

12 Janke, C. & Steinmetz, M. O. Optochemistry to control the microtubule cytoskeleton. EMBO J. 34, 2114–2116, doi:10.15252/embj.201592415 (2015).

13 Castle, B. T. & Odde, D. J. Optical Control of Microtubule Dynamics in Time and Space. Cell 162, 243–245, doi:10.1016/j.cell.2015.06.064 (2015).

14 Pettit, G. R. et al. Isolation and structure of the strong cell growth and tubulin inhibitor combretastatin A-4. Experientia 45, 209–211 (1989).

15 Tozer, G. M., Kanthou, C., Parkins, C.S., Hill, S.A. The biology of the combretastatins as tumour vascular targeting agents. Int. J. Exp. Pathol. 83, 21–38, doi:10.1046/j.1365-2613.2002.00211.x (2002).

16 Tron, G. C. et al. Medicinal chemistry of combretastatin A4: Present and future directions. J. Med. Chem. 49, 3033–3044, doi:10.1021/jm0512903 (2006).

17 Woods, J. A., Hadfield, J. A., Pettit, G. R., Fox, B. W. & McGown, A. T. The interaction with tubulin of a series of stilbenes based on combretastatin A-4. Br. J. Cancer 71, 705–711, doi:10.1038/bjc.1995.138 (1995).

18 Scherer, K. M., Bisby, R. H., Botchway, S. W., Hadfield, J. A. & Parker, A. W. Anticancer phototherapy using activation of *E*-combretastatins by two-photon-induced isomerization. J. Biomed. Opt. 20, 051004, doi:10.1117/1.Jbo.20.5.051004 (2015).

19 Jørgensen, K. B. Photochemical oxidative cyclisation of stilbenes and stilbenoids-the Mallory reaction. Molecules 15, 4334–4358, doi:10.3390/molecules15064334 (2010).

20 Morstein, J., Awale, M., Reymond, J.-L. & Trauner, D. Mapping the Azolog Space Enables the Optical Control of New Biological Targets. ACS Central Science, doi:10.1021/acscentsci.8b00881 (2019).

21 Engdahl, A. J. et al. Synthesis, Characterization, and Bioactivity of the Photoisomerizable Tubulin Polymerization Inhibitor azo-Combretastatin A4. Org. Lett. 17, 4546–4549, doi:10.1021/acs.orglett.5b02262 (2015).

22 Sheldon, J. E., Dcona, M. M., Lyons, C. E., Hackett, J. C. & Hartman, M. C. T. Photoswitchable anticancer activity via *trans*-*cis* isomerization of a combretastatin A-4 analog. Org. Biomol. Chem. 14, 40–49, doi:10.1039/c5ob02005k (2016).

23 Rastogi, S. K. et al. Photoresponsive azo-combretastatin A-4 analogues. Eur. J. Med. Chem. 143, 1–7, doi:10.1016/j.ejmech.2017.11.012 (2018).

24 Eguchi, K. et al. Wild-Type Monomeric α-Synuclein Can Impair Vesicle Endocytosis and Synaptic Fidelity via Tubulin Polymerization at the Calyx of Held. J. Neurosci. 37, 6043–6052, doi:10.1523/jneurosci.0179-17.2017 (2017).

25 Singh, A. et al. Polarized microtubule dynamics directs cell mechanics and coordinates forces during epithelial morphogenesis. Nat. Cell Biol. 20, 1126–1133, doi:10.1038/s41556-018-0193-1 (2018).

26 Samanta, S. et al. Photoswitching azo compounds in vivo with red light. J. Am. Chem. Soc. 135, 9777–9784, doi:10.1021/ja402220t (2013).

27 Samanta, S., Babalhavaeji, A., Dong, M.-x. & Woolley, G. A. Photoswitching of ortho-Substituted Azonium Ions by Red Light in Whole Blood. Angew. Chem., Int. Ed. 52, 14127–14130, doi:10.1002/anie.201306352 (2013).

28 Lei, H. et al. Bioactivatable Reductive Cleavage of Azobenzene for Controlling Functional Dumbbell Oligodeoxynucleotides. Bioorg. Chem., 103106, doi:10.1016/j.bioorg.2019.103106 (2019).

29 Boulègue, C., Löweneck, M., Renner, C. & Moroder, L. Redox potential of azobenzene as an amino acid residue in peptides. ChemBioChem 8, 591–594, doi:10.1002/cbic.200600495 (2007).

30 Kleele, T. et al. An assay to image neuronal microtubule dynamics in mice. Nat. Commun. 5, 4827, doi:10.1038/ncomms5827 (2014).

31 Hofmann, A. W. Zimmtsäurederivat des Amidophenylmercaptans. Chemische Berichte, 1235–1238 (1880).

32 Awad, M. K., El-Hendawy, M. M., Fayed, T. A., Etaiw, S. E. H. & English, N. J. Aromatic ring size effects on the photophysics and photochemistry of styrylbenzothiazole. Photochem. Photobiol. Sci. 12, 1220–1231, doi:10.1039/c3pp25367h (2013).

33 Horspool, W. M. & Lenci, F. CRC handbook of organic photochemistry and photobiology. 2nd ed. edn, (CRC Press, 2004).

34 Mishra, A., Thangamani, A., Chatterjee, S., Chipem, F. A. S. & Krishnamoorthy, G. Photoisomerization of trans-2-[4’-(Dimethylamino)styryl]benzothiazole. Photochem. Photobiol. 89, 247–252, doi:10.1111/j.1751-1097.2012.01227.x (2013).

35 El-Hendawy, M. M. et al. Photophysics, photochemistry and thermal stability of diarylethene-containing benzothiazolium species. J. Photochem. Photobiol. 301, 20–31, doi:10.1016/j.jphotochem.2014.12.015 (2015).

36 Shan, Y. S., Zhang, J., Liu, Z., Wang, M. & Dong, Y. Developments of Combretastatin A-4 Derivatives as Anticancer Agents Current Medicinal Chemistry 18, 523–538, doi:10.2174/092986711794480221 (2011).

37 Penthala, N. R., Thakkar, S. & Crooks, P. A. Heteroaromatic analogs of the resveratrol analog DMU-212 as potent anticancer agents. Bioorg. Med. Chem. Lett. 25, 2763–2767, doi:10.1016/j.bmcl.2015.05.019 (2015).

38 Penthala, N. R., Crooks, P. & Sonar, V. Combretastatin analogs. US20160068506A1 (2016).

39 Sailer, A. et al. Hemithioindigos for Cellular Photopharmacology: Desymmetrised Molecular Switch Scaffolds Enabling Design Control over the Isomer-Dependency of Potent Antimitotic Bioactivity. ChemBioChem 20, 1305–1314, doi:10.1002/cbic.201800752 (2019).

40 Gaspari, R., Prota, A. E., Bargsten, K., Cavalli, A. & Steinmetz, M. O. Structural Basis of *cis*- and *trans*-Combretastatin Binding to Tubulin. Chem 2, 102–113, doi:10.1016/j.chempr.2016.12.005 (2017).

41 Lozano, C. M. et al. Cytotoxic anionic tribromo platinum(II) complexes containing benzothiazole and benzoxazole donors: synthesis, characterization, and structure-activity correlation. Inorganica Chimica Acta 271, 137–144, doi:10.1016/S0020-1693(97)05952-5 (1998).

42 Berdnikova, D., Fedorova, O., Gulakova, E. & Ihmels, H. Photoinduced in situ generation of a DNA-binding benzothiazoloquinolinium derivative. Chem. Commun. 48, 4603–4605, doi:10.1039/c2cc30958k (2012).

43 Moreno, J. et al. Two-Photon-Induced versus One-Photon-Induced Isomerization Dynamics of a Bistable Azobenzene Derivative in Solution. The Journal of Physical Chemistry B 119, 12281–12288, doi:10.1021/acs.jpcb.5b07008 (2015).

44 Pecqueur, L. et al. A designed ankyrin repeat protein selected to bind to tubulin caps the microtubule plus end. Proceedings of the National Academy of Sciences 109, 12011, doi:10.1073/pnas.1204129109 (2012).

45 Volgraf, M. et al. Allosteric control of an ionotropic glutamate receptor with an optical switch. Nature Chemical Biology 2, 47, doi:10.1038/nchembio756 (2005).

46 Lester, H. A., Krouse, M. E., Nass, M. M., Wassermann, N. H. & Erlanger, B. F. A covalently bound photoisomerizable agonist: comparison with reversibly bound agonists at Electrophorus electroplaques. The Journal of general physiology 75, 207–232 (1980).

47 Urban, P. et al. Light-Controlled Lipid Interaction and Membrane Organization in Photolipid Bilayer Vesicles. Langmuir 34, 13368–13374, doi:10.1021/acs.langmuir.8b03241 (2018).

48 Frank, J. A., Franquelim, H. G., Schwille, P. & Trauner, D. Optical Control of Lipid Rafts with Photoswitchable Ceramides. Journal of the American Chemical Society 138, 12981–12986, doi:10.1021/jacs.6b07278 (2016).

49 Konrad, D. B., Frank, J. A. & Trauner, D. Synthesis of Redshifted Azobenzene Photoswitches by Late-Stage Functionalization. Chem.–Eur. J 22, 4364–4368, doi:10.1002/chem.201505061 (2016).

50 Bléger, D., Schwarz, J., Brouwer, A. M. & Hecht, S. o-Fluoroazobenzenes as Readily Synthesized Photoswitches Offering Nearly Quantitative Two-Way Isomerization with Visible Light. J. Am. Chem. Soc. 134, 20597–20600, doi:10.1021/ja310323y (2012).

51 Simeth, N. A., Kneuttinger, A. C., Sterner, R. & König, B. Photochromic coenzyme Q derivatives: Switching redox potentials with light. Chem. Sci. 8, 6474–6483, doi:10.1039/c7sc00781g (2017).

52 Hoorens, M. W. H. et al. Iminothioindoxyl as a molecular photoswitch with 100 nm band separation in the visible range. Nat. Comm. 10, 2390, doi:10.1038/s41467-019-10251-8 (2019).

53 Klán, P. et al. Photoremovable Protecting Groups in Chemistry and Biology: Reaction Mechanisms and Efficacy. Chemical Reviews 113, 119–191, doi:10.1021/cr300177k (2013).

54 Reeßing, F. & Szymanski, W. Beyond Photodynamic Therapy: Light-Activated Cancer Chemotherapy. Curr. Med. Chem. 24, 4905–4950, doi:10.2174/0929867323666160906103223 (2017).

55 Bisby, R., Botchway, S., Hadfield, J., McGown, A. & Scherer, K. M. Multi-Photon Isomerisation Of Combretastatins And Their Use In Therapy. WO2013021208 (2013).

56 Gilbert, A. in CRC Handbook of Organic Photochemistry and Photobiology (eds William Horspool & Francesco Lenci) Ch. 33, (CRC Press, 2003).

57 Tozer, G. M. et al. Mechanisms Associated with Tumor Vascular Shut-Down Induced by Combretastatin A-4 Phosphate: Intravital Microscopy and Measurement of Vascular Permeability. Cancer Res. 61, 6413–6422 (2001).

58 Morstein, J. et al. Optical control of sphingosine-1-phosphate formation and function. Nat. Chem. Biol. 15, 623–631, doi:10.1038/s41589-019-0269-7 (2019).

59 Siewertsen, R. et al. Highly Efficient Reversible Z-E Photoisomerization of a Bridged Azobenzene with Visible Light through Resolved S1(nπ*) Absorption Bands. J. Am. Chem. Soc. 131, 15594–15595, doi:10.1021/ja906547d (2009).

60 Reynders, M. et al. PHOTACs Enable Optical Control of Protein Degradation. (2019).

61 Lin, C. M. et al. Interactions of tubulin with potent natural and synthetic analogs of the antimitotic agent combretastatin: a structure-activity study. Molecular Pharmacology 34, 200–208 (1988).

62 Kabsch, W. XDS. Acta crystallographica. Section D, Biological crystallography 66, 125–132, doi:10.1107/S0907444909047337 (2010).

63 Weinert, T. et al. Serial millisecond crystallography for routine room-temperature structure determination at synchrotrons. Nature Communications 8, 542, doi:10.1038/s41467-017-00630-4 (2017).

64 Adams, P. D. et al. PHENIX: a comprehensive Python-based system for macromolecular structure solution. Acta Crystallographica Section D 66, 213–221, doi:doi:10.1107/S0907444909052925 (2010).

65 Emsley, P. & Cowtan, K. Coot: model-building tools for molecular graphics. Acta Crystallographica Section D 60, 2126–2132, doi:doi:10.1107/S0907444904019158 (2004).

