## Supplemental Information for "Photoswitchable microtubule inhibitors enabling robust, GFP-orthogonal optical control over the tubulin cytoskeleton"

ORCIDs: LG: 0000-0003-3963-4847; OTS: 0000-0003-3981-651X

**Keywords**: *tubulin polymerization inhibitor, microtubule dynamics, photopharmacology, colchicine, cytoskeleton.*

<u>Authorship Statement</u>: Following the ICMJE guidelines, the authors declare their roles in the present work as follows: L.G. performed synthesis, single-photon photocharacterisation, *in vitro* studies and coordinated data assembly. Y.K. performed *in cellulo* studies. M.W. performed tubulin protein production, purification, crystallisation, crystal handling, and X-ray data collection, processing, and refinement. T.W. performed tubulin crystal handling, data collection, data processing and refinement. S.P. performed two-photon photocharacterisation. J.M. performed temporally reversible live cell studies. R.B. performed *in vitro* tubulin polymerisation assays. N.O. performed tubulin protein production, purification and crystallisation. A.A. supervised temporally reversible cell studies. T.L. supervised two-photon photocharacterisation. M.O.S. supervised protein crystallography. O.T.-S. designed the concept and experiments, supervised all other experiments, coordinated data assembly and wrote the manuscript with input from all authors.

<u>Funding Statement</u>: This research was supported by funds from the German Research Foundation (DFG: SFB1032 *Nanoagents for Spatiotemporal Control* project B09 to O.T.-S. and project A08 to T.L.; SFB TRR 152 project P24 number 239283807 to O.T.-S; and Emmy Noether grant to O.T.-S.); the Swiss National Science Foundation (31003A_166608 to M.O.S.); and the Munich Centre for NanoScience initiative (CeNS).

### Part A: Chemical Synthesis

#### *Conventions*

Abbreviations: The following abbreviations are used: Hex – distilled isohexanes, EA – ethyl acetate, Me – methyl, MeCN – acetonitrile, DMSO – dimethylsulfoxide, PBS – phosphate buffered saline aqueous buffer.

Safety Hazards: no remarkable safety hazards were encountered.

Reagents and Conditions: Unless stated otherwise, (1) all reactions and characterisations were performed with unpurified, undried, non-degassed solvents and reagents, used as obtained, under closed air atmosphere without special precautions; (2) "hexane" used for chromatography was distilled from commercial crude isohexane fraction by rotary evaporation; (3) "column" and "chromatography" refer to manual flash column chromatography on Merck silica gel Si-60 (40–63 µm); (4) "MPLC" refers to flash column chromatography purification on a Biotage Isolera Spektra system, using prepacked silica cartridges purchased from Biotage; (5) procedures and yields are unoptimised; (6) yields refer to isolated chromatographically and spectroscopically pure materials; (7) all eluent and solvent mixtures are given as volume ratios unless otherwise specified, thus "1:1 Hex:EA" indicates a 1:1 mixture (by volume) of hexanes and ethyl acetate; (8) chromatography eluents e.g. "3:1 → 1:1" indicate a stepwise or continual gradient of eluent composition.

Thin-layer chromatography (TLC) was run on 0.25 mm Merck silica gel plates (60, F-254), typically with Hex:EA eluents. UV light (254 nm) was used as a visualising agent, with cross-checking by 365 nm UV lamp for **SBTub** fluorescence. TLC characterisations are abbreviated as $\mathbf{R_f}$ = 0.64 (UV 254 nm, EA:Hex = 1:1).

NMR: Standard NMR characterisation was by $^{1}$H- and $^{13}$C-NMR spectra on a Bruker Ascend 400 (400 MHz & 100 MHz for $^{1}$H and $^{13}$C respectively). Chemical shifts (δ) are reported in ppm calibrated to residual non-perdeuterated solvent as an internal reference[1]. Peak descriptions singlet (s), doublet (d), triplet (t), and multiplet (m) are used. NMR spectra are given in Part F.

HRMS: High resolution mass spectrometry (HRMS) was carried out by the Zentrale Analytik of the LMU Munich using ESI ionisation in the positive mode.

IR Spectroscopy: IR spectra were recorded on a PerkinElmer Spectrum BX II FT-IR system. Both solids and liquids were directly applied as thin films of neat substance on the ATR unit. The measured wavenumbers are reported with relative intensities, abbreviated by s (strong), m (medium) and w (weak).

#### ***Synthesis procedures***

**2-(2,3,4-trimethoxystyryl)benzothiazole (SBTub1)**

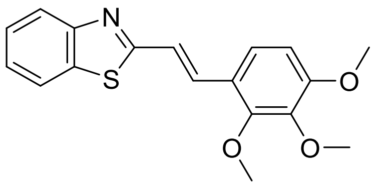

2-methylbenzothiazole (0.12 mL, 1 mmol) and 2,3,4-trimethoxybenzaldehyde (196 mg, 1 mmol, 1 eq) were dissolved in DMSO (2 mL). NaOMe (5.4 M in MeOH, 185 $\mu$L, 1 mmol, 1 eq) was added and the reaction was stirred overnight. $H_2O$ (10 mL) was added. The mixture was extracted with ethyl acetate (3 × 15 mL). The combined organic layers were washed with brine (10 mL) and dried over $MgSO_4$. Flash column purification of the crude material (EA:Hex = 3:7) yielded **SBTub1** as light yellowish solid (298 mg, 0.91 mmol, 91%).

**$R_f$** = 0.64 (UV 254 nm, EA:Hex = 1:1). **HRMS (ESI$^+$):** 328.10002 calculated for $C_{18}H_{18}NO_3S^+$ $[M+H]^+$, 328.10003 found. **$^1$H-NMR (400 MHz, $CDCl_3$):** δ = 7.98 (d, *J* = 8.1 Hz, 1H), 7.84 (d, *J* = 8.0 Hz, 1H), 7.71 (d, *J* = 16.4 Hz, 1H), 7.45 (t, *J* = 8.3 Hz, 1H), 7.41 (d, *J* = 16.4 Hz, 1H), 7.35 (t, *J* = 8.2 Hz, 1H), 7.35 (d, *J* = 8.8 Hz, 1H), 6.73 (d, *J* = 8.8 Hz, 1H), 3.98 (s, 3H), 3.91 (s, 3H), 3.90 (s, 3H) ppm. **$^{13}$C-NMR (100 MHz, $CDCl_3$):** δ = 168.2, 155.0, 154.0, 152.8, 142.5, 134.3, 132.9, 126.4, 125.3, 122.9, 122.6, 122.4, 121.6, 121.5, 108.0, 61.6, 61.1, 56.2 ppm. **IR (FT, ATR):** ṽ = 2946 (w), 2835 (w), 1619 (w), 1588 (m), 1497 (m), 1455 (m), 1435 (m), 1412 (m), 1288 (s), 1254 (m), 1195 (m), 1093 (s), 1042 (m), 1014 (m), 968 (m), 945 (m), 914 (w), 851 (w), 798 (m), 754 (m) $cm^{-1}$.

**2-(3,4,5-trimethoxystyryl)benzothiazole (SBTub2)**

Note that this compound was reported - although only in the *trans* state - in 2014-2015 by Penthala *et al.*, as "Example Compound 4 (Formula (I)(d)" in their patent WO2014/172363[2] and as compound 13 in their paper[3]. In these reports it was synthesised and biologically evaluated for antiproliferative activity in 2D cell culture (patent Table 1, showing growth inhibition $GI_{50}$ < 1 $\mu$M in almost all reported cell lines (NCI60 panel), down to $GI_{50}$ = 40 nM. In the paper, the authors rationalise their finding by computer docking of the *trans*-state to the colchicine site of tubulin using Sybyl software and conclude, "[compound 13] exhibited significant growth inhibition against most of the human cancer cells in the 60-cell panel, and the results from the molecular modeling studies are consistent with the *in vitro* anti-cancer activit[y] …being mediated via binding to the colchicine binding site on tubulin… [it was] considered as [an] important lead compound for further development as [an] anti-cancer drug".

The authors implicitly argue against the widely-reported requirement[4,5] for colchicine-site binders of this type to be *cis*oid isomers, which has been structurally rationalised by us[6].

Penthala *et al*'s prior reports of **SBTub2** properties also argue directly against our cellular bioactivity assays and our experimental ligand:protein crystallisation results, in which we show that *trans*-**SBTub2** has no specific cytotoxicity whereas *cis*-**SBTub2** is strongly bioactive. We consider that Penthala *et al* may initially have synthesised **SBTub2** purely as its *trans*-isomer, but that the possibility of its photoisomerisation-dependent bioactivity (which has not been reported before our current work) escaped their attention. This may have led them to perform bioactivity tests with insufficiently rigorous conditions to be certain of what compound was being tested in what situation; and that also would nullify the predictive value of their reports. We consider that their work has therefore, from conceptual grounds, not delivered an understanding of these compounds' bioactivity, so we here wish to report upon the **SBTub** independently of Penthala *et al.*'s reported cellular activities and docking study, by careful and rationalised studies controlling for isomerisation and experimentally verifying the docking mode of these ligands.

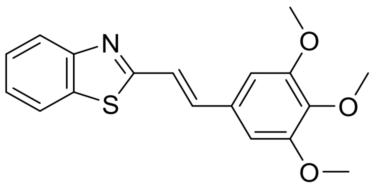

2-methylbenzothiazole (0.12 mL, 1 mmol) and 3,4,5-trimethoxybenzaldehyde (196 mg, 1 mmol, 1 eq) were dissolved in DMSO (2 mL). NaOMe (5.4 M in MeOH, 185 $\mu$L, 1 mmol, 1 eq) was added and the reaction mixture was stirred overnight. $H_2O$ (10 mL) was added and the mixture was extracted with ethyl acetate (3 × 15 mL). The combined organic layers were washed with brine (10 mL) and dried over $MgSO_4$. Flash column purification of the crude material (EA:Hex = 3:7) yielded **SBTub2** as faint-yellow solid (158 mg, 0.48 mmol, 48%).

**$R_f$** = 0.68 (UV 254 nm, EA:Hex = 1:1). **HRMS (ESI$^+$):** 328.10002 calculated for $C_{18}H_{18}NO_3S^+$ $[M+H]^+$, 328.10002 found. **$^1$H-NMR (400 MHz, $CDCl_3$):** δ = 7.99 (d, $J$ = 8.1 Hz, 1H), 7.86 (d, $J$ = 8.0 Hz, 1H), 7.47 (t, $J$ = 7.1 Hz, 1H), 7.45 (d, $J$ = 16.1 Hz, 1H), 7.38 (t, $J$ = 7.0 Hz, 1H), 7.33 (d, $J$ = 16.1 Hz, 1H), 6.82 (s, 2H), 3.92 (s, 6H), 3.90 (s, 3H) ppm. **$^{13}$C-NMR (100 MHz, $CDCl_3$):** δ = 167.0, 153.9, 153.7, 139.6, 137.7, 134.4, 131.1, 126.5, 125.5, 123.0, 121.7, 121.7, 104.6, 61.1, 56.3 ppm. **IR (FT, ATR):** ṽ = 2997 (w), 2966 (w), 2934 (w), 2833 (w), 1626 (w), 1581 (m), 1504 (m), 1452 (m), 1417 (m), 1329 (m), 1240 (m), 1202 (w), 1155 (w), 1126 (s), 1005 (m), 983 (w), 964 (m), 812 (m), 758 (m), 729 (m) $cm^{-1}$.

<u>**5-(2-(benzothiazol-2-yl)vinyl)-2-methoxyphenol (SBTub3)**</u>

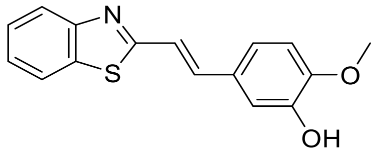

2-methylbenzothiazole (0.25 mL, 2 mmol), 3-hydroxy-4-methoxybenzaldehyde (304 mg, 2 mmol, 1 eq) were dissolved in DMSO (4 mL). NaOMe (5.4 M in MeOH, 741 μL, 4 mmol, 2 eq) was added. After stirring overnight sat. aq. $NH_4Cl$ solution (30 mL) was added and the reaction mixture was extracted with EA (3 × 20 mL). The combined organic layers were washed with water (20 mL), brine (20 mL) and dried over $MgSO_4$. The crude material was purified by normal phase MPLC using gradient elution (EA:Hex = 0:100 → 100:0) to yield **SBTub3** as slightly yellowish solid (502 mg, 1.77 mmol, 89%).

**$R_f$** = 0.55 (UV 254 nm, EA:Hex = 1:1). **HRMS (ESI$^+$):** 284.07398 calculated for $C_{16}H_{14}NO_2S^+$ $[M+H]^+$, 284.07384 found. **$^1$H-NMR (400 MHz, $CDCl_3$):** δ = 7.98 (d, *J* = 8.1 Hz, 1H), 7.85 (d, *J* = 8.0 Hz, 1H), 7.46 (t, *J* = 7.1 Hz, 1H), 7.44 (d, *J* = 16.2 Hz, 1H), 7.36 (t, *J* = 7.0 Hz, 1H), 7.26 (d, *J* = 16.1 Hz, 1H), 7.20 (d, *J* = 2.1 Hz, 1H), 7.09 (dd, *J* = 8.3, 2.1 Hz, 1H), 6.88 (d, *J* = 8.3 Hz, 1H), 5.73 (s, 1H, *OH*), 3.93 (s, 3H) ppm. **$^{13}$C-NMR (100 MHz, $CDCl_3$):** δ = 167.5, 153.9, 148.1, 146.1, 137.7, 134.4, 129.2, 126.4, 125.3, 122.9, 121.6, 120.9, 120.5, 112.7, 110.8, 56.2 ppm. **IR (FT, ATR):** ṽ = 3053 (w), 2846 (w), 1626 (w), 1598 (w), 1585 (w), 1522 (m), 1483 (w), 1435 (s), 1358 (w), 1308 (w), 1293 (m), 1245 (m), 1231 (m), 1199 (s), 1164 (s), 1123 (s), 1026 (m), 981 (w), 946 (s), 923 (w), 838 (w), 797 (s), 765 (s), 731 (m) $cm^{-1}$.

<u>**4-(2-(benzothiazol-2-yl)vinyl)-2-methoxyphenol (SBTub4)**</u>[7]

Note that this compound has previously been reported in the *trans* state by Lozano *et al.* as compound **1i**[7], where it was used as a monodentate ligand in a tribromo platinum(II) complex (**2i**). In that report it was assessed for cytotoxicity as a free ligand and also when complexed, in both cases only in *trans* and without consideration of isomerisation. In this work its role was to serve as a designed inactive biological control compound *in the* cis *state,* being a permutation control of **SBTub3** that is designed to be *cis*-active. Therefore we proceeded to synthesise and evaluate it ourselves, paying particular attention to the isomer state.

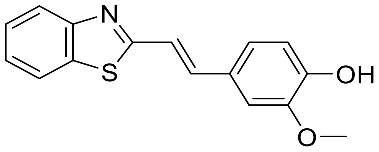

2-methylbenzothiazole (0.41 mL, 3.3 mmol), 4-hydroxy-3-methoxybenzaldehyde (0.5 g, 3.3 mmol, 1.0 eq) and NaOMe (5.4 M solution in MeOH, 1.2 mL, 6.5 mmol, 2 eq) were dissolved in DMSO (6.5 mL). After stirring overnight, sat. aq. $NH_4Cl$ solution (30 mL) was added and the reaction mixture was extracted with EA (3 × 20 mL). The combined organic layers were washed with water (20 mL), brine (20 mL) and dried over $MgSO_4$. The crude material was purified by normal phase MPLC using gradient elution (EA:Hex = 0:100 → 100:0) and $\lambda_{collect}$ = 360 nm to yield **SBTub4** as a yellow solid (256 mg, 0.9 mmol, 28%).

**$R_f$** = 0.6 (UV 254 nm, EA:Hex = 1:1). **HRMS (ESI$^+$):** 284.07398 calculated for $C_{16}H_{14}NO_2S^+$ $[M+H]^+$, 284.07385 found. **$^1$H-NMR (400 MHz, $CDCl_3$):** δ = 7.98 (d, *J* = 8.1 Hz, 1H), 7.85 (d, *J* = 7.9 Hz, 1H), 7.46 (t, *J* = 7.1 Hz, 1H), 7.45 (d, *J* = 15.7 Hz, 1H), 7.36 (t, *J* = 7.1 Hz, 1H), 7.27 (d, *J* = 16.1 Hz, 1H), 7.13 – 7.07 (m, 2H), 6.95 (d, *J* = 8.7 Hz, 1H), 6.02 (s, 1H, *OH*), 3.94 (s, 3H) ppm. **$^{13}$C-NMR (100 MHz, $CDCl_3$):** δ = 167.5, 153.9, 147.5, 147.1, 138.0, 134.3, 128.1, 126.4, 125.3, 122.9, 122.4, 121.6, 120.0, 115.0, 108.8, 56.1 ppm. **IR (FT, ATR):** ṽ = 3004 (w), 2829 (w), 2643 (w), 1623 (m), 1587 (s), 1518 (s), 1456 (m), 1431 (m), 1399 (m), 1338 (w), 1308 (m), 1286 (s), 1254 (s), 1220 (m), 1197 (w), 1169 (w), 1138 (m), 1112 (s), 1064 (w), 1037 (m), 1014 (w), 967 (m), 922 (w), 908 (w), 829 (m), 802 (m), 781 (w), 751 (s), 722 (m) $cm^{-1}$.

#### Part B: Photocharacterisation in vitro

##### HPLC for UV-Vis spectroscopy on separated isomers

Analytical high-performance liquid chromatography (HPLC) was performed on an Agilent 1100 SL coupled HPLC system with (a) a binary pump to deliver $H_2O$:MeCN eluent mixtures containing 0.1% formic acid at a 0.4 mL/min flow rate, (b) Thermo Scientific Hypersil GOLD™ C18 column (1.9 µm; 3 × 50 mm) maintained at 25°C, whereby the solvent front eluted at $t_{ret}$ = 0.5 min, (c) an Agilent 1100 series diode array detector, which was used to acquire peak spectra of separated compound isomers in the range 200–550 nm after manually baselining across each elution peak of interest to correct for eluent composition effects. Run conditions were a linear gradient of $H_2O$:MeCN eluent composition from 90:10 through to 1:99, applied during the separation phase (first 5 min), then 0:100 for 2 min for flushing; the column was (re)equilibrated with 90:10 eluent mixture for 2 min before each run.

##### UV-Vis spectrophotometry of bulk samples

Absorption spectra in cuvette (“UV-Vis”) were acquired on a Varian CaryScan 60 (1 cm pathlength). For photoisomerisation measurements, Hellma microcuvettes (108-002-10-40) taking 500 µL volume to top of optical window were used with test solution such that the vertical pathlength of the isomerization light is less than 7 mm to the bottom of the cuvette, with the default test solution concentrations of 25 µM. Measurements were performed by default in PBS at pH ~7.4 with 10–20% of DMSO. Photoisomerisations and relaxation rate measurements were performed at room temperature. “Star” LEDs (H2A1-models spanning 360–435 nm from Roithner Lasertechnik and a 260 nm HP-LED from Sahlmann Photochemical Solutions) were used for photoisomerisations in the cuvette that were also predictive of what would be obtained in LED-illuminated cell culture.

#### *E- and Z-isomers' absorption spectra and photostationary state equilibria*

Pure *E* and *Z* isomer spectra of **SBTub**s (Fig S1a) were obtained on the HPLC DAD as outlined in *HPLC*. Whole-sample absorption spectra of **SBTub**s at different photostationary state equilibria (Fig S1b) were measured as outlined in *Spectrophotometry*.

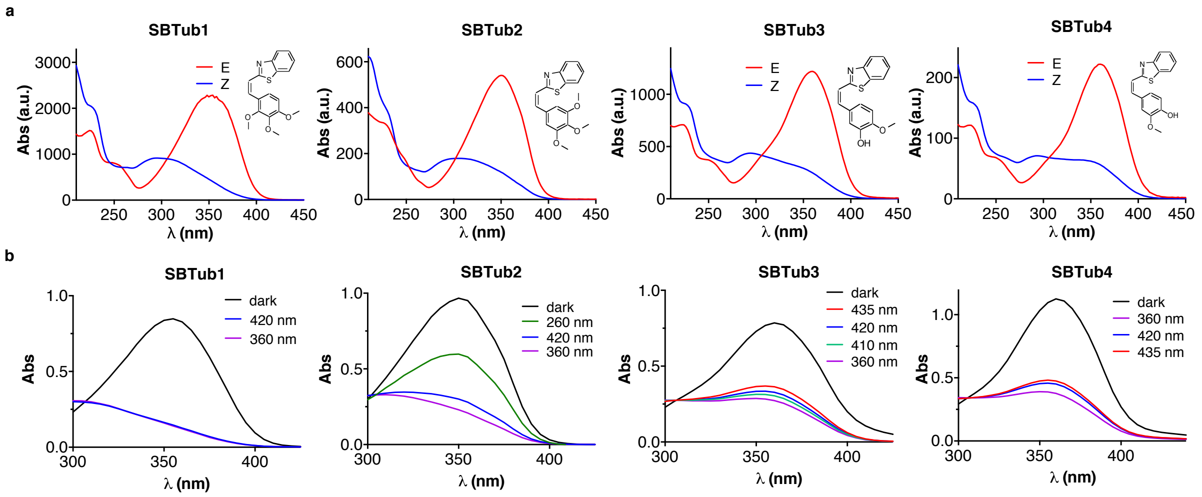

Figure S1: **a** Pure *E* and *Z* spectra of **SBTub1-4** from inline HPLC-DAD. **b** Absorption spectra of **SBTub** samples under saturating illumination with different wavelengths generating the given photostationary state (PSS) equilibria for **SBTub1-4**, from UV-Vis spectrophotometry, in 90% PBS:10% DMSO (**SBTub**s at 25 $\mu$M).

This illustrates the efficient photoswitching to nearly-all-*Z* that may be achieved by a range of violet/near-UV wavelengths.

#### *SBTubs' absorption spectra and photostationary states*

**SBTubs** feature substantially solvent- and pH-independent absorption spectra and photostationary states (Fig S2).

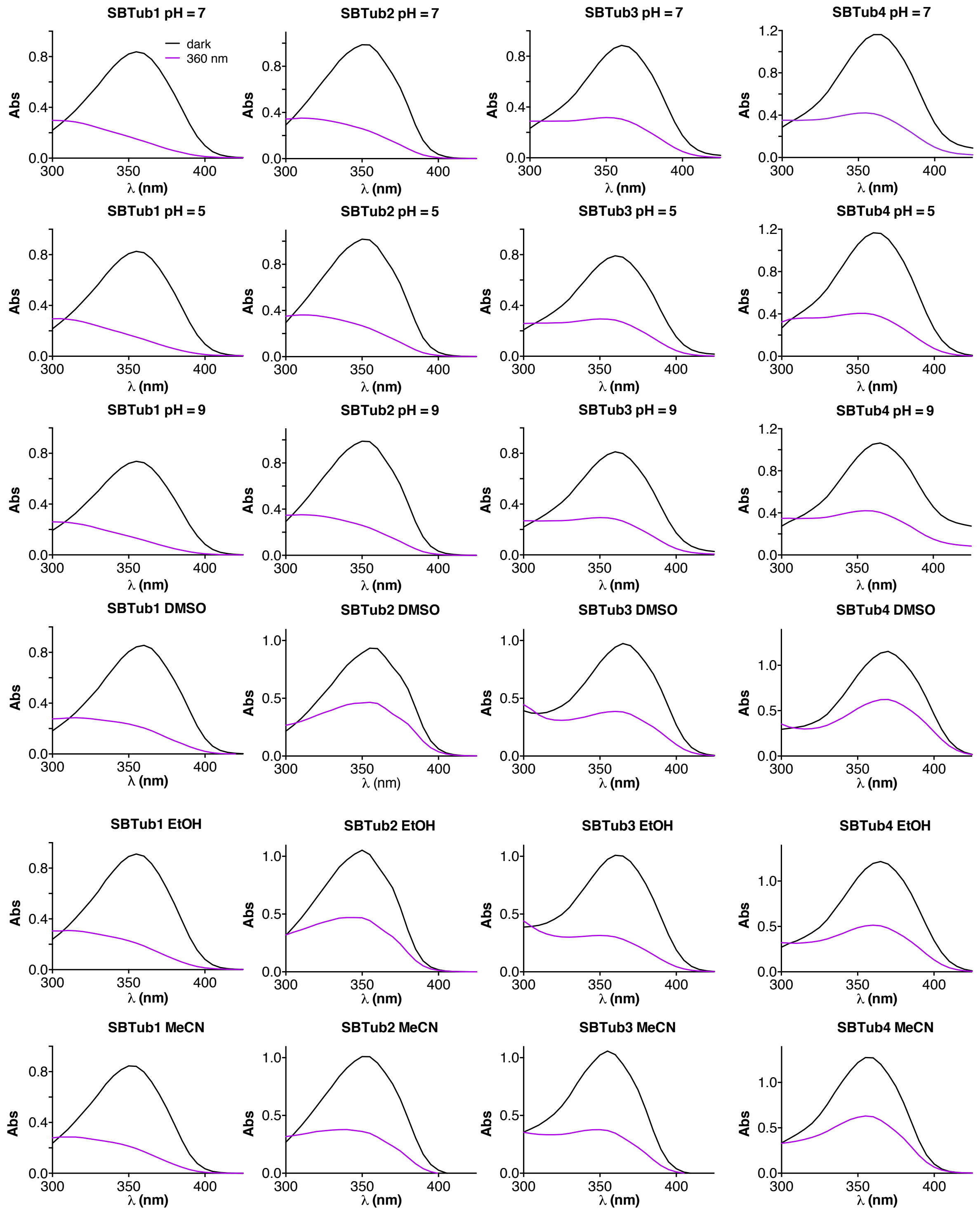

Figure S2: Absorption spectra of **SBTub1-4** in the dark (black) and after photoisomerization with 360 nm light (purple) in 10% DMSO+90% PBS (pH = 7, 5 or 9), or in pure DMSO, EtOH or MeCN (25 $\mu$M).

This outlines their robustness of photoresponse with respect to variations of local conditions as may be encountered in biology, with e.g. localisation into different cellular compartments or changes of environmental dielectric constant during partitioning. We feel that this particularly recommends the SBT scaffold as a reproducible photoswitch for biology use.

##### *Thermal relaxation of p-hydroxy SBTub4 differs from that of non-p-hydroxy SBTubs*

*para*-hydroxy **SBTub4** was the only **SBTub** to show appreciable spontaneous *Z→E* isomerization at 25°C in aqueous media (Fig S3a), and this only under basic aqueous conditions (pH ~9): the non-*para*-hydroxy **SBTub**s did not show relaxation under these conditions (Fig S3b). We presume **SBTub4**'s pH-dependent relaxation rate acceleration results from resonance between the phenolate form (where the bridging bond is a C=C double bond) and its quinoidal form (bridging C-C single bond, for which free rotation to the thermodynamically more stable *transoid* conformation could occur).

a) b)

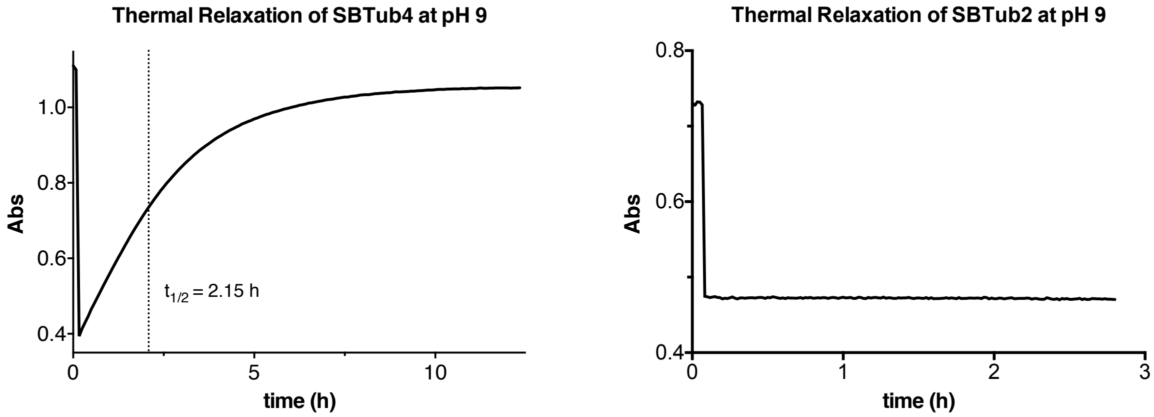

Figure S3: Thermal relaxation of **a SBTub4** and **b SBTub2** (25 $\mu$M) in PBS at pH ~9 with 10% DMSO at 25°C.

In azobenzenes the choice of substituents in *ortho* or *para* to the diazene is restricted, if photoswitching is to be performed in predominantly aqueous environments (e.g. in the cytosol in biological systems) since *o*/*p*-hydroxyl, -thiol or -amine groups give fast thermal relaxation, often complete in under a second, and in many cases their *Z* states cannot even be observed in aqueous environments.[8,9] However these groups are powerful interaction points for docking to proteins, which it would be desirable to be able to place freely around the azobenzene scaffold while maintaining reliable photoswitchability. This can be achieved using SBTs, which we feel is a particular advantage for SBTs as a new photoswitch scaffold for chemical biology.

#### ***Photostability of SBTubs – isomerisation and cycloaddition***

To examine whether degradative photochemical side reactions (e.g. 6-π electrocyclisation, [2+2] cycloaddition, etc) could affect the **SBTub**s during chemical biology use, a sample of **SBTub2** (3.3 mg) in $d_6$-DMSO (0.5 mL) in a quartz NMR tube capped under air, was irradiated with 350 nm UV within a Rayonet RPR-200 photochemistry reactor (Raynonet RPR-3500A lamps, air cooled with a cooling fan). The NMR tube was mounted to an RMA-500 Merry Go-Round unit to maintain an approximate distance of 2 cm from the UV-lamps. An NMR was taken before and after constant irradiation for 1 h, showing exclusively a mixture of *E*- and *Z*-isomers (no photodegradation or cycloaddition). During standard biological assays, illuminations are conducted with far lower photon fluxes (e.g. < 1 mW/cm$^2$ pulses from low-power LEDs for total <1 min illuminated time over long-term viability experiments), therefore it can be assumed that negligible photodegradation of the **SBTub**s should occur. Only after constant Rayonet reactor irradiation overnight, were some photodegradation products (primarily [2+2] cycloaddition product) evidenced by NMR and LC-MS analysis (detected by HPLC coupled to a Bruker Daltonics HCT-Ultra spectrometer used in ESI mode, unit m/z) although the major compound present was still the **SBTub** (as a mixture of *E*- and *Z*-isomers) (Fig S4-S5).

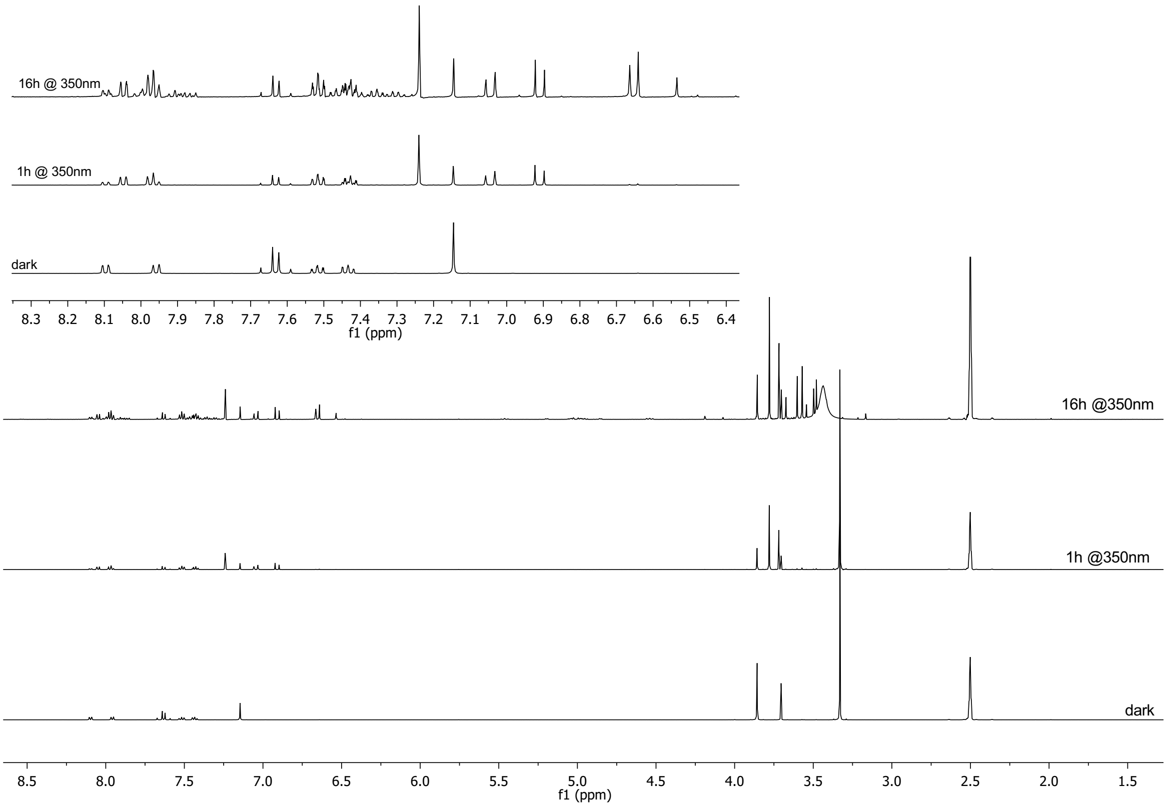

Figure S4: NMR of **SBTub2** in the dark (all-*trans*) and after irradiation with 350 nm UV in Rayonet photoreactor for 1 h and for 16 h, with zoom-ins on the aromatic region that better illustrates isomerisation/cycloaddition.

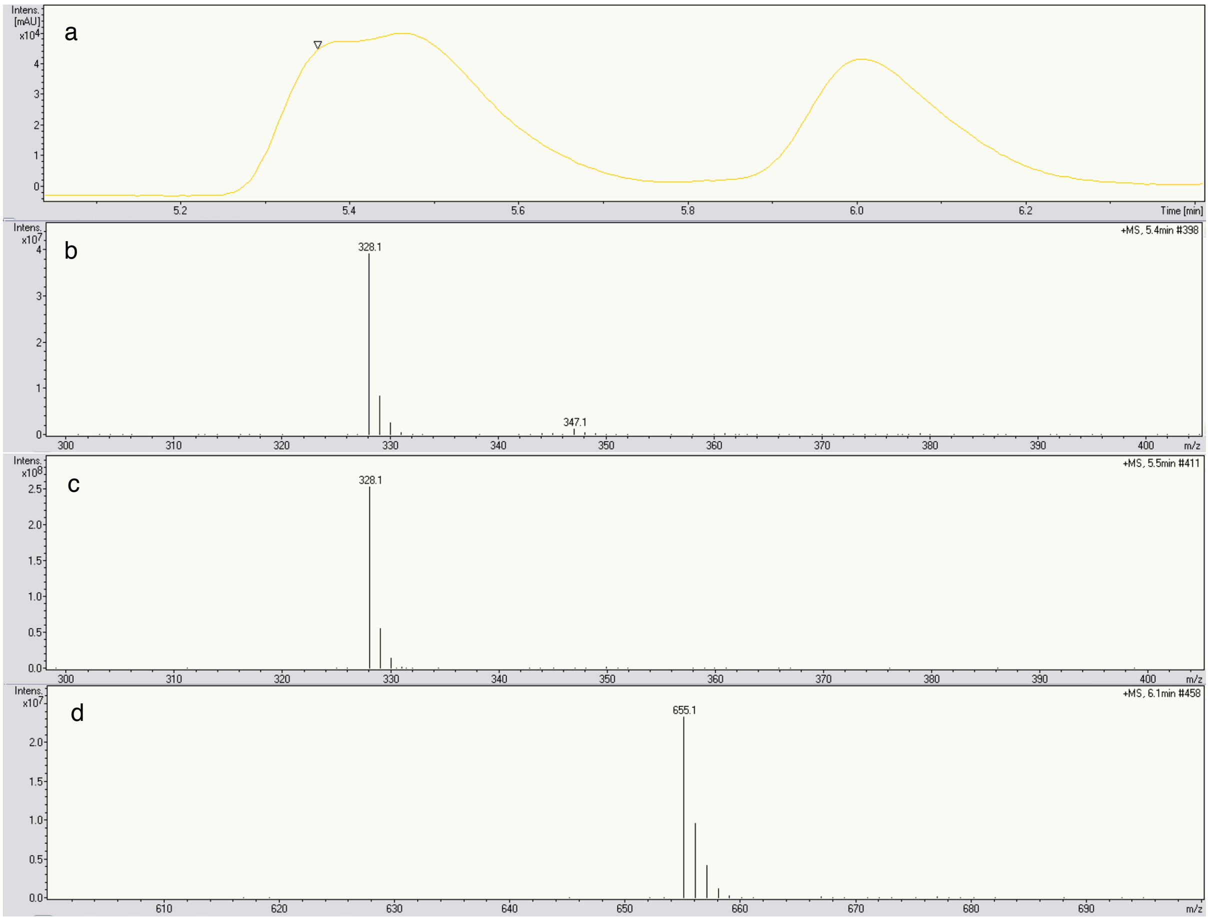

Figure S5: **a** HPLC-MS trace of **SBTub2** after irradiation with 350 nm in the UV photoreactor for 16 h displaying all major peaks. **b** Mass spectrum at 5.4 min showing Z-**SBTub2**. **c** Mass spectrum at 5.5 min showing E-**SBTub2** **d** Mass spectrum at 6.1 min showing [2+2] cycloaddition product.

#### ***PSS measurements and their comparison to simulated PSS(λ)***

For photopharmaceutical assays in biology it is helpful to straightforwardly estimate the *E/Z* ratio at any wavelength's photostationary state (PSS), to choose optimal wavelength conditions for illumination during biological assays or to understand the limits of what is possible on a given setup. The absorption spectrum of a bulk sample at a certain PSS is a linear combination of spectra of the constituent *E* and *Z*-isomers. Therefore we compared experimentally measured UV-spectra from different PSS equilibria, with calculated PSS spectra obtained by assuming (1) that quantum yields for *E→Z* and *Z→E* isomerisations are approximately equal over the measured photoisomerisation region (which is experimentally verifiable by confirming that illumination at the isosbestic point gives an *E*/*Z* ratio approximately 1:1), and that (2) the isomer spectra measured inline in HPLC are approximately identical to those that underlie UV-Vis measurements in biological media in cuvette (which we verified by solvent indepency measurements, as shown in Fig S2, comparing HPLC eluent solvent to biological media): these assumptions allow us to simulate PSS spectra on the basis of HPLC traces as follows (see also Fig S6):

a) Taking experimental PSS absorption spectra on a UV-Vis spectrometer under different illumination wavelengths, thus also determining the isosbestic point $\lambda_{iso}$.

b) Obtaining absorption spectra ε*E*(λ) and ε*Z*(λ) of the pure *E* and *Z*-isomers from an inline diode array detector (DAD) of an HPLC system. Absorption spectra of both isomers are scaled by multiplication in order to cross at $\lambda_{iso}$, generating data ε'*E*(λ) and ε'*Z*(λ) in arbitrary units, such that ε'*E*(λ) / ε'*Z*(λ) = ε*E*(λ) / ε*Z*(λ) across all wavelengths.

c) Estimation of $\text{Frac}[E]_{PSS}(\lambda)$, the fraction of *E*-isomer that should be generated at photostationary state illumination at wavelength λ (assumes identical quantum yields for *E→Z* and *Z→E* isomerisations) *via*

$$\text{Frac}[E]_{PSS}(\lambda) = \frac{\varepsilon' Z(\lambda)}{\varepsilon' Z(\lambda) + \varepsilon' E(\lambda)}$$

d) Simulating absorption spectra in arbitrary units *via*

$$Abs_{SIM}(\lambda) = \text{Frac}[E]_{\text{PSS}}(\lambda) \times \varepsilon' E(\lambda) + \left(1 - \text{Frac}[E]_{\text{PSS}}(\lambda)\right) \times \varepsilon' Z(\lambda)$$

and comparing the predicted-PSS *curve shape* to that of experimental data (more important than comparing single absolute absorption values since more robust to variation). When the curve shapes match across a range of PSS wavelengths, we assume that we can interpolate other PSS *E/Z* ratios under any bracketed illumination wavelength, from the measured isolated-isomer spectra (HPLC). This was found to be exquisitely accurate for **SBTub**s.

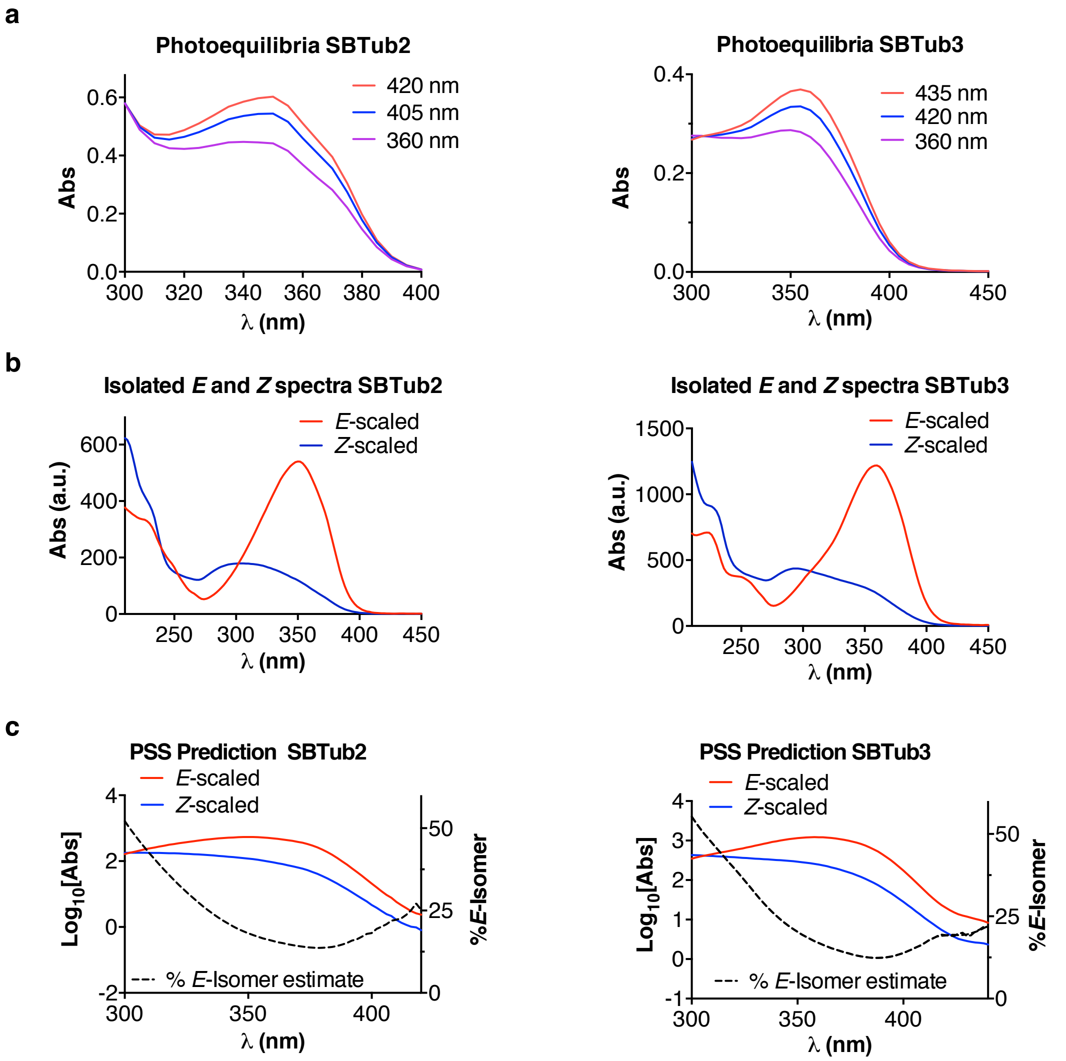

| PSS %E @ λ (nm) | | | | | | | | |
|---|---|---|---|---|---|---|---|---|
| | 260 | 280 | 300 | 320 | 360 | 405 | 420 | *445* |
| **SBTub2** | 58 | 70 | 52 | 34 | 15 | 20 | 25 | *n.d.* |
| **SBTub3** | 53 | 70 | 50 | 40 | 17 | 15 | 17 | *n.d.* |

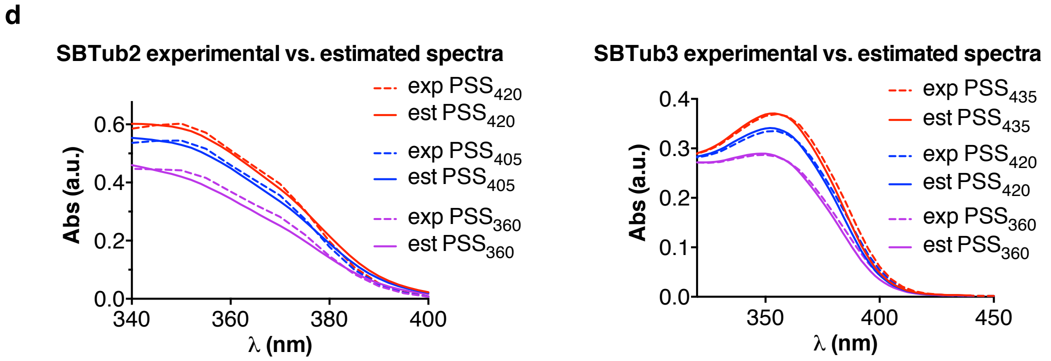

Figure S6: **a** Experimental sample PSS absorption spectra in biological media. **b** HPLC spectra of pure isomers. **c** simulated %E at PSS using HPLC spectra **d** correspondence of simulated (est. PSS) and measured PSS (exp. PSS) sample absorption spectra for both **SBTub2** and **SBTub3** supports PSS interpolation from HPLC spectra.

##### ***Two-photon excitation***

Two-photon excitation of **SBTub3** (5 mM in DMSO) was performed using a mode-locked Ti-Sapphire Laser operating at 780 nm (Spectra Physics, Tsunami) with a pulse repetition frequency of 80 MHz (pulse with <100 fs) and an output power of 0.65 W. The laser was coupled into an upright Microscope (Zeiss Axiotech 100) and focused with a 40× reflective objective (**Thorlabs, LMM-40X-UVV**) onto the sample. Transmittance through the sample was measured using a HAL 100 illuminator (Carl Zeiss) and a SP300i spectrograph equipped with a MicroMAX CCD camera for signal recording (both Princeton Instruments), between 360 nm and 380 nm. This spectral range was chosen to be around the absorbance maximum of **SBTub3** in DMSO (~366 nm, cf. Fig S2). Spectra were recorded with low intensity incident light, and an exposure time of only 1 s to prevent any unintended photoisomerization of the molecules during the measurement (and as a control, 12 min of continuous spectral recording produced an *E* to *Z* isomerisation of only 4%). The sample was measured in a home-made microcuvette with optical path length d=0.1 mm and volume ~7.8 µL. Photoisomerization was analyzed by measuring the change in light intensity after passing through the sample ($I_{SBTub3}$) in relation to light passing through a reference sample of only DMSO ($I_{DMSO}$). The absorbance of the sample was calculated using $\mathbf{A = -log_{10}(I_{SBTub3} / I_{DMSO})}$:

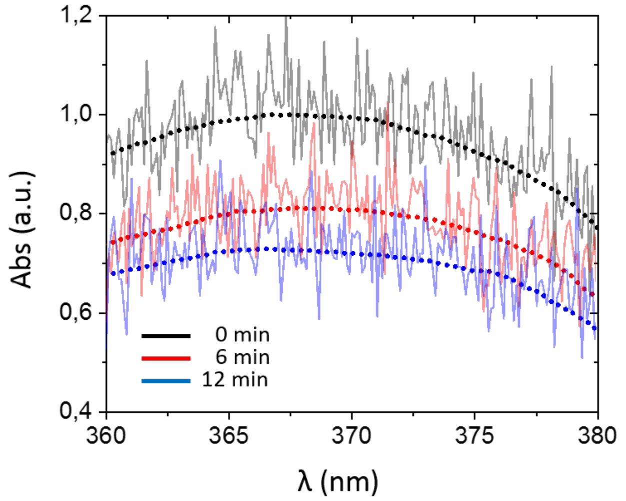

Figure S7: Absorbance spectra of **SBTub3** (starting from all-*E* sample at 0 min – black curve) after 6 min and 12 min 2PE (see also Fig 1e). Dotted lines represent the smoothed using a Savitzky-Golay filter and is a guide to the eye. The spectra show an absorbance maximum at ~367 nm, which matches that obtained from UV-Vis measurements of **SBTub3** in pure DMSO (see Fig S2).

### Part C: Biochemistry

#### *Stability towards glutathione – comparison to an azobenzene photoswitch*

Stability towards glutathione (GSH) of samples of both all-*E* (“dark”) and mostly-*Z* (“pre-lit”) **SBTub2** was determined similarly to published methods, and compared to that of **PST-1** (all-*E* “dark”, and mostly-*Z* “lit”).[10] All solutions were prepared in PBS pH ~7.4 with 10% DMSO, containing GSH (10 mM), in UV-Vis cuvettes that were sealed under air atmosphere (gas head volume <2 mL) with parafilm and maintained at 37°C, during absorbance measurements over several hours in a Varian Cary 60 spectrophotometer. “Dark” assays were performed with the *E*-isomers of the test compounds; “pre-lit” **SBTub2** (note: relaxation far slower than experiment time) had been pre-illuminated to reach PSS under 360 nm light; while “lit” **PST-1** (note: relaxation faster than experiment time) was maintained under continuous illumination from above with a 390 nm LED to maintain undegraded **PST-1** in a mostly-*cis* state. The test compounds’ π→π* transition absorbance (345 nm) was measured in one timecourse (Fig S8a-b); in a second experiment, the full absorbance spectra of the test samples or of a no-compound GSH-only control were recorded in time series (Fig S8c-e).

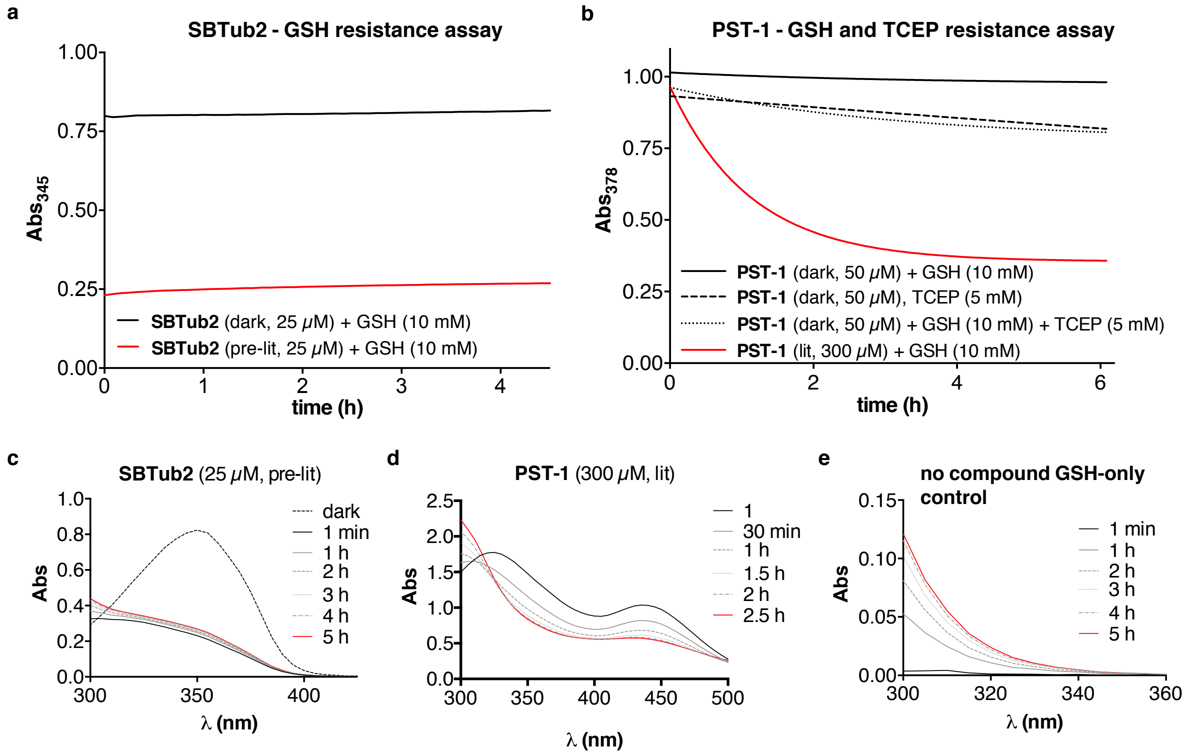

Figure S8: GSH resistance assays, incubating test compounds with 10 mM GSH + 10% DMSO. **a-b** π→π* transition absorbances of **SBTub2** and **PST-1** over time during GSH challenge. **c-e** absorption spectra of **SBTub2** (25 μM), **PST-1** (300 μM), and no-compound control sample (dark, only 10 mM GSH + 10% DMSO), during GSH challenge.

**SBTub2** has a large enough π→π* absorbance coefficient to be reliably measured at 25 μM in both dark and lit experiments; the smaller absorbance coefficient of *trans*-**PST-1**

encouraged us to use it at 50 *μ*M for the dark assay, while we performed the crucial lit assay of **PST-1** (*Z* isomer absorbance ca. 6 times lower than *E* at 345 nm) at 300 *μ*M.

**SBTub2** was stable against GSH addition and/or GSH-dependent reduction of the chromophore (which should both decrease its π→π* absorbance at 345 nm towards zero) in both all-*E* and mostly-*Z* assays. *Z*-**SBTub2** did not appear susceptible to significant *Z* to *E* isomerisation catalysed by reversible GSH addition-elimination since if this had continued to completion it would increase the π→π* absorbance for the “lit” solution by a factor of 3; we instead assign the slight absorbance increase (see Fig S8c) primarily to GSH oxidation to GSSG (see Fig S8e) with almost negligeable thermal relaxation of *Z*-**SBTub2** over the experimental time course.

We compared the SBT scaffold’s GSH stability to that of the widely used photopharmaceutical scaffold azobenzene, using similarly-decorated azobenzene **PST-1** as a comparison.

In our hands *E*-**PST-1** (50 *μ*M) was stable to the GSH challenge (Fig S8b) which does not match the "GSH/reductive susceptibility” described in the literature for this as well as other azobenzenes.[10] However, we trust our result to replicate the average cellular situation, and ascribe the discrepancy to literature use of co-reductants such as phosphines which we feel do not well predict the cellular situation. Those literature assays are typically performed with a powerful co-reductant such as TCEP, that is intended to keep GSH in its reduced form, with the non-explicit assumption that the phosphine should not perform Mitsunobu-type addition to the diazene or otherwise affect the reaction paths available to the photoswitch. When we tested for degradative effects due to 5 mM TCEP (with and without GSH) we found that TCEP alone caused appreciable *E*-**PST-1** degradation, and to an identical degree as the TCEP + GSH combination. This suggests that TCEP (not GSH) is the agent responsible for degradation of *E*-**PST-1** in such literature “GSH assays”, arguing more generally for caution in interpreting prior *in vitro* assessments of azobenzene “bioinstability” against cellular thiol concentrations if non-innocent co-reductants were employed. In our opinion, limiting the amount of oxygen available to a sample (e.g. capping a vial/cuvette under air with a negligible headspace volume) should be sufficient for running such assays reliably: since it seems to us that applying an approx. 10 mM concentration of GSH (i.e. at least 30-fold excess), should allow enough GSH to persist in the reduced state throughout an assay lifetime, despite potential oxidation from dissolved oxygen or from air in the headspace, such that large excesses (with respect to test compound) of additional reductants of a chemically different nature (especially those that are not physiological in nature or in concentration) can be productively avoided. Certainly, a less risky approach to test whether autoxidative GSSG formation precludes biochemically relevant compound reduction in the assay lifetime would be to vary the concentration and/or equivalents of GSH applied.

However, we observed unequivocally that *Z*-**PST-1** (300 μM) evolved rapidly in the absence of TCEP, indicating near-stoichiometric degradation by GSH (Fig S8b, S8d). We ruled out addition-elimination-catalysed *Z* to *E* isomerisation as the *major* mechanism of absorbance evolution, since complete *Z* to *E* isomerisation would increase the assay absorbance reading from the starting value of ca. 1 to ca. 6, whereas in fact the absorbance decreases to 0.40 at plateau[11]. We presume that GSH adduct formation, and/or reduction through to the hydrazine or still further to the anilines, is the major cause of loss of absorbance.

##### *Further exploration of glutathione-induced Z-azobenzene degradation*

To examine the degradation of *Z*-azobenzene by GSH further, a timecourse of HPLC analyses of a similar "lit" sample of **PST-1** (0.5 mM, lit at 395 nm) with 10 mM GSH in PBS + 10% DMSO was performed following our standard HPLC conditions (Fig S9). The results showed progressive formation of two major UV-active impurities, which it is tempting to assign as *ortho*-SG adducts on north and south aryl rings but which we were not able to observe in MS.

The greater susceptibility of *Z*-**PST-1** was to be expected from the literature-known faster kinetics of nucleophile addition to *Z*-azobenzenes, as well as their greater oxidising potential, as compared to the *E* isomers.[12] The stoichiometry and the limiting amount of reduction in the presence of physiological concentrations of thiol reductant can however be questioned; and we prefer not to conclude broadly about the final fate of azobenzenes in cells, nor about the completeness of adduct formation or reduction in general, since while the cell-free model gives a clear result of *Z*-azobenzene degradation with GSH, this is not equivalent to showing *cellular* degradation: (1) Cells are not homogenous aqueous cosolvent environments but have complex compartmentalisation effects controlling compound distribution as well as redox environments[13], e.g. that lipid-environment-bound azobenzenes could very reasonably be expected to be protected from reaction with cytosolic GSH; (2) cellular, embryonic and adult animal assays[14-16] have robustly shown photoreversible switching with **PST-1**; and this implies that cells maintain a cellularly-available pool of freely photoisomerisable, non-adduct-state *E*- and *Z*-**PST-1** at a basal concentration that is above the inhibitory threshold, hence that reduction is by no means complete in the cellular context and certainly does not impact the cellular utility of this cytosol-active azobenzene; (3) the survival of cells treated briefly with high *Z*-**PST-1** concentrations (see one-time-activation assay below) argues strongly against cellularly-relevant stoichiometric disturbance of the thiol pool or of thiol-based enzymes through poorly-reversible adduct formation and/or reduction through to the hydrazine and beyond; (4) our results do not exclude a non-innocent role for the dimethylsulfoxide cosolvent which is present at very high equivalence (or its potential contaminants) in modifying the **PST**s' apparent GSH stability. Addressing these issues is beyond the scope of this study into styrylbenzothiazoles and will be tackled elsewhere.

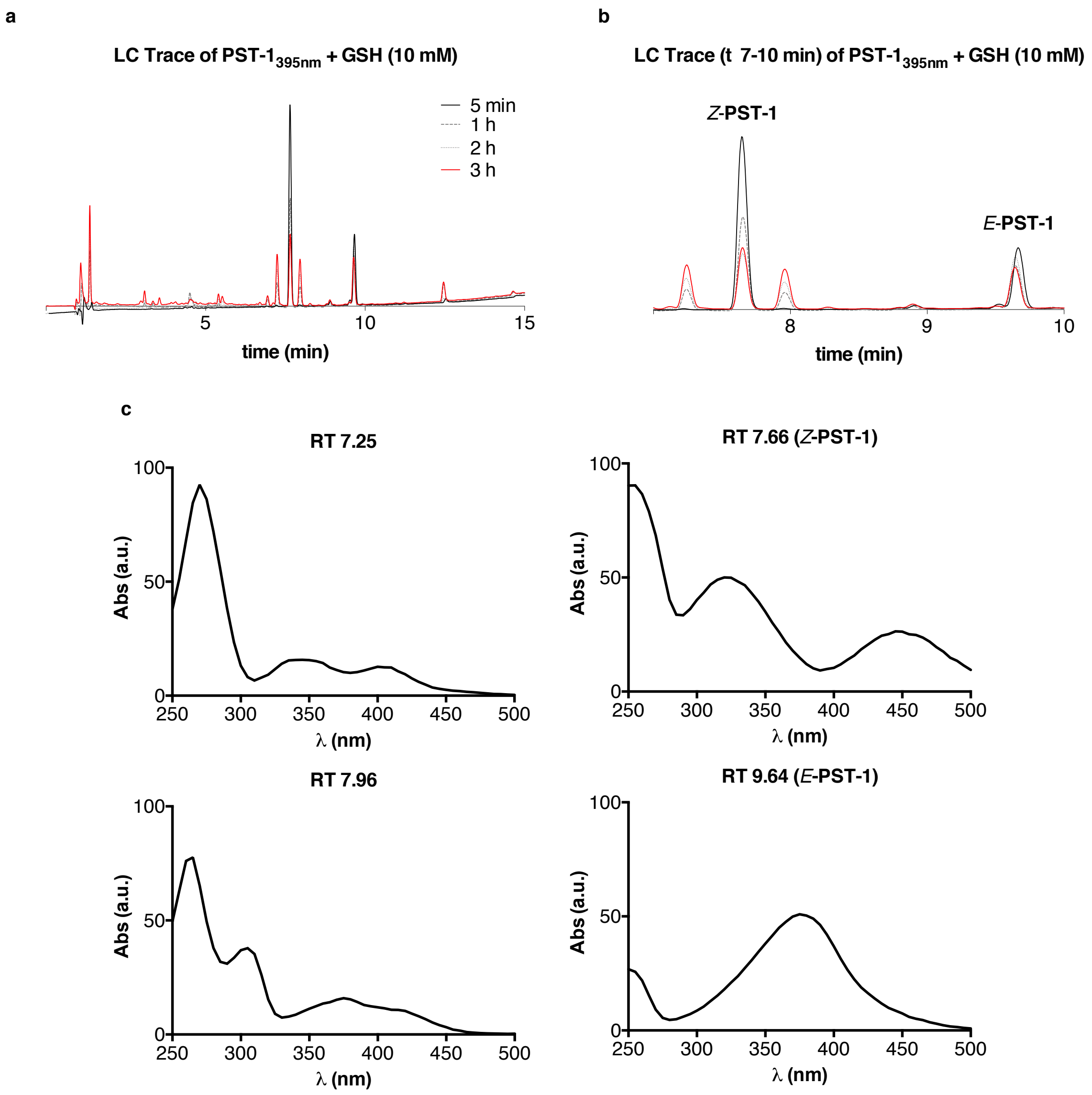

Figure S9: **a** 245 nm LC trace timecourse of **PST-1** (0.5 mM) incubated with 10 mM GSH. **b** Zoom in on panel a on the region of interest from 7–10 min showing *Z*-**PST-1** ($t_{ret}$ 7.66 min), *E*-**PST-1** ($t_{ret}$ 9.64 min) and two new signals increasing over time ($t_{ret}$ 7.25 min and $t_{ret}$ 7.96 min). **c** absorption spectra of the four major signals.

#### ***SBTubs show light specific inhibition of tubulin polymerisation*** **in vitro**

99% tubulin from porcine brain was obtained from Cytoskeleton Inc. (cat. #T240). The polymerisation reaction was performed at 5 mg/mL tubulin, in polymerisation buffer BRB80 (80 mM piperazine-N,N′-bis(2-ethanesulfonic acid) (PIPES) pH = 6.9; 0.5 mM EGTA; 2 mM $MgCl_2$), in a cuvette (120 µL final volume, 1 cm path length) in a Varian CaryScan 60 with Peltier cell temperature control unit maintained at 37°C; with glycerol (10 *µ*L). Tubulin was first incubated for 10 min at 37°C with “lit”- [360 nm-pre-illuminated; mostly-*Z*-] or dark- [all-*E*] **SBTub** (final **SBTub** concentration 20 *µ*M) in buffer with 3% DMSO, without GTP. Then GTP was added to achieve final GTP concentration 1 mM (with mixing), and the change in absorbance at 340 nm was monitored for 15 min, scanning at 15 s intervals[17]. While **SBTub3** in the “dark”-state (20 *µ*M) displayed almost similar MT growth dynamics as the cosolvent-only control, “lit” **SBTub3** (20 *µ*M) showed noticeable slowdown of polymerization kinetics, similar to known microtubule inhibitors such as colchicine (assayed at 16 *µ*M) (Fig S10).

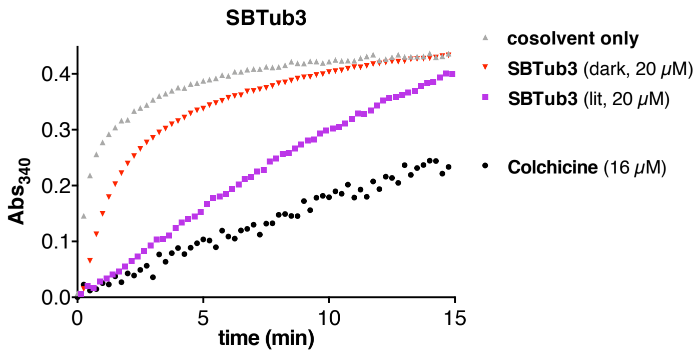

Figure S10: **SBTub3** inhibits tubulin polymerization after 360 nm illumination (violet squares) similar to the known tubulin inhibitor colchicine (grey circles), but in its dark state (black triangles) it shows near-identical behavior to the cosolvent (DMSO) control (gray triangles). Turbidimetric *in vitro* polymerization assay; greater absorbance corresponds to a greater degree of polymerization.

##### *CYP450 inhibition*

The potential for CYP450 inhibition was assessed by *in vitro* inhibition studies using fluorogenic CYP450 substrates with the corresponding CYP450 enzymes and NADPH regeneration system (Vivid CYP450 Screening Kits) with some minor changes to the manufacturer's protocols, performed by Bienta Biology Services (Fig S11a). The fluorescent signal produced from reaction is directly proportional to the cytochrome P450 activity. In the cases when tested compounds interfere with the CYP450 enzyme-substrate reaction, the fluorescent signal decreases.

In brief, the tested compounds were first dissolved in DMSO at 1 mM, then diluted in aqueous assay buffer to 25 µM, then the dilute solutions were mixed with a pre-mix consisting of human CYP450 + oxidoreductase and $NADP^+$ regeneration system (glucose-6-phosphate and glucose-6-phosphate dehydrogenase). After 10 min pre-incubation, the enzymatic reaction was initiated by the addition of a mix of NADPH and the appropriate CYP450 substrates yielding a test compound concentration of 10 *µ*M. The reaction was incubated for the desired reaction time (25 min for CYP1A2, CYP2C9, CYP2D6, and CYP3A4, 60 min for CYP2C19) after which Stop Reagent was added and fluorescence measured using SpectraMax Paradigm Multi-Mode Microplate Reader. All test points were performed in quadruplicates at concentration 10 µM (1% DMSO). Reference compounds used to benchmark the CYP inhibition under these conditions are listed in Fig S11b.

**a**

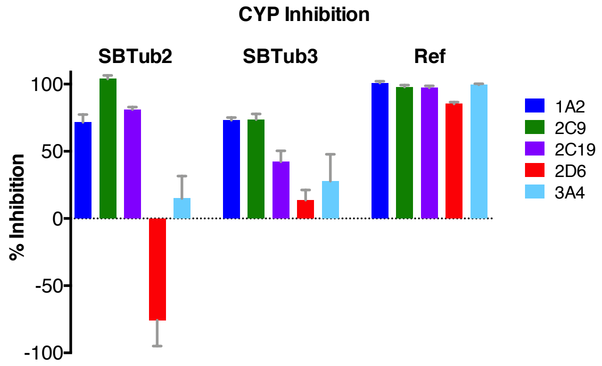

**b**

| CYP | Reference inhibitor | Ref. inhibitor conc., *µ*M | % Inhibition |
|---|---|---|---|
| 1A2 | α-naphthoflavone | 2 | 100.80 |
| 2C9 | sulfaphenazole | 5 | 97.86 |
| 2C19 | ticlopidine | 25 | 97.32 |
| 2D6 | quinidine | 0.5 | 85.23 |
| 3A4 | ketoconazole | 2 | 99.03 |

Figure S11: **a** summary of CYP450 inhibition assay by **SBTub**s at 10 *µ*M. **b** reference compounds used to compare CYP inhibition.

At 10 µM, **SBTub2** and **SBTub3** showed strong inhibition of 1A2, 2C9 and 2C19 CYP isoforms. *Higher* fluorescence was detected after incubation of **SBTub2** with CYP2D6 comparing to positive control, which resulted in negative value of inhibition, but we assign that result to assay interference. Further evaluations are ongoing.

#### *hERG Inhibition*

hERG inhibition experiments were performed by Bienta Biology Services using Invitrogen PredictorTM hERG Fluorescence Polarization Assay in accordance with the manufacturer's protocol (Protocol PV5365). In brief, a fluorescent tracer is incubated with **SBTub** and membranes bearing hERG channel for 2–4 hours in the solution and the polarization of fluorescence emission (higher when bound to hERG, lower in solution due to free tumbling) is assayed with reference to control inhibitor E-4031 to validate assay performance. **SBTubs** were tested in quadruplicates at 1 µM, 5 µM and 25 µM with reference to a positive control (only tracer, fluorescence polarization is maximal when nothing interferes with the reaction of the tracer and hERG membranes - minimal tracer rotation) and a negative control (30 µM of E-4031, i.e. 100% tracer displacement and minimum assay polarization value, due to tested compound competing with the tracer for the hERG channel, the polarization of emitted light) (Fig S12).

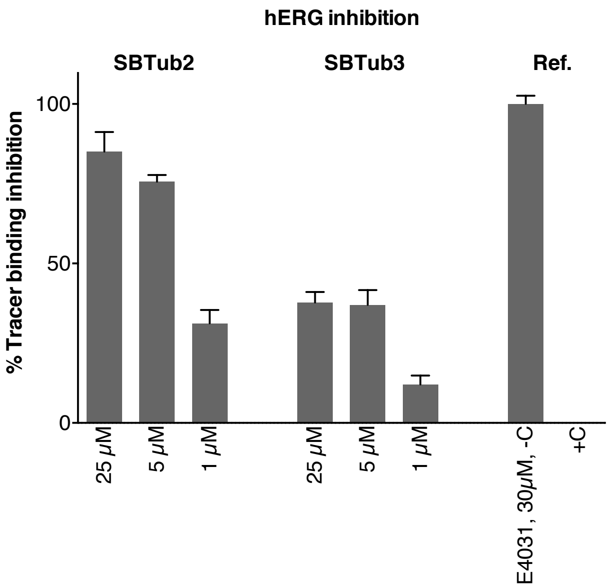

Figure S12: inhibition of tracer binding to hERG by **SBTub2** and **SBTub3** alongside negative control "-C" (30 µM of E4031) and positive control "+C" (cosolvent only). The data are represented as mean with SE, n=4.

**SBTub2** showed strong and dose dependent hERG binding, while **SBTub3** showed moderate and dose dependent hERG binding. However, tracer inhibition can be exaggerated if low compound solubility or if promiscuous protein binding are issues. Further evaluations in more sophisticated models are ongoing.

### Part D: Biological Data

#### *Cell assay methods*

##### General cell culture

HeLa cells were maintained under standard cell culture conditions in Dulbecco's modified Eagle's medium (DMEM; PAN-Biotech: P04-035550) supplemented with 10% fetal calf serum (FCS), 100 U/mL penicillin and 100 U/mL streptomycin. Cells were grown and incubated at 37°C in a 5% $CO_2$ atmosphere. Cells were typically transferred to phenol red free medium prior to assays (DMEM; PAN-Biotech: P04-03591). Substrates and cosolvent (DMSO; 1% final concentration) were added *via* a D300e digital dispenser (Tecan). Cells were either incubated under "lit" or "dark" conditions; "lit" indicates a pulsed illumination protocol was applied by multi-LED arrays to create the *Z*-isomers of the compounds in *situ* in cells and then maintain the wavelength-dependent PSS isomer ratio throughout the experiment, as described previously.[11] Typical "lit" timing conditions were 75 ms pulses applied every 15 s. "Dark" indicates that compounds were set to the all-*trans* state by thermal relaxation of the DMSO stocks at 60°C overnight, applied while working under red-light conditions, and cells were then incubated in light-proof boxes to shield from ambient light, thereby maintaining the all-*E*-isomer population throughout the experiment. *Single shot activation* indicates, that cells are treated with *E*-**SBTub**s and illuminated *in situ* for a short time (75 ms every 15 s during the first hour only; total illumination time 18 s) before stopping illuminations and incubating for the remaining 47 h without any further illumination.

##### MTT antiproliferation assay

Cells were seeded in 96-well plates at 5,000 cells/well and left to adhere for 24 h before treating with test compounds. *E*-**SBTub**s were added under the indicated lighting conditions for 48 h (final well volume 100 $\mu$L, 1% DMSO; six technical replicates); the "cosolvent control" ("ctrl") indicates treatment with DMSO only. Cells were then treated with 0.5 mg/mL (3-(4,5-dimethylthiazol-2-yl)-2,5-diphenyl tetrazolium bromide (**MTT**) for 3 h. The medium was aspirated and formazan crystals were re-dissolved in DMSO (100 $\mu$L); absorbance was measured at 550 nm ($Abs_{550}$) using a FLUOstar Omega microplate reader (BMG Labtech). Absorbance data was averaged over the technical replicates, then normalized to viable cell count from the cosolvent control cells (%control) as 100%, where 0% viability was assumed to correspond to absorbance zero. Three independent experiments were performed; data are shown from a single representative experiment, as indicated; data were plotted against the log of **SBTub** concentration ($\log_{10}$([**SBTub**]) (M)).

**Cell cycle analysis**

*E*-**SBTub**s were added to HeLa cells in 24-well plates (50,000 cells/well; three technical replicates, three biological replicates) and incubated under “dark” or “lit” conditions for 24 h. Cells were collected, permeabilised and stained with 2 μg/mL propidium iodide (PI) in HFS buffer (PBS, 0.1% Triton X-100, 0.1% sodium citrate) at 4°C for 30 min then analysed by flow cytometry using a FACS Canto II flow cytometer (Becton Dickinson) run by BD FACSDiva software. 30,000 events per technical replicate were analysed; the PI signal per event (corresponding to cellular DNA content) was measured and cells were binned into sub-G1, G1, S and G2 phase according to DNA content using *Flowing* software. Results (means of three technical replicates) from one experiment of three independent trials are shown.

**Immunofluorescence staining**

HeLa cells were seeded on glass coverslips in 24-well plates (50,000 cells/well) and treated with **SBTub**s the next day under “dark” or “lit” conditions for 24 h. Cells were washed with pre-warmed (37°C) MTSB buffer (80 mM PIPES pH 6.8; 1 mM $MgCl_2$, 5 mM ethylene glycol tetraacetic acid (EGTA) dipotassium salt; 0.5% Triton X-100) for 30 s then fixed with 0.5% glutaraldehyde for 10 min. After quenching with 0.1% $NaBH_4$ (7 min), samples were blocked with PBS + 10% FCS (30 min). The cells were treated with primary antibody (1:400 rabbit alpha-tubulin; Abcam ab18251) in PBS containing 10% FCS for 1 h and then washed with PBS. Cells were incubated with secondary antibody (1:400 goat-anti-rabbit Alexa fluor 488; Abcam ab150077) in PBS containing 10% FCS for 1 h. After washing with PBS, the coverslips were mounted onto glass slides using Roti-Mount FluorCare DAPI (Roth) and imaged with a Zeiss LSM Meta confocal microscope (CALM platform, LMU). Images were processed using the free Fiji software. Postprocessing was only performed to improve visibility. For maximum intensity projections, images were recorded at different focal planes incrementally stepping through the sample (step size 1–2 μm) and maximum intensity projections were obtained using Fiji software.

**EB3-comet live cell assays: general procedure**

HeLa cells (12,000 cells/well) were seeded on 8-well ibiTreat μ ibidi slides (ibidi, Martinsried) 24 h prior to transfection. Cells were transiently transfected with *EB3-GFP* or *EB3-YFP* plasmids using jetPRIME reagent (Polyplus) according to the manufacturer’s instructions. Cells were imaged 24 h later, at 37°C under 5% $CO_2$ atmosphere, using an UltraVIEW Vox spinning disc confocal microscope (PerkinElmer) operated with *Volocity* software equipped with an EMCCD camera (Hammamatsu, Japan), and an environmental chamber kept at 37°C and 5% $CO_2$ using a 63 × 1.4 NA Plan-Apochromat oil-immersion objective (Zeiss), while applying the specified illumination conditions. EB3-comets[18] were counted with a plugin for the *Fiji* software, based on the "Find maxima" function from the NIH (https://imagej.nih.gov/ij/macros/FindStackMaxima.txt).

**Live cell imaging for photobleaching/wavelength orthogonality**

For photobleaching/wavelength orthogonality experiments, a suitable focus plane was first chosen on the microscope (white light for focusing). **SBTub** was then added *cautiously* while the cells were still on the microscope stage, and cells incubated for 10 min before imaging. This protocol avoids exposure of the **SBTub** to white focusing light, preventing unwanted isomerization prior to imaging which could falsify results when testing for GFP/YFP orthogonality. Cells were imaged either at 488 nm (GFP; 23% laser power, 400 ms exposure time, 45 frames/min) or 514 nm (YFP; similar parameters; data not shown since GFP orthogonality implies YFP orthogonality); cells were optionally additionally exposed to interleaved dummy frames of **SBTub**-isomerizing 405 nm light for compound activation (250 ms exposure time, 45 frames/min). For analysis statistics, 6 cells per condition from three independent trials were taken. First-order exponential decay curves were fitted to each cell's data with their comet counts normalized to 100% at time zero, enabling intercomparison of cells with different starting comet counts (depending on their size, the position of the focal plane, etc).

**Treatment of live cells with *Z*-SBTub3 under GFP imaging**

To show the effect of photoisomerized *Z*-**SBTub3** (as a control for the orthogonality experiment), a suitable focus plane was first chosen on the microscope (white light for focusing) and 5 frames were acquired at 488 nm, to set a reference for basal comet count. An **SBTub3** stock that had been isomerized to PSS with 360 nm was carefully applied on the stage, and after one minute incubation time, 180 frames were taken at 488 nm (400 ms exposure time, 23% laser power, 60 frames/min). The number of remaining comets per frame after treatment was counted and expressed as percentage of the starting reference timepoint.

**Temporally reversible MT dynamics modulation with GFP-orthogonality**

HeLa cells were transfected with EB3-GFP using FuGENE 6 (Promega) according to manufacturer's instructions. Experiments were imaged on a Nikon Eclipse Ti microscope equipped with a perfect focus system (Nikon), a spinning disk-based confocal scanner unit (CSU-X1-A1, Yokogawa), an Evolve 512 EMCCD camera (Photometrics) attached to a 2.0× intermediate lens (Edmund Optics), a Roper Scientific custom-made set of Stradus 405 nm (100 mW, Vortran) and Calypso 491 nm (100 mW, Cobolt) lasers, a set of ET-BFP2 and ET-GFP filters (Chroma), a motorized stage MS-2000-XYZ, a stage top incubator INUBG2E-

ZILCS (Tokai Hit) and lens heating calibrated for incubation at 37°C with 5% $CO_2$. Microscope image acquisition was controlled using MetaMorph 7.7 and images were acquired using a Plan Apo VC 60× NA 1.4 oil objective. Comet count analysis was performed in ImageJ using the ComDet plugin (E. Katrukha, University of Utrecht, Netherlands, https://github.com/ekatrukha/ComDet). GFP imaging was performed at 491 nm (0.17 mW; 300 ms every 2 s). In-frame **SBTub** photoactivation was performed at 405 nm (77 $\mu$W; 100 ms every 90 s; cell of interest plus in-frame surrounding area). Results were presented in the main text and correspond to Movie M1.

#### *FACS cell cycle analysis*

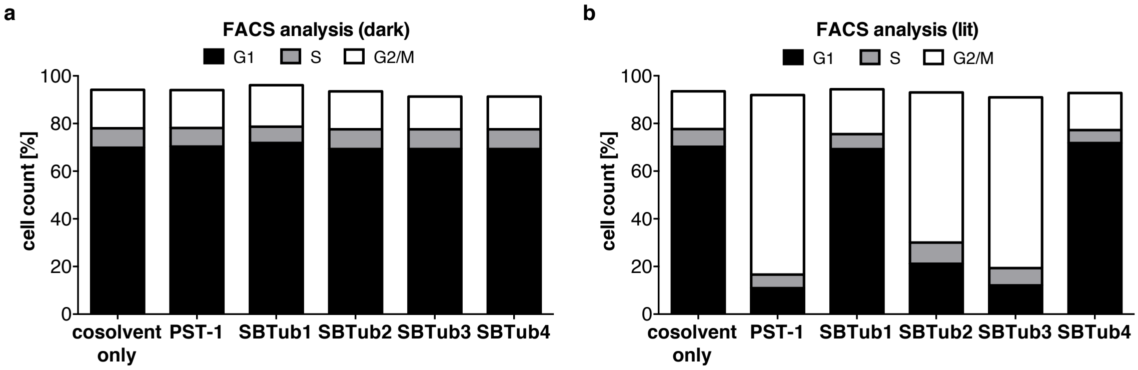

Figure S13: Cell cycle analysis of **a** “dark” and **b** “lit” states of **SBTub1-4** (20 $\mu$M) in comparison to positive photoswitchable control **PST-1** (2.5 $\mu$M) and negative cosolvent-only control. Note: panel **b** is identical to Fig 3d.

Results show full photoswitch-based control of G2/M-phase cell cycle arresting properties of **SBTub2** and **SBTub3**, with no cell cycle effects from either isomer of **SBTub1** or **SBTub4**.

#### *Immunofluorescence imaging of microtubule network structure*

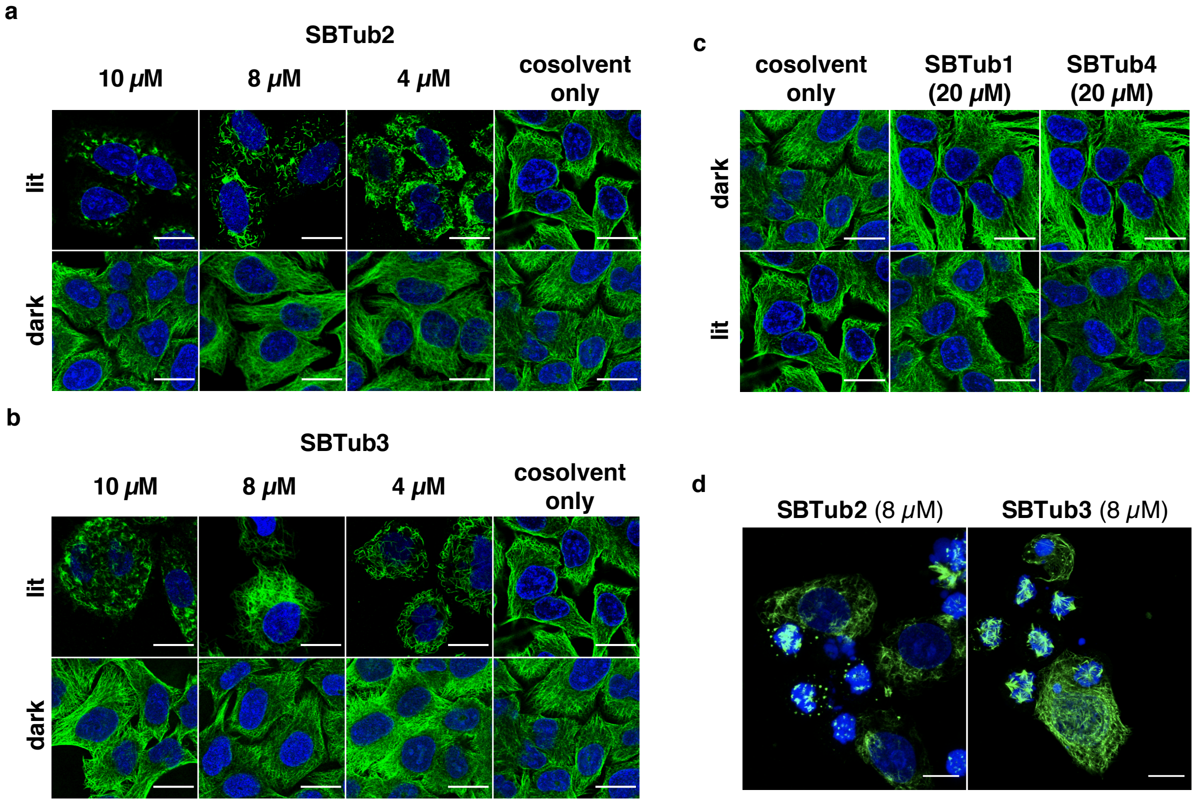

Figure S14: Immunofluorescence imaging of cells treated under lit/dark conditions with **a SBTub2** or **b SBTub3**, or with permutation controls **c SBTub1** and **SBTub4**. **d** Maximum intensity projection (along z-axis) of cells treated for 24 h with lit **SBTub2** or lit **SBTub3**; expanded field of view from data shown in Fig 3d (Hela cells, MTs immunostained with anti-$\alpha$-tubulin (green), nuclei stained with DAPI (blue), scale bars = 20 μm).

The results in Fig S14a-c show light- and dose- dependent depolymerisation of microtubule structure by **SBTub2** and **SBTub3**, while **SBTub1** and **SBTub4** have no effect on MT structure under dark and illuminated conditions even at high concentrations of 20 μM. The maximum intensity projection (along the z-axis) of cells treated for 24 h with lit **SBTub2** or lit **SBTub3** (Fig S14d) shows that most treated cells have been arrested in mitosis and display severe MT depolymerisation and organisational defects, while only a minor population of adherant cells persists albeit with disorganized microtubule networks.

#### *Live cell imaging: GFP/YFP orthogonality via EB3 comet assay*

Two assays were used, such that taken together, the results show orthogonality of both *E*- and *Z*-**SBTub**s to GFP/YFP imaging.

Firstly: stability of the *Z*-**SBTub** against *Z*-to-*E* isomerisation under GFP/YFP imaging was assessed. As per the "live cell imaging" methods section, cells were pre-incubated with all-*E* **SBTub3** (10 μM) or with cosolvent only (control) in the dark for 10 min before acquiring uninhibited cellular EB3 comet count statistics at timepoint $t_{ref}$. **SBTub3** was then isomerized substantially to *Z* by whole-field illumination at 405 nm (5 × 300 ms) and 5 GFP/YFP imaging frames at time points 30 s, 5 min and 10 min later were acquired (Fig S15a). Three independent trials were performed. For analysis, 8 cells per condition were chosen from the three independent trials. The EB3-comets were counted and each time point's comet count value was set as the average from those of its 5 frames. The average comet count before illumination (before isomerisation of **SBTub** or photobleaching of the marker) was set to 100%. The number of comets at the following timepoints are expressed as percentage of the initial pre-treatment comet numbers and are represented as mean with SD (Fig S15b-c).

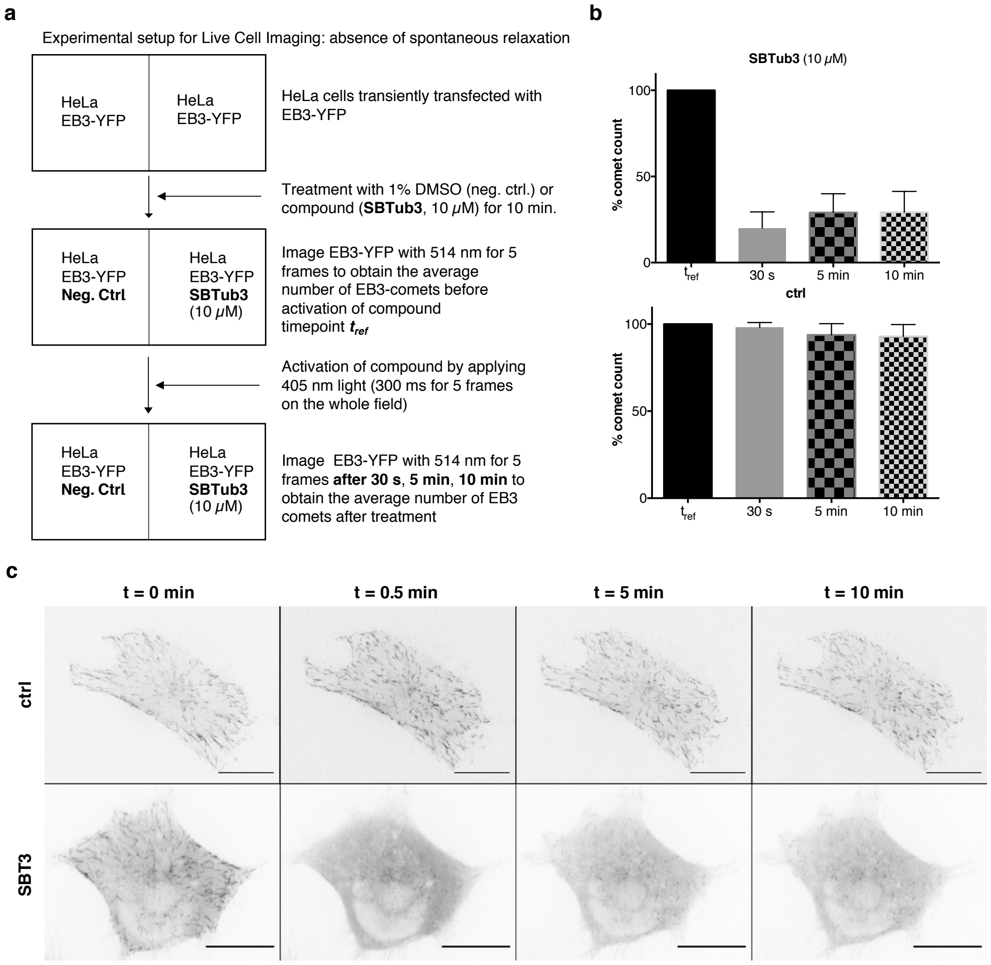

Figure S15: **a** Experimental setup for photoswitching during live cell imaging of **SBTub3** to evaluate orthogonality to YFP imaging (similar protocol for GFP). **b** Quantification of EB3-YFP comets at different timepoints upon photoswitching during live cell imaging, normalised relative to comet count at $t_{ref}$. **c** Stills of **SBTub3** live cell imaging experiment corresponding to Fig S15b (color scheme inverted to highlight EB3 comets, which here appear as black dots); scale bar represents 20 μm.

Comet counts were abruptly and, in this experiment permanently reduced, upon the first global **SBTub3** activation with 405 nm while the cosolvent control showed no effect from the photoswitching procedure, indicating the photostability of the bioactive *Z*-**SBTub3**.

Secondly: evaluation of *E*-**SBTub**s' photostability ability to effect *in situ* photocontrol over protein dynamics while avoiding interference from the imaging wavelengths of common fluorescent labels GFP and YFP was performed by comparing the EB3 comet count under different lighting and compound conditions (Fig S16).

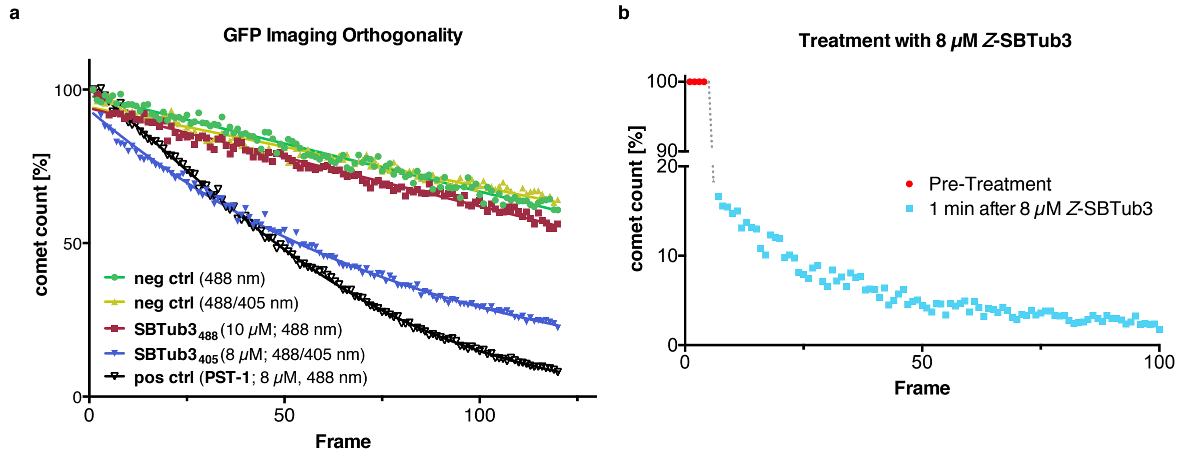

Figure S16: EB3 comet count plotted against frames **a** under different imaging and compound conditions **b** upon treatmend with pre-lit **SBTub3**. Note that panel **a** is identical to Fig 4b in the main text.

Fig S16a shows the following conditions and conclusions:

**neg ctrl**: cosolvent-only controls (DMSO) imaged under 488 nm only (green circles); or under alternating 488/405 nm (yellow triangles) lighting protocol as used for **SBTub** photoactivation. Shows the timecourse of “normal” reduction of EB3 comets due to photobleaching only.

**SBTub3$_{488}$** (red squares): cells treated with **SBTub3** (10 μM) and imaged at 488 nm only: reduction of EB3 comets was comparable to those of both **neg ctrl**s, indicating that **SBTub3** is not photoactivated by 488 nm light.

**SBTub3$_{405}$** (blue triangles): cells treated with **SBTub3** (8 μM) and imaged with 488 nm frames with interleaved 405 nm frames to induce **SBTub** activation. These show a faster decrease of EB3 comet count than of both **neg ctrl**s and of **SBTub3$_{488}$**; decrease is comparable to that of **pos ctrl PST-1** imaged at 488 nm, suggesting comparable inhibition of MT polymerisation.

**pos ctrl:** known azobenzene based MT depolymerizer **PST-1** (8 μM, non-GFP orthogonal) was imaged at 488 nm, causing compound activation and fast reduction of EB3 comet count (beyond the photobleaching rate).

Additionally, cells were treated with pre-illuminated *Z*-**SBTub3** enriched stock solutions and incubated for 1 min before imaging at 488 nm (Fig S16b). This experimental setup rules out a reduction of EB3 comets due to photobleaching, while also showing that reduction of EB3 comets is caused by adding *Z*-**SBTub3**.

Taken together, these results show the full orthogonality of both *E*- and *Z*-**SBTub**s to GFP/YFP imaging.

### Part E: Protein Crystallization

#### *Protein crystallisation materials and methods*

##### Protein production, crystallization and soaking.

The DARPin D1 was prepared as previously described[19]. Tubulin from bovine brain was purchased from the Centro de Investigaciones Biológicas (Microtubule Stabilizing Agents Group), CSIC, Madrid, Spain.

The tubulin-DARPin D1 (TD1) complex was formed by mixing the respective components in a 1:1.1 molar ratio. The TD1 complex was crystallized overnight by the hanging drop vapor diffusion method (drop size 2 µL, drop ratio 1:1) at a concentration of 15.9 mg/mL and at 20°C with a precipitant solution containing 18% PEG 3350, 0.2 M ammonium sulfate and 0.1 M bis-tris methane, pH 5.5. All drops were subsequently hair-seeded with crystalline material obtained in previous PEG-screening, which resulted in single and large (0.5 µm) TD1 complex crystals. The crystals were fished and transferred into new precipitant solution drops containing 10% DMSO and respective compounds (**SBTub2**/**SBTub3**) at a final concentration of 2 mM and were pre-switched with 360 nm LEDs for 5 min. After 4 h of soaking in the dark, the **SBTub:**tubulin crystals were mounted for X-ray diffraction data collection.

##### Data Collection, processing and refinement.

Data were collected at beamline X06SA at the Swiss Light Source (Paul Scherrer Institute, Villigen, Switzerland). The beam was focused to 30 x 30 µm, the flux was 3 x $10^{10}$ photons/ s and the data collection speed was 2°/s at an oscillation range of 0.2° per frame (see Table 1). For TD1-**SBTub2** and TD1-**SBTub3**, 210° and 220° of data were collected, respectively. Data processing was done with XDS[20]. Due to anisotropy, the data were corrected using the Staraniso server (http://staraniso.globalphasing.org/). The structures were solved by molecular replacement using PDB ID 5NQT as a search model[21]. The ligands and restraints were generated with the grade server (http://grade.globalphasing.org/) using their SMILES annotation. The structures were then refined iteratively in PHENIX[22] with manual editing cycles in COOT[23].

##### *X-ray structural analysis*

| **Data Statistics** | **TD1-SBTub2** | **TD1-SBTub3** |
|---|---|---|
| Space group | P $2_1$ | P $2_1$ |
| Unit cell (a; b; c; β) | 73.90 91.85 82.81 96.58 | 73.87 91.72 83.00 97.06 |
| Wavelength (Å) | 1.0 | 1.0 |
| Resolution (Å) | 45.18 – 2.05 (2.33 – 2.05) | 45.85 – 1.86 |
| $R_{pim}$ (%) | 7.4 (59.1) | 5.5 (50.0) |
| I/σI | 7.1 (1.6) | 9.3 (1.7) |
| Spherical completeness (%) | 59.5 (9.3) | 68.4 (12.9) |
| Ellipsoidal completeness (%) | 88.2 (53.1) | 91.2 (59.9) |
| Ellipsoidal truncation resolution limits (Å) | 2.04, 2.50, 2.55 | 1.84, 2.23, 2,12 |
| Multiplicity | 4.1 (4.7) | 4.2 (4.3) |
| $CC_{1/2}$ | 0.992 (0.951) | 0.997 (0.526) |
| **Refinement Statistics** | | |
| Resolution | 45.18 – 2.05 | 45.04 – 1.75 |
| No. reflections | 40926 | 68146 |
| $R_{work}$ / $R_{free}$ | 22.5 / 28.1 | 18.5 / 23.3 |
| Ramachandran favoured | 94.98 % | 97.15 |
| Ramachandran outliers | 0.79 % | 0.1% |
| R.m.s.d. bond length (Å) | 0.002 | 0.003 |
| R.m.s.d. bond angles (°) | 0.463 | 0.627 |
| PDB code | XXX (to be added) | XXX (to be added) |

Table S1 Crystallographic Data for **SBTub2** and **SBTub3**

Both ligands, **SBTub2** and **SBTub3**, bind to the colchicine site at the interface of α- and β-tubulin (Fig S17-S22). The two ligands bind in exactly opposite poses despite interacting with identical residues in the binding pocket, according to the design goals of this study, whereby the poses should be defined by the specific part of the ligand inherited from the combretastatin-A4 archetype (trimethoxyphenyl in case of **SBTub2** and isovanillinyl in case of **SBTub3**). The benzothiazole group is able to bind in both orientations (Fig S20).[6] The β-T7 loop is not well ordered in both tubulin-**SBTub2** and tubulin-**SBTub3** structures, likely due to the influence of the ligands. Despite the weak density for some parts of the loop, it is apparent that the larger trimethoxyphenyl-ring of **SBTub2** occupies more space and pushes β-Leu-248 and the β-T7 loop further out than the smaller flat benzothiazole group in **SBTub3** (Fig S21-22).

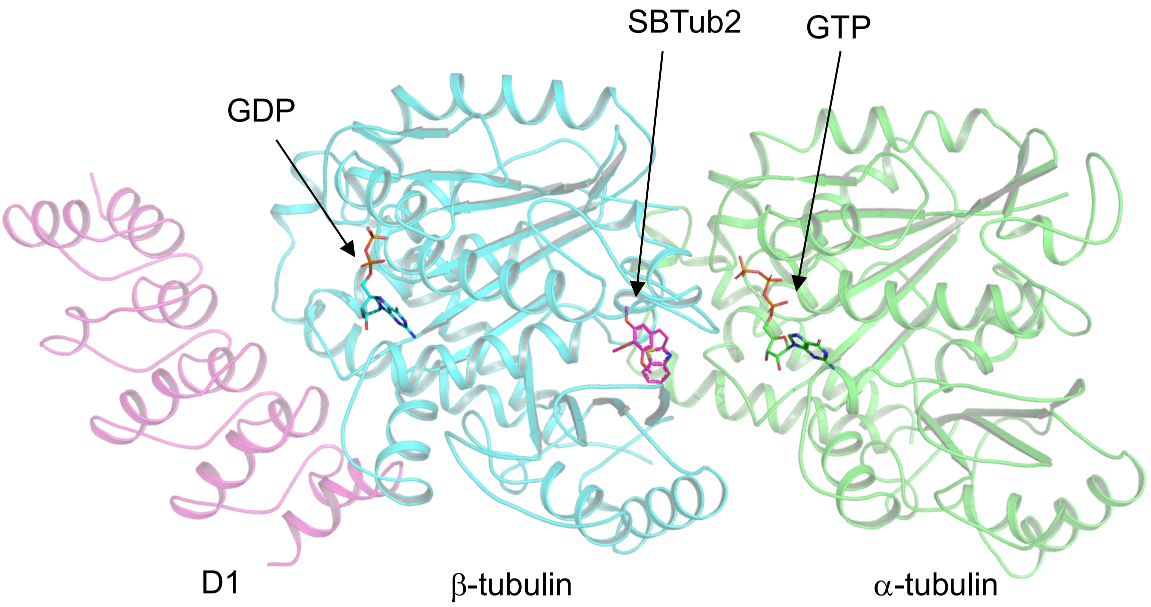

Figure S17: Overall view of the TD1-**SBTub2** complex structure shown in cartoon representation with the different ligands shown in sticks representation.

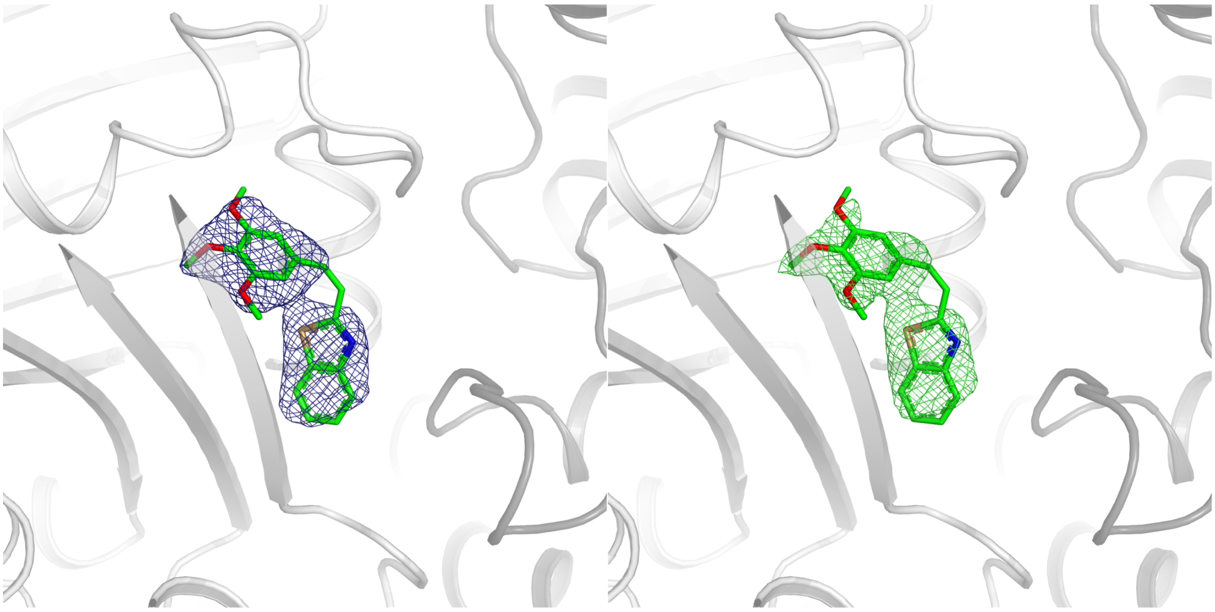

Figure S18: Colchicine site of tubulin-**SBTub2** with 2FoFc map contoured at 1σ shown in blue (left) and simulated annealing omit map at 2.5σ shown in green (right) mesh representation.

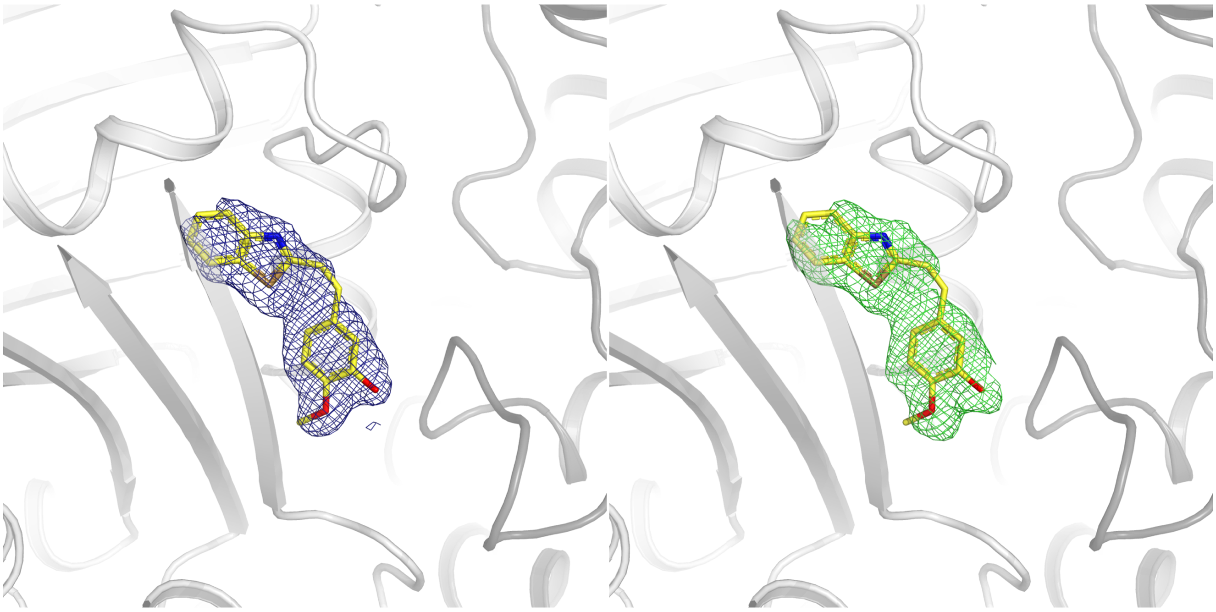

Figure S19: Colchicine site of tubulin-**SBTub3** with 2FoFc map contoured at 1σ shown in blue (left) and simulated annealing omit map at 2.5σ shown in green (right) mesh representation.

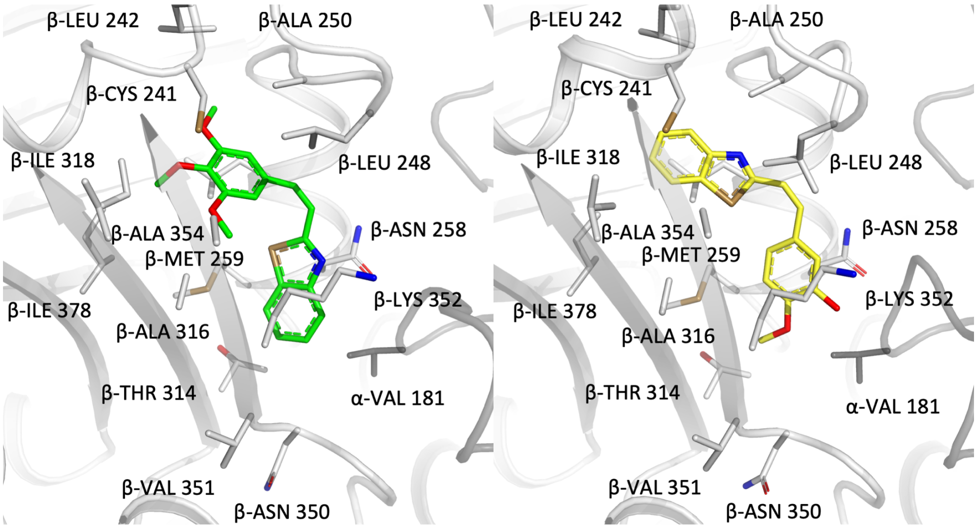

Figure S20: Close-up views of the colchicine sites of the tubulin-**SBTub2** (left) and tubulin-**SBTub3** (right) complex structures. Tubulin is shown in cartoon representation (dark grey for α-tubulin and light grey for β-tubulin); the ligands and interacting residues are shown in sticks representation. Oxygen and nitrogen atoms are colored red and blue, respectively; carbon atoms are colored in green (**SBTub2**), yellow (**SBTub3**), dark grey (α-tubulin) or light grey (β-tubulin).

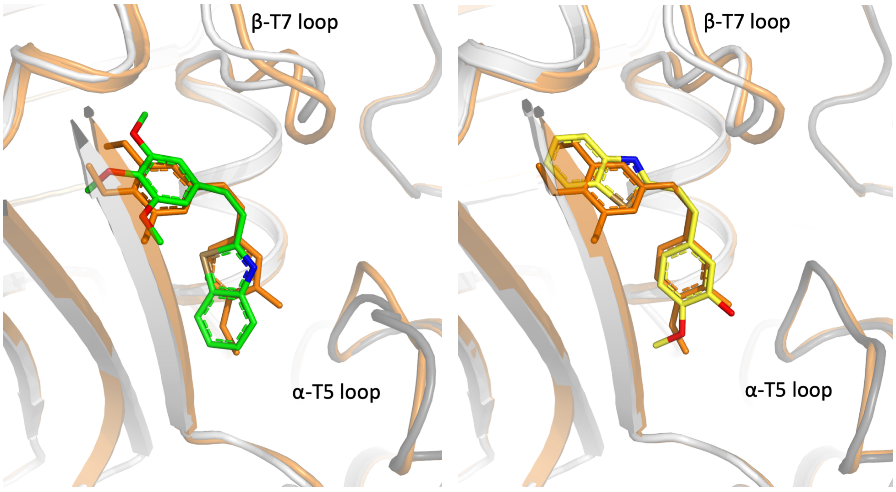

Figure S21: Superimpositions of the colchicine sites of tubulin-**SBTub2** (left, green carbons) and tubulin-**SBTub3** (right, yellow carbons) onto the tubulin-combretastatin A4 (orange carbons; PDB ID 5LYJ) complex structures.

The hydrophilic OH-group at the isovanillyl-ring of **SBTub3** (which binds in the "top-down" pose) appears to stabilize the α-T5 loop in the conformation the ligand-free structure displays (Fig S21). **SBTub2** binds in the "bottom-up" pose and the α-T5 loop displays an altered conformation similar to that seen with colchicine (Fig S22). While both ligands, being colchicine site binders, likely interfere with the "curved-to-straight" conformational transition of tubulin during the formation of microtubules[24,25], their difference in disturbing the α-T5 loop conformation may be of pharmacological relevance, given that the archetypical colchicine disturbs the α-T5 loop, while the vascular disrupting agent combretastatin A4 does not.[6]

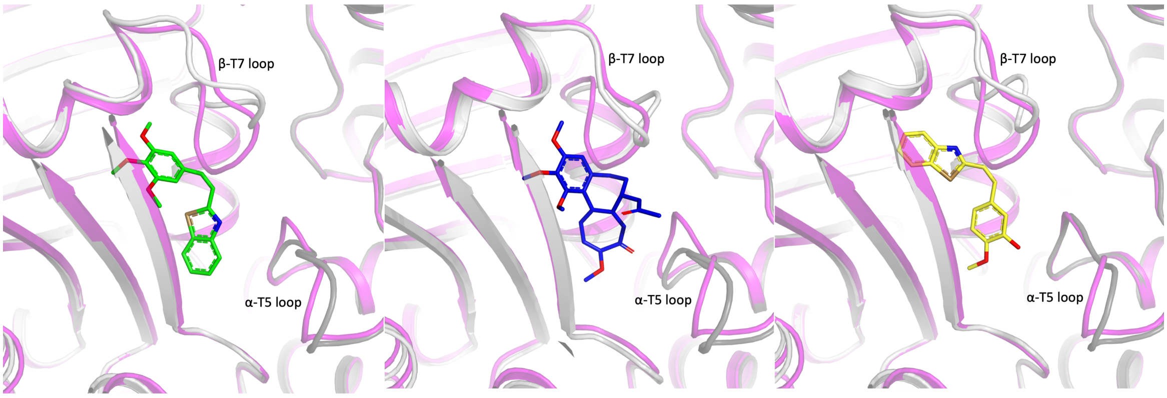

Figure S22: Side-by-side comparison of the colchicine sites of the tubulin-colchicine (mid; PDB ID 5NM5), tubulin-**SBTub2** (left) and tubulin-**SBTub3** (right) complex structures. On each structure, the ligand free structure (PDB ID 5NQT) has been superimposed (shown in pink).

### Part F: NMR Spectra

#### 2-(2,3,4-trimethoxystyryl)benzothiazole (SBTub1)

##### [1]H-NMR:

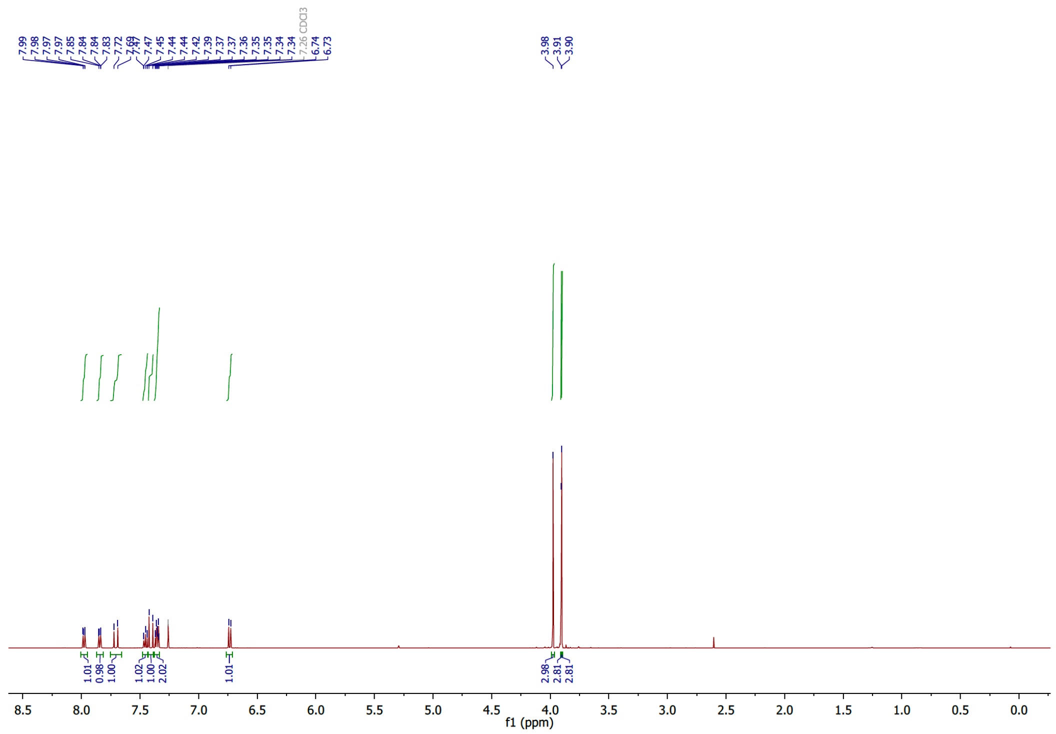

##### [13]C-NMR:

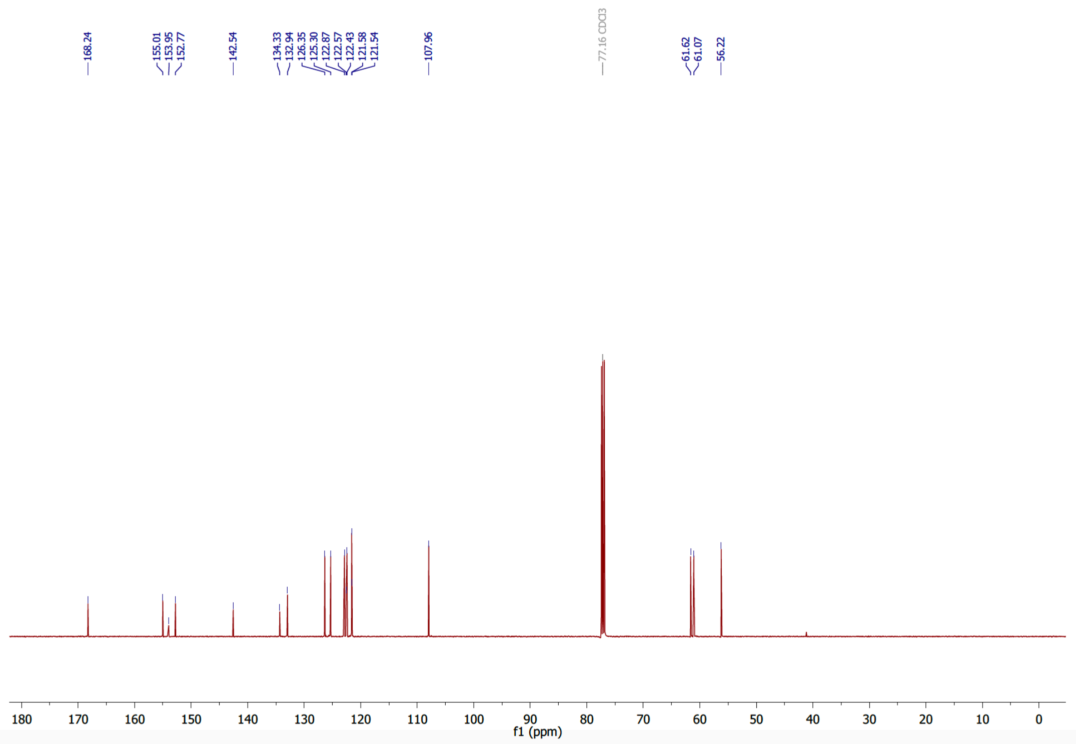

#### 2-(3,4,5-trimethoxystyryl)benzothiazole (SBTub2)

##### $^1$H-NMR:

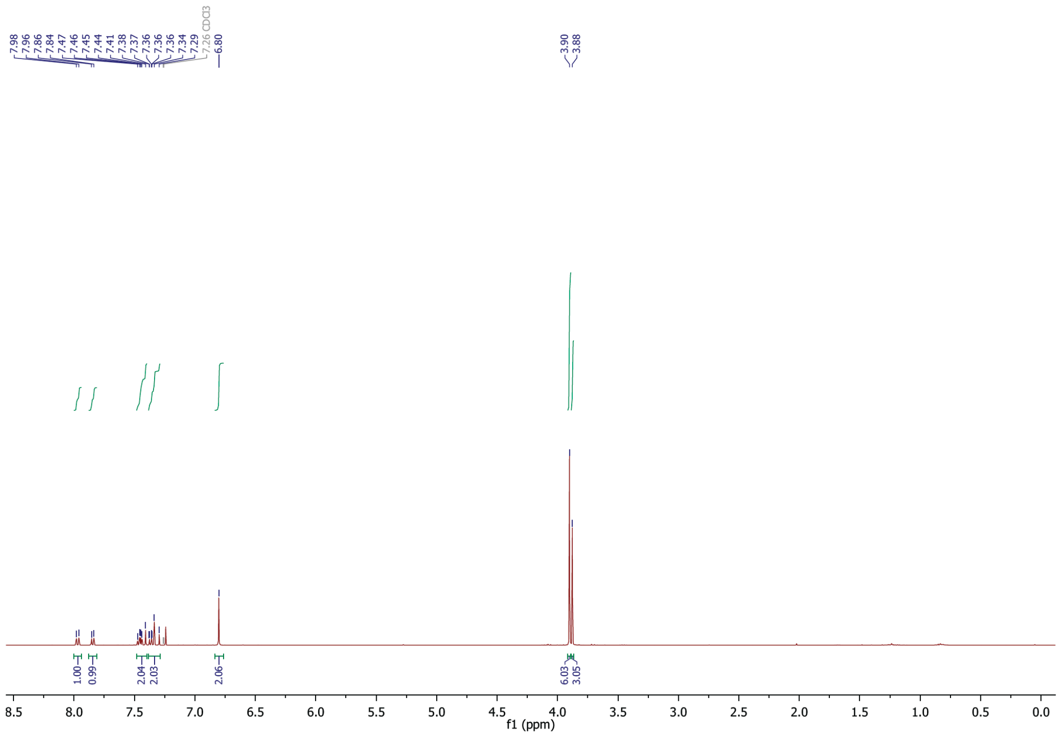

##### $^{13}$C-NMR:

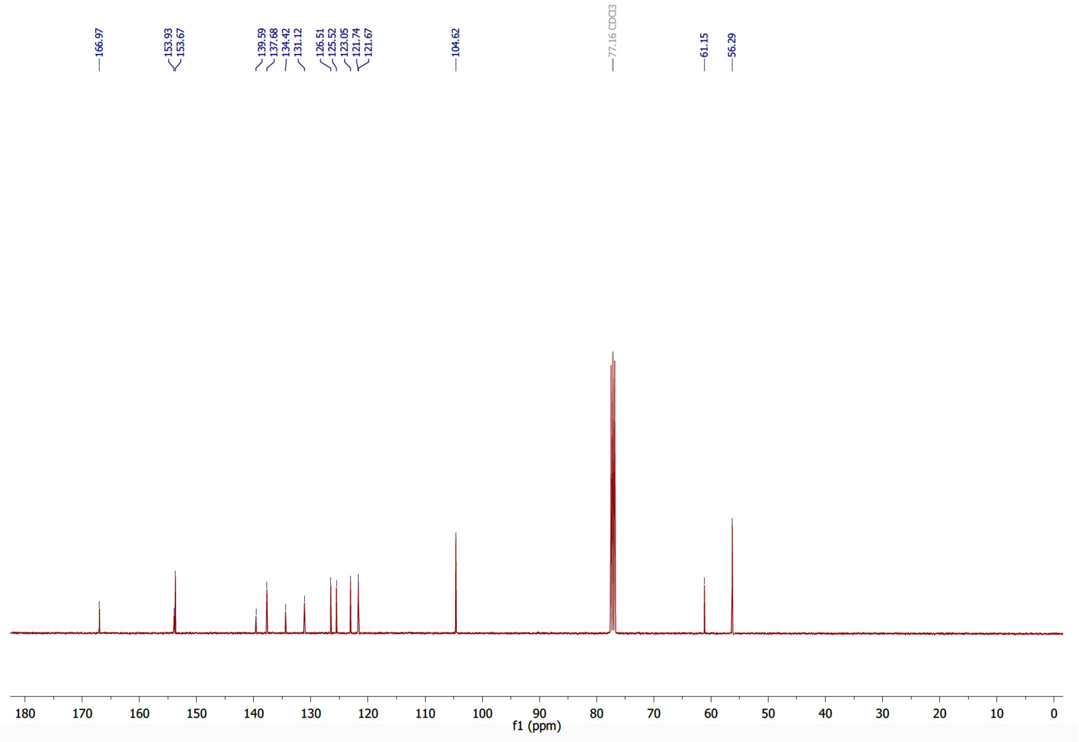

**5-(2-(benzothiazol-2-yl)vinyl)-2-methoxyphenol (SBTub3)**

**¹H-NMR:**

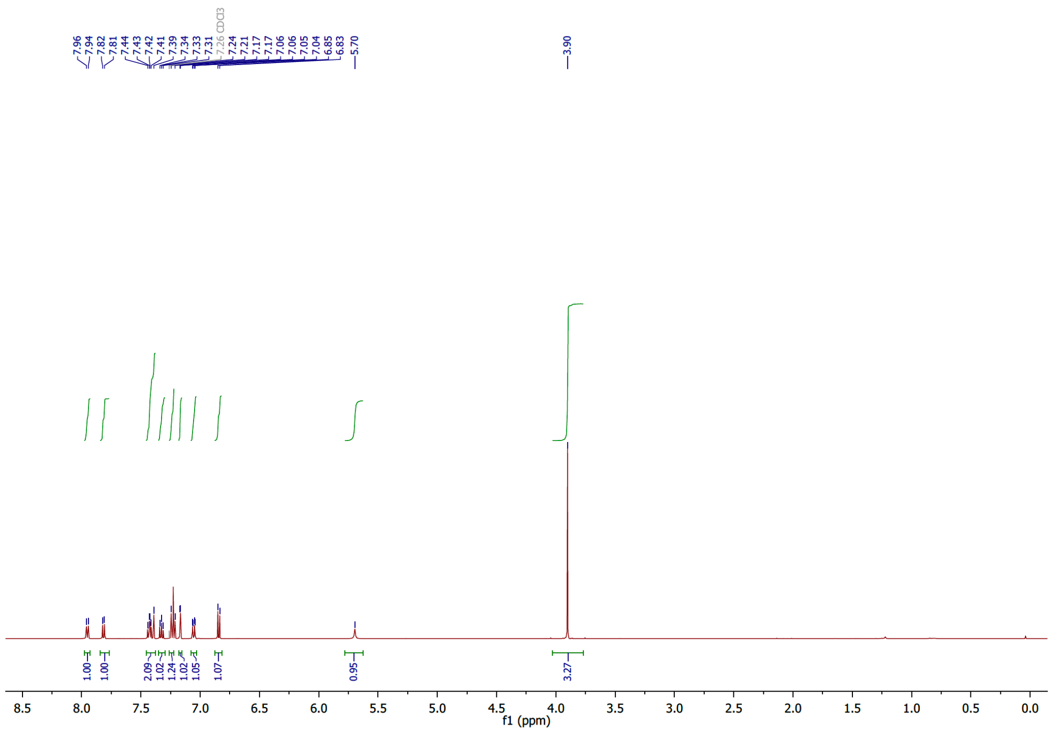

**¹³C-NMR:**

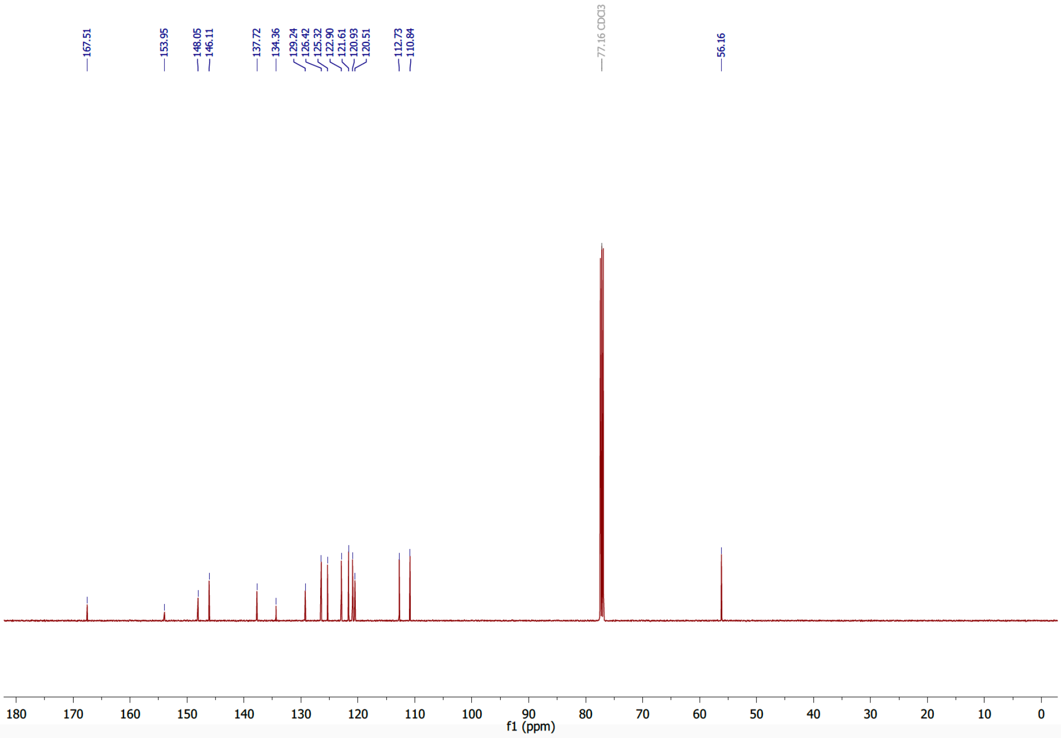

#### 4-(2-(benzothiazol-2-yl)vinyl)-2-methoxyphenol (SBTub4)

##### $^{1}$H-NMR:

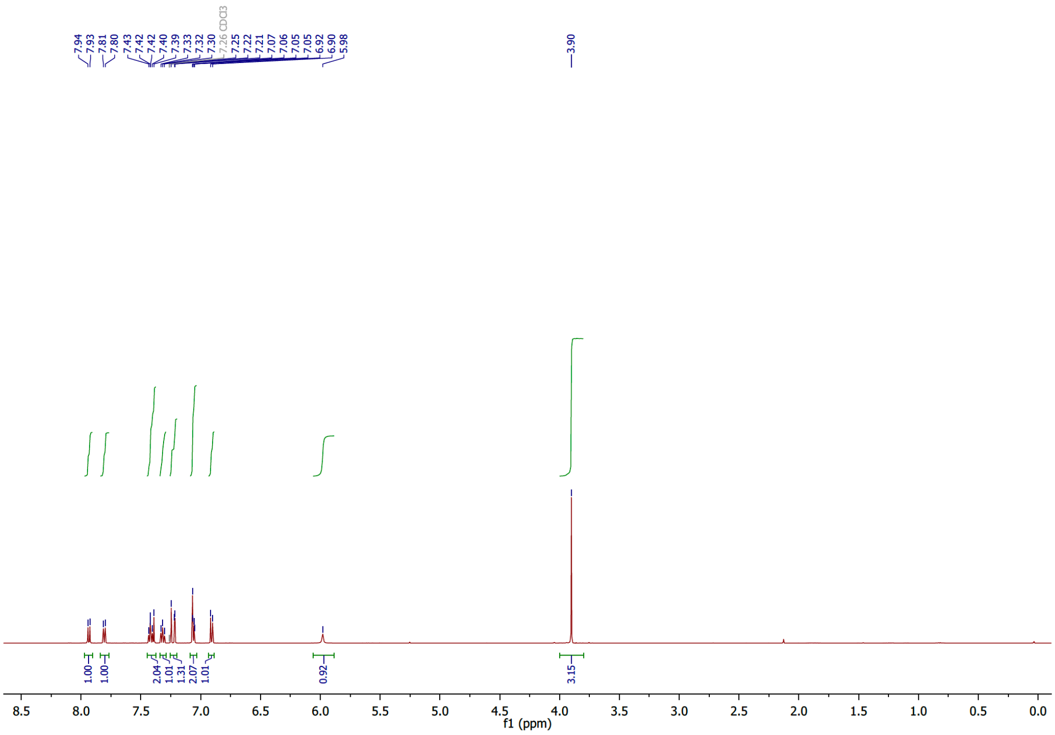

##### $^{13}$C-NMR:

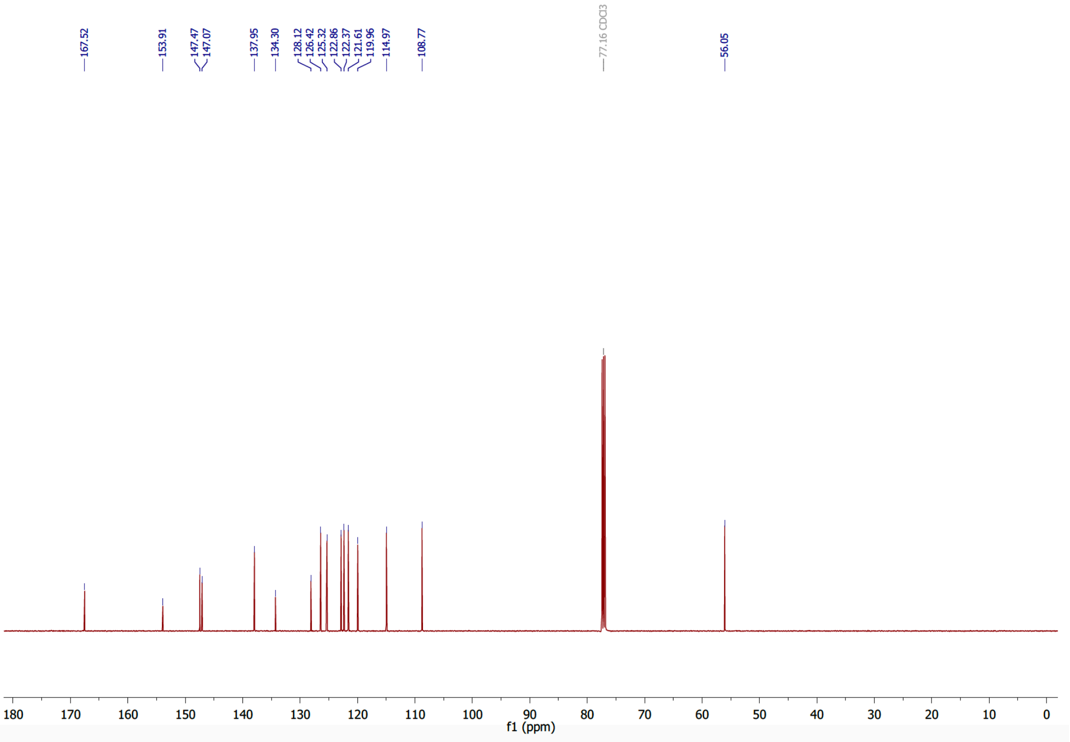

### Supplementary Information Bibliography

1 Gottlieb, H. E., Kotlyar, V. & Nudelman, A. NMR Chemical Shifts of Common Laboratory Solvents as Trace Impurities. *J. Org. Chem.* **62**, 7512-7515, doi:10.1021/jo971176v (1997).

2 Penthala, N. R., Crooks, P. & Sonar, V. Combretastatin analogs. US20160068506A1 (2016).

3 Penthala, N. R., Thakkar, S. & Crooks, P. A. Heteroaromatic analogs of the resveratrol analog DMU-212 as potent anti-cancer agents. *Bioorg. Med. Chem. Lett.* **25**, 2763-2767, doi:10.1016/j.bmcl.2015.05.019 (2015).

4 Tron, G. C. *et al.* Medicinal Chemistry of Combretastatin A4: Present and Future Directions. *J. Med. Chem.* **49**, 3033-3044, doi:10.1021/jm0512903 (2006).

5 Woods, J. A., Hadfield, J. A., Pettit, G. R., Fox, B. W. & McGown, A. T. The interaction with tubulin of a series of stilbenes based on combretastatin A-4. *Br. J. Cancer* **71**, 705-711, doi:10.1038/bjc.1995.138 (1995).

6 Gaspari, R., Prota, A. E., Bargsten, K., Cavalli, A. & Steinmetz, M. O. Structural Basis of *cis*- and *trans*-Combretastatin Binding to Tubulin. *Chem* **2**, 102-113, doi:10.1016/j.chempr.2016.12.005 (2017).

7 Lozano, C. M. *et al.* Cytotoxic anionic tribromo platinum(II) complexes containing benzothiazole and benzoxazole donors: synthesis, characterization, and structure-activity correlation. *Inorganica Chimica Acta* **271**, 137-144, doi:doi.org/10.1016/S0020-1693(97)05952-5 (1998).

8 Sanchez, A. M., Barra, M. & Rossi, R. H. d. On the Mechanism of the Acid/Base-Catalyzed Thermal *cis−trans* Isomerization of Methyl Orange. *J. Org. Chem.* **64**, 1604–1609, doi:10.1021/jo982069j (1999).

9 Dunn, N. J., Humphries, W. H., Offenbacher, A. R., King, T. L. & Gray, J. A. pH-Dependent *cis-trans* isomerization rates for azobenzene dyes in aqueous solution. *J. Phys. Chem. A* **113**, 13144–13151, doi:10.1021/jp903102u (2009).

10 Sheldon, J. E., Dcona, M. M., Lyons, C. E., Hackett, J. C. & Hartman, M. C. T. Photoswitchable anticancer activity via *trans-cis* isomerization of a combretastatin A-4 analog. *Org. Biomol. Chem.* **14**, 40–49, doi:10.1039/c5ob02005k (2016).

11 Borowiak, M. *et al.* Photoswitchable Inhibitors of Microtubule Dynamics Optically Control Mitosis and Cell Death. *Cell* **162**, 403-411, doi:10.1016/j.cell.2015.06.049 (2015).

12 Boulègue, C., Löweneck, M., Renner, C. & Moroder, L. Redox potential of azobenzene as an amino acid residue in peptides. *ChemBioChem* **8**, 591–594, doi:10.1002/cbic.200600495 (2007).

13 Kulkarni, C. A. & Brookes, P. Cellular Compartmentation and the Redox/Non-Redox Functions of NAD+. *Antioxid Redox Signal*, doi:10.1089/ars.2018.7722 (2019).

14 Zenker, J. *et al.* A microtubule-organizing center directing intracellular transport in the early mouse embryo. *Science* **357**, 925-928, doi:10.1126/science.aam9335 (2017).

15 Singh, A. *et al.* Polarized microtubule dynamics directs cell mechanics and coordinates forces during epithelial morphogenesis. *Nat. Cell Biol.* **20**, 1126-1133, doi:10.1038/s41556-018-0193-1 (2018).

16 Eguchi, K. *et al.* Wild-Type Monomeric α-Synuclein Can Impair Vesicle Endocytosis and Synaptic Fidelity via Tubulin Polymerization at the Calyx of Held. *J. Neurosci.* **37**, 6043–6052, doi:10.1523/jneurosci.0179-17.2017 (2017).

17 Lin, C. M. *et al.* Interactions of tubulin with potent natural and synthetic analogs of the antimitotic agent combretastatin: a structure-activity study. *Molecular Pharmacology* **34**, 200-208 (1988).

18 Kleele, T. *et al.* An assay to image neuronal microtubule dynamics in mice. *Nat. Commun.* **5**, 4827, doi:10.1038/ncomms5827 (2014).

19 Pecqueur, L. *et al.* A designed ankyrin repeat protein selected to bind to tubulin caps the microtubule plus end. *Proceedings of the National Academy of Sciences* **109**, 12011, doi:10.1073/pnas.1204129109 (2012).

20 Kabsch, W. XDS. *Acta crystallographica. Section D, Biological crystallography* **66**, 125-132, doi:10.1107/S0907444909047337 (2010).

21 Weinert, T. *et al.* Serial millisecond crystallography for routine room-temperature structure determination at synchrotrons. *Nature Communications* **8**, 542, doi:10.1038/s41467-017-00630-4 (2017).

22 Adams, P. D. *et al.* PHENIX: a comprehensive Python-based system for macromolecular structure solution. *Acta Crystallographica Section D* **66**, 213-221, doi:doi:10.1107/S0907444909052925 (2010).

23 Emsley, P. & Cowtan, K. Coot: model-building tools for molecular graphics. *Acta Crystallographica Section D* **60**, 2126-2132, doi:doi:10.1107/S0907444904019158 (2004).

24 Steinmetz, M. O. & Prota, A. E. Microtubule-Targeting Agents: Strategies To Hijack the Cytoskeleton. *Trends in Cell Biology* **28**, 776-792, doi:10.1016/j.tcb.2018.05.001 (2018).

25 Ravelli, R. B. G. *et al.* Insight into tubulin regulation from a complex with colchicine and a stathmin-like domain. *Nature* **428**, 198-202, doi:10.1038/nature02393 (2004).
